# Infection-induced vascular inflammation in COVID-19 links focal microglial dysfunction with neuropathologies through IL-1/IL-6-related systemic inflammatory states

**DOI:** 10.1101/2023.06.23.546214

**Authors:** Rebeka Fekete, Alba Simats, Eduárd Bíró, Csaba Cserép, Anett D Schwarcz, Balázs Pósfai, Eszter Szabadits, Zsuzsanna Környei, Krisztina Tóth, Anna Kellermayer, Csaba Dávid, László Acsády, Levente Kontra, Carlos Silvestre-Roig, Judit Moldvay, János Fillinger, Tibor Hortobágyi, Arthur Liesz, Szilvia Benkő, Ádám Dénes

## Abstract

COVID-19 is associated with diverse neurological abnormalities, which predict poor outcome in patients. However, the mechanisms whereby infection-induced inflammation could affect complex neuropathologies in COVID-19 are unclear. We hypothesized that microglia, the resident immune cells of brain, are centrally involved in this process. To study this, we developed an autopsy platform allowing the integration of molecular anatomy-, protein- and mRNA data sets in post-mortem mirror blocks of brain and peripheral organ samples from COVID-19 cases. Nanoscale microscopy, single-cell RNA sequencing and analysis of inflammatory and metabolic signatures revealed distinct mechanisms of microglial dysfunction associated with cerebral SARS-CoV-2 infection. We observed focal loss of microglial P2Y12R at sites of virus-associated vascular inflammation together with dysregulated microglia-vascular-astrocyte interactions, Cx3Cr1-fractalkine axis deficits and mitochondrial failure in severely affected medullary autonomic nuclei and other brain areas. Microglial dysfunction occurs at sites of excessive synapse- and myelin phagocytosis and loss of glutamatergic terminals. While central and systemic viral load is strongly linked in individual patients, the regionally heterogenous microglial reactivity in the brain correlated with the extent of central and systemic inflammation related to IL-1 / IL-6 via virus-sensing pattern recognition receptors (PRRs) and inflammasome activation pathways. Thus, SARS-CoV-2-induced central and systemic inflammation might lead to a primarily glio-vascular failure in the brain, which could be a common contributor to diverse COVID-19-related neuropathologies.

## Introduction

Neurological impairments associated with COVID-19 markedly contribute to disease severity and outcome after SARS-CoV-2 infection (*1*). However, the mechanisms underlying neurological symptoms that are highly heterogenous in appearance, duration and severity in individual patients are not well understood. Clinical imaging and neuropathological studies have revealed diverse pathologies in multiple brain areas including focal hyperintensities, loss of vascular integrity, microcoagulations, gliosis, demyelination, neuronal- and glial injury as well as cell death (*2–7*). Of note, neurological changes such as reduced brain size and grey matter thickness may persist for months after the onset of SARS-CoV-2 infection (*8*). In line with this, symptoms including mental health conditions such as anxiety, depression, cognitive dysfunction, „brain fog” and autonomic / neuroendocrine dysfunction associated with respiratory- and cardiovascular complications after recovery from acute disease and in long-COVID may be present in patients even after apparently mild infection (*2, 9, 10*).

COVID-19 is also associated with broad inflammatory changes, which could contribute to disease pathophysiology via promoting procoagulant states, vascular dysfunction, hypoperfusion, edema, blood brain barrier (BBB) injury or altered glial responses among others (*4, 10–12*). As a hallmark of central pathologies, microglia, the main inflammatory cells in the CNS parenchyma that are highly sensitive to tissue disturbance show marked morphological transformation, microglial nodules and disease associated inflammatory fingerprints in COVID-19 brains (*13–19*). However, mechanistic links between SARS-CoV-2 infection, CNS inflammation and microglial phenotype changes are vaguely established at present (*20, 21*). While microglia may respond to SARS-CoV-2 directly, and the virus may persist in the brain for months (*22, 23*), controversial data exist concerning the extent of productive SARS-CoV-2 infection in the brain parenchyma and its possible links with vascular- and glial states. In particular, how infection and related inflammatory mechanisms could drive microglial phenotype changes and in turn, how altered microglial function could impact on neuropathologies that are found regionally heterogenous across the brain, are not known.

COVID-19 is a multiorgan disease associated with systemic inflammation and shares several common risk factors (old age, chronic vascular disease, metabolic dysfunction, etc.) with those of cardio- and cerebrovascular diseases, where links between systemic inflammation and central vascular- and microglial pathologies have been established (*24, 25*). Thus, we hypothesized that microglial dysfunction due to virus exposure, impaired vascular function a nd/or systemic inflammatory mediators may occur at sites of focal neuropathologies in the affected brain areas. To investigate this, we developed an autopsy platform allowing direct spatial correlation of focal neuropathological- and microglial phenotype changes with levels of inflammatory mediators and viral load in the brain and peripheral organs. We find that focal microglial pathologies affecting different cellular pathways and intercellular communication occur at sites of SARS-CoV-2 associated vascular inflammation, which are strongly linked with an IL-1 / IL-6-related multiorgan proinflammatory response and characteristic virus-sensing pattern recognition receptor signatures, marking sites of diverse neuropathological changes across the brain.

## Results

### COVID-19 is associated with heterogenous microglial pathologies and focal downregulation of microglial P2Y12R driven by viral load

To investigate associations between microglial phenotypes, neuropathological- and inflammatory changes, we developed an autopsy platform allowing correlative analysis of molecular anatomy with tissue proteomic and transcriptomic features after short post-mortem time. We collected CSF samples and mirror tissue blocks from several brain areas and peripheral organs of 11 post-mortem COVID-19 cases either immersion fixed or rapidly frozen on dry ice (Fig. S1). Fixed and frozen brain tissues were also collected from 2 further COVID cases and 9 control cases without neurological disease for comparative studies (Tables S1-S2).

We found major microglial morphological changes with large region-dependent heterogeneities in all examined brain areas of COVID-19 cases in comparison to control. In the bulbus olfactorius, Iba1-positive microglial cuffs and cells with abnormal morphology were associated with sites of disintegrated neuronal cell bodies and nerve fibers (Fig.1a). Iba1 cells showed amoeboid morphology in the choroid plexus and in cranial nerves (n.IX and X assessed), while diverse morphological states ranging from enlarged microglial processes to complete loss of processes and cell cuffs were found in the medulla, hypothalamus and cerebral cortex (Fig.1a). To quantify microglial changes, four brain areas were selected for automated morphometric analysis: the gyrus rectus, the temporal cortex, the hypothalamus and the medulla. The gyrus rectus was chosen due to the proximity of the olfactory bulb, to assess any possible impact of contact viral infection on microglial changes. Unbiased morphology analysis revealed dramatic morphological heterogeneity based on microglial cell volume, sphericity and number of process endings (terminal branches) in the medulla, while all sites contained several cells with marked morphological transformation (Fig.1b-e) with white matter tracts and central autonomic nuclei broadly affected (Fig.S2a). Next, we tested microglial levels of the core purinergic receptor P2Y12R, a key regulator of microglial responses to infection and injury (*26*). We observed a marked downregulation of P2Y12R expression on microglia in focal lesions present in all examined brain areas of COVID-19 cases, including different areas of the cerebral cortex, the thalamus (Fig.S2b) and the hindbrain, with most extensive loss seen in the medulla at sites of central autonomic nuclei (Fig.1f-g). Affected sites of the gyrus rectus and the medulla both showed significantly lower P2Y12R levels compared to control cases (Fig.1h) with profound reduction in the medulla (*p*<0.0001). Of note, the 11 COVID-19 cases examined showed remarkable regional heterogeneities among different brain sites concerning microglial P2Y12R loss (Fig.1i-j). While most of microglia with low P2Y12R showed viable morphology across the brain, in severely affected sites of the medulla, hypothalamus or cerebral cortex we also observed signs of microglial degeneration that was confirmed by electronmicroscopy (Fig.S2c). Surprisingly, we found that P2Y12R downregulation occurred proportionally with increased viral load as measured with qPCR across the brain (r=-0.41, *p*=0.025). Of note, SARS-CoV-2 RNA levels particularly strongly correlated with the loss of P2Y12R mRNA in the medulla (r=-0.87, *p*=0.0025), in line with the most severe microglial pathologies observed among the brain areas examined (Fig.1k), suggesting that viral load may directly or indirectly influence microglial states. SARS-CoV-2 RNA levels were recovered from all brain areas and showed a positive correlation with IFNα mRNA levels (but not with IFNβ) in both the medulla and the gyrus rectus (Fig.1l and Fig.S3a,b), indicating the development of an anti-viral immune response with type I interferon signatures (*12*). Remarkably, the gyrus rectus showed particularly high levels of IFN mRNAs, even compared to peripheral tissues (Fig.S3c).

**Figure 1.**
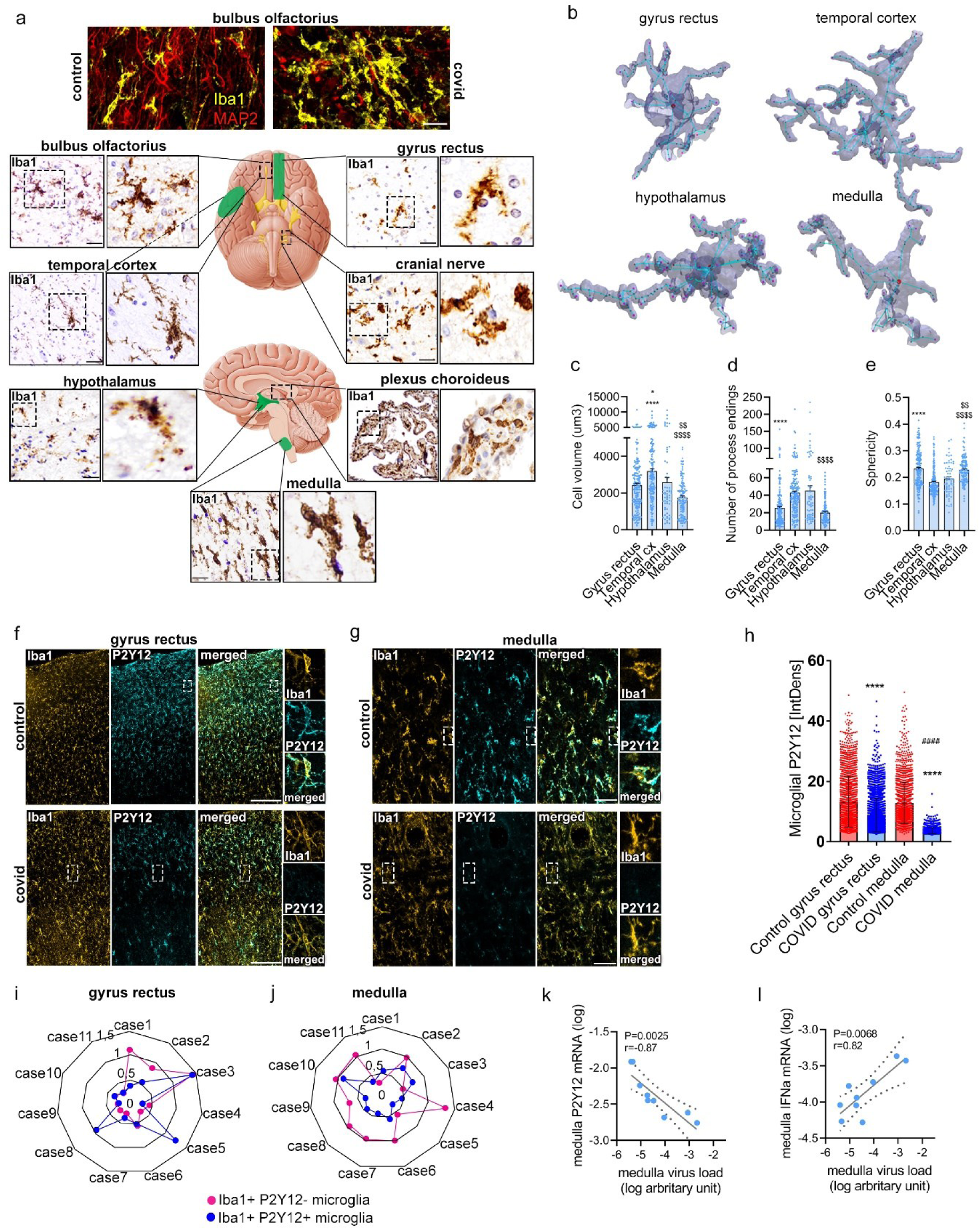
COVID-19 induces microglial reactivity and focal downregulation of microglial P2Y12R proportionally with viral load. **a**. Fluorescent images show microglial morphology changes (Iba1, yellow) associated with marked loss of neurons and nerve fibers (MAP2, red) in a COVID-19 case compared to a non-COVID case in the bulbus olfactorius. Iba1 (DAB immunoperoxidase) labeling shows marked morphological transformation of microglia at multiple brain areas in COVID-19 brains. **b**. Representative 3D surfaces and skeletons showing typical microglial morphologies from unbiased automated analysis. **c-d-e**. Automated morphology analysis reveals major brain region-related differences in microglial phenotypes with most severe changes seen in the medulla. One-way ANOVA with Tukey’s post-hoc test (Cell Volume: \**p*<0.05 vs hypothalamus, \*\*\*\**p*<0.0001 vs gyrus rectus and medulla, ^$$$$^*p*<0.0001 vs gyrus rectus and temporal cx, ^$$^*p*<0.01 vs hypothalamus; Ending nodes: \*\*\*\**p*<0.0001 and ^$$$$^ *p*<0.0001 vs temporal cx and hypothalamus; Sphericity: **** *p*<0.0001 vs temporal cx and hypothalamus, ^$$$$^P<0.0001 vs temporal cx, ^$$^*p*<0.01 vs hypothalamus). **f-g-h.**. Confocal fluorescent images show downregulation of microglial P2Y12R in the affected areas of the gyrus rectus and the medulla (yellow: Iba1, cyan: P2Y12). \*\*\*\**p*<0.0001, COVID vs Control in both brain areas and ^####^*p*<0.0001, COVID gyrus rectus vs COVID medulla, two-way ANOVA followed by Tukey’s post hoc comparison. **i-j.**. Spider charts display heterogeneity of microglial P2Y12R loss in the gyrus rectus and medulla of 11 COVID-19 cases. Charts show an arbitrary heterogeneity score based on the range and average of microglial density values of ROIs measured in given brain areas (0: completely homogenous distribution of microglia). Blue dots represent normal, Iba1+P2Y12+ microglia, pink dots represent Iba1+ P2Y12 microglia. Note that the severity of microglial distribution abnormalities and P2Y12R loss show high variability among COVID-19 cases and microglial pathologies in given patients are also heterogenous concerning the brain area affected. **k**. Viral load in the medulla shows a strong negative correlation with P2Y12R mRNA levels. **l**. Virus load shows a positive correlation with IFNα mRNA levels in the medulla (individual samples where SARS CoV-2 mRNA levels were detectable were included). Correlation graphs show logged data from qPCR with P values and Pearson r correlation coefficient displayed. Scale bars: a. 50 µm, 25 µm; f, 100 µm, g, 50 µm.

### Loss of microglial P2Y12R marks sites of vascular inflammation and viral antigen accumulation in vascular structures and recruited immune cells

We next studied whether focal accumulation of viral infected cells or viral RNA in the brain tissue could explain the heterogenous microglial phenotypes and P2Y12R loss observed across the brain. To this end, two different antibodies to detect SARS-CoV-2 S1 protein and nucleocapsid with immunohistochemistry were tested in paraffin embedded lung tissues (Fig.S3d), and the nucleocapsid (nc) antibody was selected for further studies. In the bulbus olfactorius, both parenchymal profiles and immune cells showed immunopositivity to SARS-CoV-2. Of note, proximity of microglia to virus containing cells and accumulation of viral antigens were observed in a set of microglia with abnormal morphology (Fig.S3e). Nc staining was found in cranial nerves (n.X and n.IX) with immune cells also containing viral antigens (Fig.S3f). We found that several brain areas showed SARS-CoV-2 nc-positive profiles intravascularly or associated with blood vessels with no evidence of widespread, productive infection in neurons, astrocytes or other parenchymal cells. In COVID-19 cases showing the highest viral RNA levels, the medulla was particularly affected, and viral-loaded cells were observed inside blood vessels, in endothelial cells, and in the perivascular space, less frequently in the vicinity of neurons and other parenchymal cells (Fig.2a, b). Here, disintegration of perivascular structures and the presence of viral antigens beyond the vascular endothelium were observed even in larger blood vessels (Fig.2c). Immunoelectron microscopy showed that microglial cell body and processes come into physical interactions with the vascular endothelium at sites showing disintegrated basement membranes and enlarged perivascular space (Fig.2d). Microglia at these sites contained mitochondria with abnormal morphology (Fig.2e). Quantification showed that vessel-associated microglia have lower P2Y12R levels (Fig.2f). P2Y12R accumulates on microglial processes at specific vascular contact sites through which microglia protect against vascular injury, modulate vasodilation and aid adaptation to hypoperfusion (*27, 28*). STED superresolution microscopy showed that while in control cases microglial P2Y12R was found enriched at both perivascular AQP4-positive astrocyte endfeet and at direct endothelial contact sites, microglial P2Y12R around inflamed blood vessels was lost in the affected areas of the medulla in COVID-19 cases (Fig.2g). Loss of P2Y12R from vessel-associated microglia and microglial processes was also observed in the gyrus rectus (Fig.S3g). Immunofluorescent labelling also identified CD45 (common leukocyte antigen)-positive, viral nc-containing leukocytes approached by perivascular microglial processes or internalized by microglia. Perivascular CD45-positive leukocytes without intracellular viral antigens were also observed in the vicinity of SARS-CoV-2–immunopositive cells (Fig.2h). Intra- and perivascular leukocytes, endothelial- and perivascular cells containing virus antigens were found in the gyrus rectus and other cortical areas too, some internalized by microglia (Fig.S3h), indicating that microglial loss of P2Y12R in association with viral load (Fig.1k) primarily takes place at vascular compartments.

**Figure 2.**
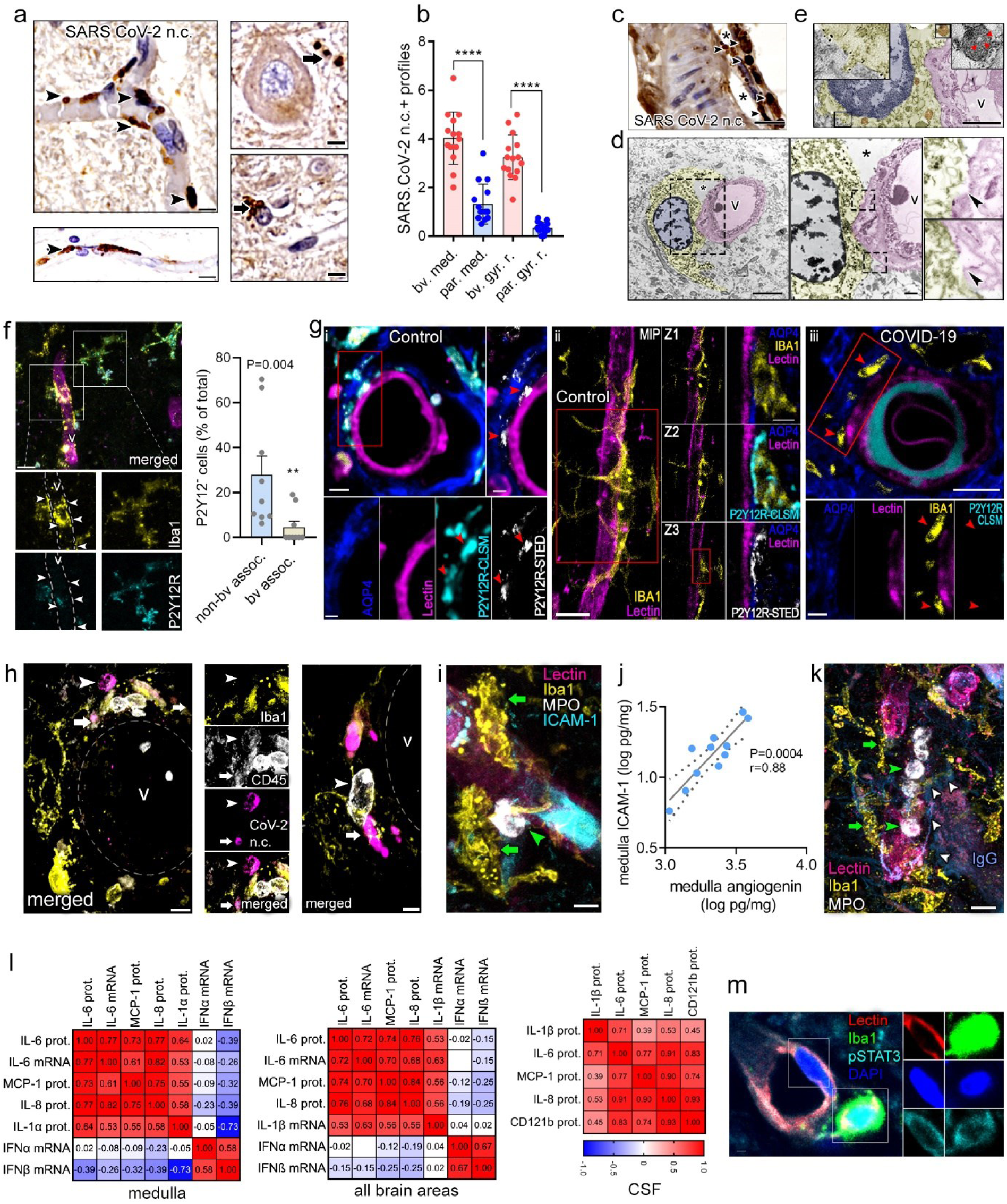
Vessel-associated accumulation of viral antigens in the medulla marks sites of microglial P2Y12R loss, vascular pathologies and an IL-1/IL-6-related inflammatory response. **a**. SARS CoV-2 nucleocapsid staining reveals intra- and perivascular profiles (arrowheads) in the medulla. Note that both clogged blood vessels (top left) and morphologically intact blood vessels (bottom left) contain immunopositive profiles (arrowheads). Viral antigens also appear in the vicinity of neurons and other cells outside blood vessels (right panels, arrows). **b**. Quantification of SARS CoV-2 nucleocapsid immunopositive profiles showing predominant vascular / perivascular association (bv.) as compared with parenchymal sites (par.) in the medulla (med.) and gyrus rectus (gyr. r.). **c**. Disintegration of perivascular structures (*) and the presence of viral antigens (arrowheads) are observed beyond the vascular endothelium in larger blood vessels of the medulla. **d**. Immunoelectron microscopy reveals that microglial cell body and processes form physical interactions with blood vessels (arrowheads) showing disintegrated basement membranes and enlarged perivascular space (*). **e**. Blood vessel-associated microglia in the medulla containing low P2Y12R levels (immunogold labelling, left insert) contain mitochondria with abnormal morphology (right insert). **f**. Microglia (Iba1, yellow) associated with inflamed blood vessels (v, lectin, magenta) show P2Y12R downregulation (cyan, arrowheads) in the medulla. Graph showing comparison of microglia with low P2Y12R levels associated with blood vessels versus non-associated cells (Mann-Whitney test). **g**. STED superresolution microscopy reveals microglial P2Y12R enriched at both perivascular AQP4-positive astrocyte endfeet and at direct endothelial contact sites in the medulla of control cases (i, ii), while microglial P2Y12R around inflamed blood vessels is lost in COVID-19 brains (iii, red arrowheads). **h**. Left panel: perivascular CD45-positive immune cells (arrowheads) containing SARS CoV-2 nucleocapsid are contacted and internalized by Iba-positive microglia (arrows). Right: CD45-positive immune cells (arrowhead) without viral antigens are also present perivascularly in the vicinity of SARS CoV-2 nucleocapsid immunopositive profiles engulfed by microglia (arrow). v – blood vessel. **i**. Iba1-positive microglia (yellow, green arrowheads) are recruited to an ICAM-1 positive blood vessel (cyan) with associated MPO-positive leukocyte (white). **j**. Graphs showing correlation between ICAM-1 vs angiogenin measured by cytometric bead array in medulla tissue homogenates. Logged data with *p* values and Pearson r correlation coefficient displayed. **k**. Iba1-positive microglia (yellow, green arrows) showing marked morphological transformation are recruited to blood vessels (lectin, pink) with intraluminal MPO-positive cells (white, green arrowheads). Plasma leakage through disrupted blood vessels into the brain parenchyma is indicated by the presence of IgG (blue, white arrows). **l**. Pearson correlation matrix showing inflammatory mediators in medulla tissue homogenates (left), in all four brain areas examined (gyrus rectus, temporal cortex, hypothalamus, medulla, middle) and in the CSF. Pearson r correlation coefficient values are displayed. **m**. pSTAT3 immunofluoresence (cyan) in medullary endothelium (lectin, red) and perivascular microglia (Iba1, green). Scale bars: a,f,h,i,k: 10µm; c: 20 µm d: 5 µm, 1 µm; e:5 µm; g: 2 µm, 5 µm; m: 1 µm.

### COVID-19 is associated with a generalized inflammatory response in the brain related to interleukin-1 (IL-1), IL-6 and microglial dysfunction at sites of vascular pathologies

We found the development of vascular inflammation in the medulla at sites of microglial pathologies, as suggested by increased ICAM-1 expression, a vascular adhesion molecule mediating leukocyte adhesion and transmigration (*29*). In line with this, medulla tissue homogenates contained higher ICAM-1 protein levels than other brain areas, while a strong correlation between ICAM-1 and angiogenin (a marker of vascular inflammation and remodelling (*30*)) was observed in individual COVID-19 cases as measured by cytometric bead array (CBA) (Fig.2j). In line with this, we found intra- and perivascular accumulation of leukocytes containing myeloperoxidase (MPO), azurophilic granules of neutrophils and monocytes in ICAM-1 positive blood vessels (Fig.S5 and Fig.2i). A set of MPO-positive cells contained matrix metalloprotease-9 (MMP-9), and leakage of immunoglobulin G (IgG) into the brain parenchyma indicating blood-brain barrier (BBB) injury was also documented with recruited microglia showing marked morphological transformation in the medulla and in the hypothalamus (Fig.2k and Fig.S4a, b) and to a lesser degree in cortical areas (Fig.S4c, d). Of note, previous studies have demonstrated the pathogenic role of neutrophils, monocytes and leukocyte-derived MMP-9 in BBB leakage and breakdown of vascular basement membranes in diverse CNS inflammatory conditions (*31, 32*).

To understand the nature of the inflammatory response associated with microglial dysfunction and vascular inflammation, levels of multiple inflammatory mediators were screened by multiplex CBA in parallel tissue blocks to those analysed with high-resolution anatomy in the selected four brain areas and in the CSF. We found that both isoforms of interleukin-1 (IL-1), a potent driver of vascular inflammation that promotes both ICAM-1, MMP-9 and MPO expression (*33*) were produced in COVID-19 brains. Microglial IL-1α was associated with MPO-positive cells and perivascular microglial pathologies (Fig.S4e) and IL-1α protein levels were significantly increased in the medulla compared to other brain areas with low levels measured in the CSF (Fig.S5a). Although low IL-1β protein levels were recovered from the brain tissue, IL-1β-positive microglia and IL-1β mRNA were detectable at all brain sites and protein levels were elevated in the CSF in half of COVID-19 patients, similarly to CD121b (IL-1R2) (Fig.S4f and Fig.S5a, b).

Further characterization of inflammatory mediators suggested the development of a generalized, but regionally heterogenous inflammatory response in COVID-19 brains related to the observed vascular- and microglial pathologies. Macrophage chemotactic protein-1 (MCP-1) tissue levels showed a strong positive correlation with IL-6 and IL-8 levels not only in the medulla, but within each brain area examined as well as in the CSF (Fig.2l), two proinflammatory cytokines induced by IL-1 that mediate the recruitment of neutrophils, monocytes and macrophages (*32–34*). Positive correlation between mediators and IL-1β mRNA or IL-1β protein was confirmed in tissue homogenates and in the CSF, respectively. Associations of IL-6 protein and mRNA levels indicated that at least in part, IL-6 is produced by cells that reside in the brain with IL-6 signaling suggested by pSTAT3 immunostaining (the main transcription factor downstream to IL-6 receptor activation) in endothelial cells and perivascular microglia (Fig.2l,m). Type I IFN mRNA responses showed weak inverse relationship with IL-1/IL-6/MCP-1/IL-8 with significant inverse correlation between IFNβ and medulla IL-1α (Fig.2l). Thus, this data suggest that COVID-19 is associated with an IL-1 / IL-6-related proinflammatory cytokine- and chemokine milieu with inflammatory cell recruitment and accumulation of viral antigens, marking sites of microglial P2Y12R loss and vascular inflammation alongside with BBB injury in key central autonomic nuclei and other brain areas.

### Microglial dysfunction and viral load associate with central and systemic inflammation in COVID-19

IL-6 and MCP-1 are known to be produced by microglia and modulate microglial activity and migration, while IL-6 has been identified as a key marker of COVID-19-related illness, neurological dysfunction and mortality (*35*). In line with the downregulation of microglial P2Y12R, we observed microglial dislocation in the severely affected brain areas, particularly in the medulla. We quantified this based on a newly developed score of microglial distribution heterogeneity (MDH), which measures spatial heterogeneity of the cells within given fileds of view (Fig.3a). MDH values were markedly higher in the medulla compared to that seen in the gyrus rectus, indicating a more heterogeneous distribution of microglia (Fig.3b). Importantly, we found a strong positive correlation between medulla IL-6, CSF IL-6 / IL-1β levels and medullary MDH scores (Fig.3c). CSF IL-6 also correlated with IL-6 concentrations in all brain areas. Surprisingly, we found that systemic IL-6 levels, particularly in the spleen, showed a strong positive correlation with medullary IL-6 levels in individual patients (*p*=0.00014; r=0.92), while lung IL-1β correlated with CSF IL-1β (*p*=0.035; r=0.64) and spleen IL-1β levels (Fig.3c). Likewise, MCP-1 levels in peripheral organs (particularly in the spleen) strongly predicted IL-6 and MCP-1 levels in every brain area studied (Fig.S6, Table S3), indicating that IL-1 and IL-6-related systemic inflammation may be a considerable contributor to COVID-19-related CNS pathologies.

**Figure 3.**
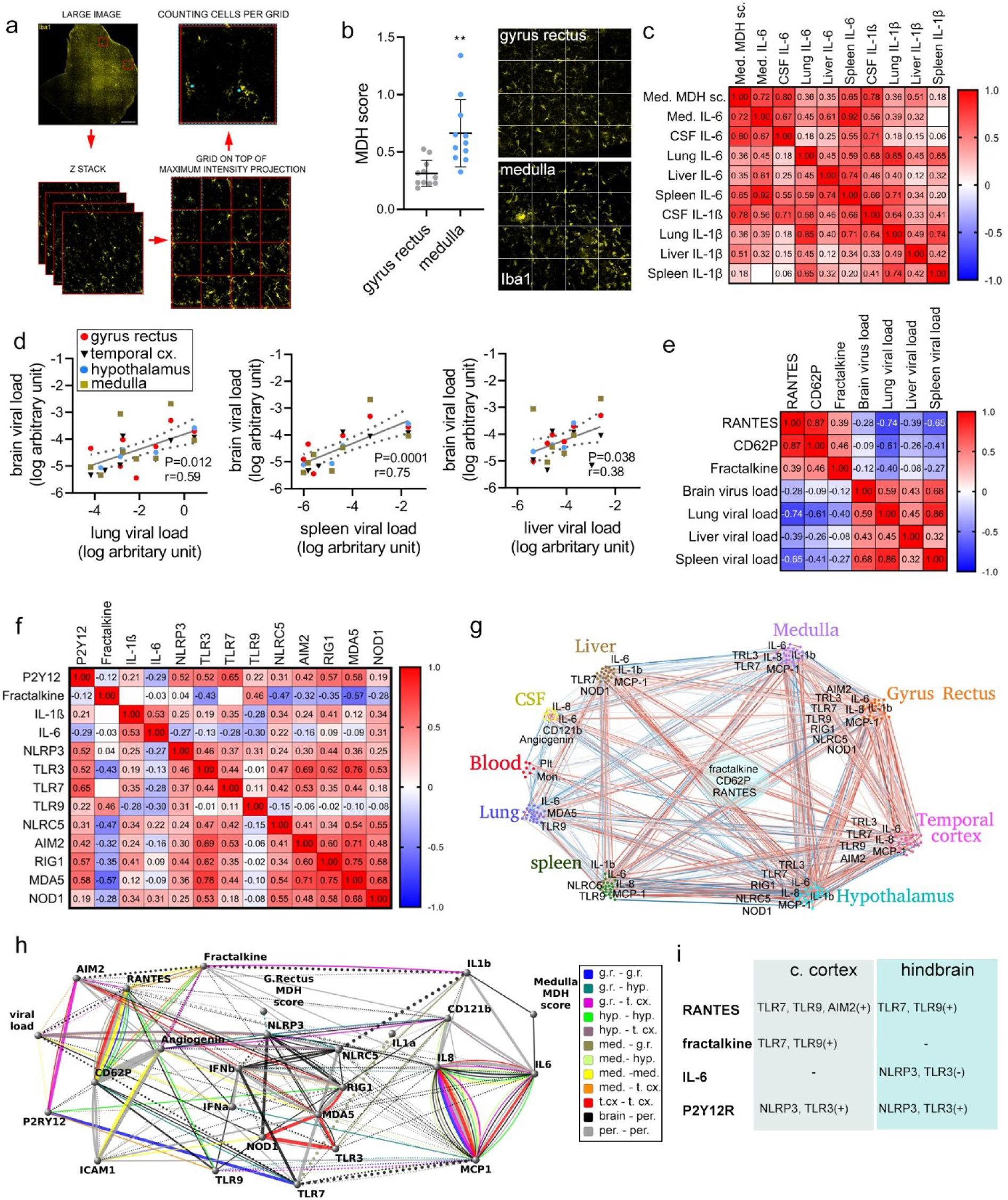
Pattern recognition receptors link microglial states, viral load and inflammation in the brain and peripheral tissues. **a**. Microglial distribution heterogeneity (MDH) score showing the deviation of microglial numbers measured in given quadrants within a brain area from the average microglial numbers in that area based on the distribution of Iba1 immunofluorescenct profiles. High MDH score indicates high level of distribution heterogeneity of (i.e. marked dislocation of parenchymal microglia) compared to normal tissues. **b**. MDH scores in the medulla were significantly higher compared to the gyrus rectus in the COVID-19 cases examined (unpaired t test, \*\**p*<0.01). **c**. MDH in the medulla correlates positively with medullary IL-6 levels, CSF IL-6 and IL-1β levels and IL-6 levels in peripheral organs, particularly in the spleen. Pearson correlation matrix with Pearson r correlation coefficient values are displayed. **d**. SARS-CoV-2 RNA levels measured by qPCR in peripheral organs show positive correlation with SARS-CoV-2 RNA levels in the brain. Note that only cases where viral RNA could be detected in the brain were plotted. **e.** Pearson correlation matrix with Pearson r correlation coefficient values displayed showing positive correlation between RANTES, CD62P and fractalkine levels in the brain tissue (medulla, hypothalamus, gyrus rectus and temporal cortex). Note negative correlation with viral load in peripheral organs. **f**. Pearson correlation matrix with Pearson r correlation coefficient values displayed showing positive correlation between P2Y12R mRNA, IL-1β mRNA and fractalkine protein levels and mRNA levels of several viral-sensing and inflammasome-forming PRRs. **g**. Network representation of motif patterns in COVID-19 cases. Line width represent p-values with emphases of connections with p-values less than 0.0025. Blue line shows negative, red line positive correlation. Correlation with a p-value lower than 0.01 has been highlighted and labeled. **h**. Network showing key sites of interconnectedness of the assessed markers among brain areas and peripheral tissues. The width of the connecting line corresponds to p-values. Dotted line is negative correlation, continuous line is positive correlation. Colored lines show connections within the brain. **i**. Key PRRs of the interaction network in the cerebral cortex (gyrus rectus + temporal cortex) and the hindbrain (medulla + hypothalamus). Note overlapping and different PRRs vs markers that strongly influence microglial states. Scale bar: a: 1000 µm.

Because SARS-CoV-2 infection may persist in peripheral organs and in the brain for months (*22*), we next wondered to what extent the development of multiorgan infection may explain associations between systemic and central inflammatory states. Of note, viral RNA levels in the brain were comparable to that seen in the liver and the spleen, while levels in the lung were 2-3 orders of magnitude higher (Fig.S3a). Surprisingly, we found that viral load in peripheral organs, particularly in the spleen (*p*=0.0001; r=0.75), the lung (*p*=0.0012; r=0.59) and the liver (*p*=0.038; r=0.38) predict central SARS-CoV-2 mRNA levels (Fig. 3d). In particular, lung viral load was strongly associated with viral load in the hypothalamus (*p*=0.025, r=0.92). While viral load did not show strong associations with the IL-1/IL-6/MCP-1/IL-8 axis, we found that high viral mRNA levels in the lung and the spleen predicted low RANTES (CCL5), CD62P (P-selectin) and fractalkine levels in the brain, mediators that have been associated with COVID-19-related disease severity, vascular inflammation, platelet activation and microglial function (*36–38*). While we found a strong positive correlation between angiogenin, RANTES, CD62P and ICAM-1 in all peripheral organs examined, central RANTES also showed a strong, positive correlation with CD62P levels in all four brain areas examined (Fig.3e; Fig.S6, Table S3). Importantly, levels of fractalkine, a key chemokine mediating microglia-neuron and microglia vascular interactions via its receptor, CX3CR1, correlated with RANTES and CD62P levels (*p*<0.0001, r=0.78) and was reduced in the hindbrain compared to cortical areas (Fig.S6a), suggesting that increasing systemic viral burden might compromise microglial inflammatory responses via the CX3CR1-fractalkine axis at these sites (*39, 40*).

### Pattern recognition receptors link inflammation and microglial dysfunction with marked brain region-related differences, bridging central and systemic inflammatory states

To further understand the mechanisms underlying neuropathological- and inflammatory changes in the brain, we assessed several pattern recognition receptors (PRRs) that sense SARS-CoV-2 and/or cellular dysfunction to modulate central inflammatory actions. Specifically, IL-1-related inflammatory signatures in the COVID-19 brain suggested the role of inflammasomes, which regulate IL-1β processing and release in response to pathogens, cellular stress or tissue injury (*41*). In line with the severity of microglial pathologies, we found that the medulla showed elevated levels of RIG1, MDA5, TLR3, TLR7 and NLRC5, while the hypothalamus showed increased RIG1 and MDA5 levels compared to the gyrus rectus and the temporal cortex with no difference observed in AIM2, NOD1 and TLR9 levels (Fig.S6, Fig.S7, Table S3). NLRC5, RIG1, MDA5, NOD1 are cytosolic sensors for viral RNA and associated inflammation, while RNA-sensing TLR3, TLR7 and TLR9 are expressed both intracellularly and on the cell membrane (*42*). Interestingly, we found that increased proinflammatory load (i.e. elevated IL-1/IL-6/IL-8/MCP-1 levels) and microglial dysfunction (downregulation of P2Y12R and increased MDH score) correlated with lower levels of viral mRNA sensors in the brain, with substantial regional heterogeneities concerning given PRRs. Of note, the PRRs AIM2, TLR3, TLR7, NLRC5, RIG1, MDA5 and NOD1 showed a cross-correlation throughout the brain and together with NLRP3 and TLR9 were differentially associated with IL-1β mRNA, P2Y12R and fractalkine levels, suggesting the involvement of different, overlapping PRR pathways in modulating central inflammatory states (Fig.3f).

To understand how anti-viral immunity and related inflammatory changes could contribute to CNS pathologies, we next looked at associations of PRRs with inflammatory mediators in peripheral organs and the brain. Bioinformatic analysis showed that brain areas showed entrenched connections especially between the temporal cortex and gyrus rectus with TLR7, TLR9 and AIM2 reoccurring with the highest correlation scores, while TLR3 and TLR7 were most representative for the hindbrain strongly associated with IL-1β/IL-6/IL-8/MCP-1 in the brain tissue and with CSF IL-6, IL-8 angiogenin and CD121b levels (Fig.3g). In peripheral organs, two main clusters of genes showed the same pattern in all patients: IL-6/IL-8/MCP1 and RANTES/CD62P/Angiogenin/ICAM1 with fractalkine and TLR7 linking between these clusters. The lung, the primary site of SARS-CoV-2 infection showed high levels of proinflammatory mediators including IL-1β, IL-6, MCP-1 and IL-8, which were positively associated with NLRP3 and IL-1β mRNA levels and negatively with anti-viral type I IFNs with MDA5 in the lung tissue, showing a strong negative correlation with brain IL-6, MCP-1 and IL-8 levels (Fig.S6). High viral load and low TLR9 mRNA in both the spleen and the lung were associated with lower central RANTES, CD62P and fractalkine levels (Fig3g,h, FigS6, Table S3). Interestingly, while C-reactive protein (CRP) measured in the blood also showed a positive association with CSF fractalkine levels (*p*<0.0029; r=0.64), D-dimer, a major biomarker in COVID-19-related mortality and procoagulant states (*4*) showed positive correlation with lung IL-1β mRNA, NLRP3 mRNA and IL-6 protein levels. Furthermore, lower platelet-, and lymphocyte counts as well as low hematocrit predicted high viral load, high proinflammatory burden and P2Y12R downregulation in the brain with similar positive association between monocyte counts and P2Y12R levels (Fig.3g-h). Finally, similarly to related inflammatory mediators, viral sensors and inflammasome-forming PRRs showed a strong association with microglial states (i.e. MDH scores, P2Y12R and fractalkine levels). However, considerable differences were seen between the hindbrain, where microglial and vascular pathologies were more severe than in the cerebral cortex, in line with augmented proinflammatory states (Fig.3g-i, Fig.S6, Table S3).

### Single nuclear RNA sequencing reveals microglial dysfunction, mitochondrial failure and disrupted cell-cell interactions in COVID-19

To further investigate how microglial dysfunction may contribute to inflammatory changes in COVID-19 brains, we performed single nuclei mRNA sequencing from medulla samples of COVID-19 and control patients. We projected a total of 16,260 nuclei and identified 9 different cell populations based on the most variable genes (Fig.4a, Fig.S8). Differential gene expression analysis on the microglial cell cluster between COVID-19 and control samples identified several functionally relevant candidates, including a reduction in P2Y12R and fractalkine expression, corresponding to the observations shown above (Fig. 4b, c). As previous microanatomical studies suggested changes of cellular interaction in COVID-19, we performed a cell-cell communication network analysis using CellChat (*43*). Comparing the total number of interactions and the interaction strength across cell types between conditions, we found that COVID-19 resulted in an overall reduction of the cell-cell communication pathways across cell populations in the brain (Fig. 4d, e). Microglial interaction with neurons, oligodendrocytes and OPCs was substantially reduced in COVID-19, but stronger interaction with T cells and astrocytes was detected, mainly through the osteopontin (OPN/SPP1)-CD44 axis (Fig. 4c, e,-f). To confirm our findings on the impairment of the microglial CX3CR1-fractalkine (CX3CL1) signaling axis in patients with COVID-19, we further studied the cell-cell communication of this particular network. We found that the signal between neurons (sender) and microglia (receiver) via the CX3CL1-CX3CR1 axis prevailed in control samples, but was completely absent in COVID-19 samples (Fig.4g). Since microglial cell function is largely influenced by its metabolic state (*44*), we further studied metabolic pathways in microglia and all other major cell populations identified by scSeq between COVID-19 and control cases (Fig.4h). Interestingly, while diverse metabolic pathways were abundantly dysregulated in COVID-19, neurons and T cells were barely affected; in contrast microglia and other glial cells showed highly significant differences across multiple metabolic pathways. Specifically, metabolic differences in microglia of COVID-19 cases indicated towards changes in mitochondrial function (TCA cycle, electron transport chain, etc.). Therefore, we focused on differential expression of (nuclear transcribed) mitochondrial proteins and identified a significant down-regulation of many mitochondrial genes, which are known to code for a very diverse set of proteins encompassing the different protein complexes from the mitochondrial electron transport chain (Fig.4i, j). These results potentially indicate mitochondrial failure in microglia due to COVID-19. As for the metabolic dysregulation, this pronounced down regulation of mitochondrial genes was also observed in oligodendrocytes and astrocytes from COVID conditions, but minimally in endothelial cells, T cells and neurons, overall suggesting glial-specific mitochondria impairment in COVID-19 (Fig.4j). Histological analysis confirmed marked morphological transformation of mitochondrial networks in COVID microglia, some immunopositive to BAX indicating induction of apoptotic cascades and abnormal ultrastructure of mitochondrial membranes, especially in cells associated with inflamed blood vessels (Fig.4k-m).

**Figure 4.**
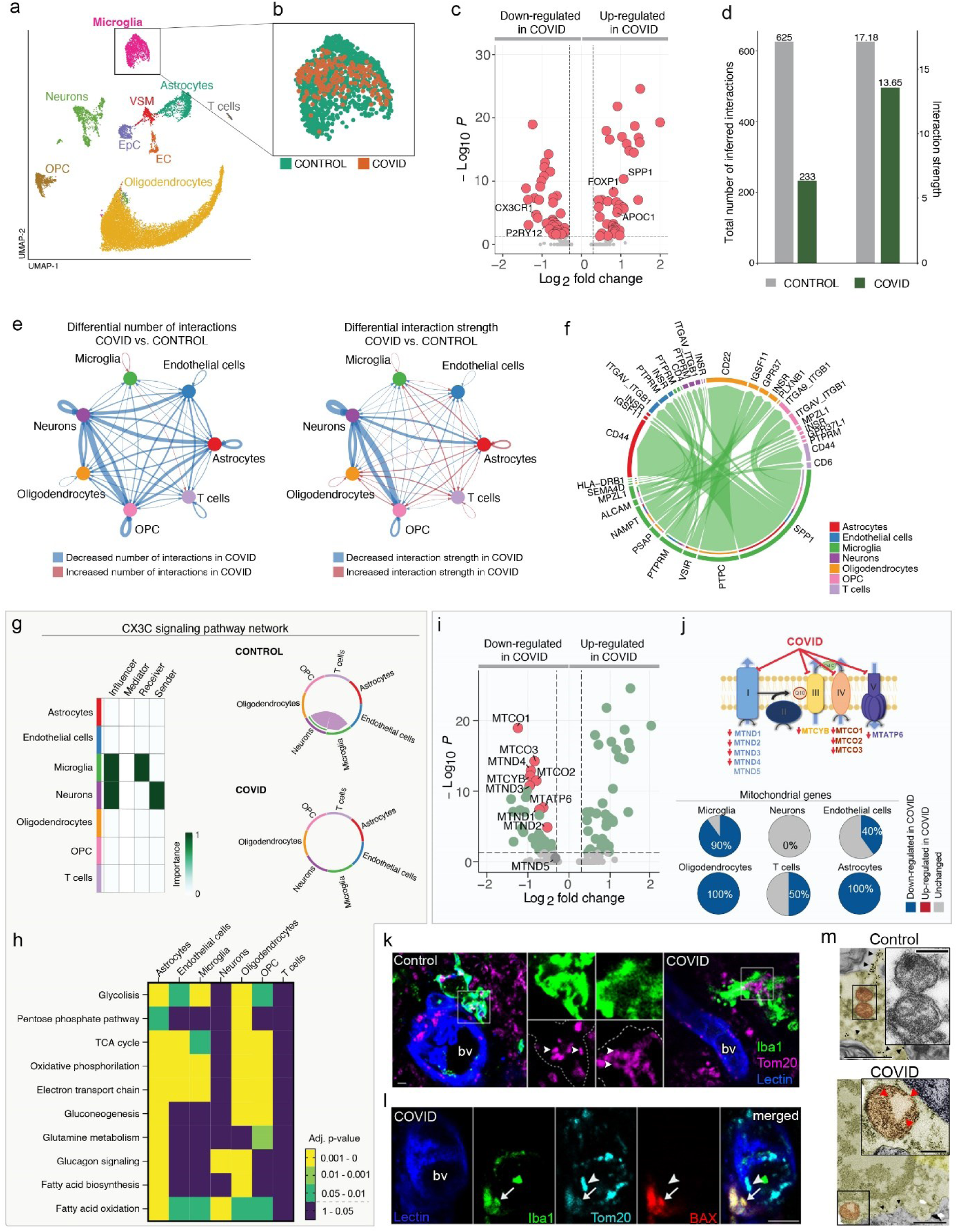
Single cell RNA seq reveals microglial dysfunction, mitochondrial failure and disrupted cell-cell interactions in COVID-19. **a**. Uniform Manifold Approximation and Projection (UMAP) plot of a total of 16,260 brain nuclei analyzed using single-nuclei mRNA sequencing (10x). Nuclei are colored by identified cell populations. **b**. UMAP plot of microglia (1,345 nuclei), colored by condition. **c**. Volcano plot showing the up- and down-regulated genes in microglia between control and COVID samples. Colored genes are *p*<0.05 and |fold-change|>1.25. **d**. Bar plot (right panel) showing the total number of interactions (left) and interaction strength (right) of the inferred cell-cell communication networks from control (grey) and COVID (dark green) conditions. **e**. Circle plots (left panels) showing the differential number of interactions (left) and differential interaction strength (right) among cell populations in the cell-cell communication network between control and COVID samples. Red and blue colored edges represent increased or decreased signaling, respectively, in the COVID compared to controls conditions. **f**. Chord diagram showing all significant interactions (L-R pairs) from microglia to all other cell populations. **g**. Heatmap showing the identification of dominant senders, receivers, mediators and influencers in the CX3C communication network (left panel). The relative importance of each cell group is based on the four computed network centrality measures of CX3C signaling network. Chord diagrams (right panels) showing significant interactions (L-R pairs) among all cell populations for the CX3C signaling network in control (left) and COVID (right) samples. **h.** Heatmap showing the adjusted p-values of the differential enrichment of metabolic signatures between control and COVID samples per each cell type. **i.** Volcano plot (left panel) showing the significantly down-regulated mitochondrial genes (red, *n*=9) in microglia from COVID patients compared to controls. Colored genes are *p*<0.05 and |fold-change|>1.25. **j.** Schematic representation of the mitochondrial electron transport chain (upper panel), showing the 5 protein complexes and the identified down-regulated mitochondrial genes in COVID microglia per complex.. Pie charts showing the percentage of differentially expressed mitochondrial genes between control and COVID conditions per cell population (lower panel). Percentages are calculated based on total number of identified mitochondrial genes per cell population. **k**. Microglial mitochondria (Tom20, magenta) in the medulla of COVID-19 patients show marked morphological changes compared to that seen in control brains (arrowheads). **l**. Morphologically normal mitochondria (arrowheads) lack BAX as opposed to BAX-positive microglial mitochondria in the medulla (arrow). **m**. Immunoelectron microscopy shows mitochondria with abnormal morphology in COVID-19 medulla tissues. Scale bars: k: 5μm, l: 10μm, m:. 1 μm, inserts: 250 nm.

### Microglial dysfunction and excessive phagocytic activity parallel synapse loss and myelin injury in COVID-19

Finally, we aimed to understand the associations between microglial dysfunction and neuropathological changes in the most affected medullary sites, where inflammatory mediator levels, histology and single nuclei mRNA sequencing data suggested major changes in microglia and their interactions with vascular structures and neurons. Of note, severely affected sites of the medulla overlapped with key central autonomic centers around the fourth ventricle, including the dorsal vagal nucleus, hypoglossal nucleus, the solitary nucleus, the vestibular nuclei, and more laterally / ventrally areas of the inferior cerebellar peduncle, the raphe nucleus, the nucleus ambiguus and ventral respiratory nuclei. Affected sites also included the medial lemniscus, the pyramidal tract and the olivary nuclei with large regional heterogeneities in given patients. We observed increased numbers of phagocytic microglia expressing CD68 (a phagolysosme marker) at these sites that engulfed glutamatergic synapses as suggested by internalized vGluT1/Homer immunopositive profiles (Fig.5a-c). Phagocytosis by microglia expressing P2Y12R was broadly observed throughout the brain, suggesting that parenchymal microglia-related pathologies may not be explained merely by loss of microglial P2Y12R. However, we also observed loss of P2Y12R from microglial processes contacting synapses and neuronal cell bodies at sites where severe pathologies and loss of P2Y12R from vascular-associated microglia were seen by using STED superresolution microscopy (Fig.5d,e), which is likely to suggest impaired microglia-neuron interactions, phagocytic responses and deficient recognition of viable and/or damaged neurons (*45*). Tissue pathology at microglia-synapse contacts was also confirmed at ultrastructural level (Fig.5f). To quantify synapse loss, we turned to a sensitive post-embedding approach (*46*) that had been optimized in the laboratory for human post-mortem tissues (Fig.5g). Because for technical reasons during tissue collection not all hypothalamic samples were suitable for high resolution anatomy, we performed analysis in the medulla, gyrus rectus and temporal cortex tissue sites showing „average” microglial pathology or „severe pathology” based on microglial P2Y12R loss and MDH scores. Post-embedding immunolabelling of vGluT1 and synapsin-positive profiles showed synapse loss in both average and severe COVID-19 pathology compared to non-COVID brains in all brain areas examined (Fig.5h-i). We also observed microglial engulfment of degenerating cells and neuronal cell bodies, which was more severe in the medulla compared to the gyrus rectus in given COVID-19 cases (Fig5j-k). We also found substantial myelin degeneration and myelin loss at sites showing marked microglial pathologies with frequent associations of microglia with disintegrated myelin sheath and myelin engulfment by microglia that was confirmed by electron- and confocal micrsocopy (Fig.5l). Thus, vascular-driven microglial dysfunction and associated inflammatory changes are likely to interfere with severe neuropathologies in COVID-19.

**Figure 5.**
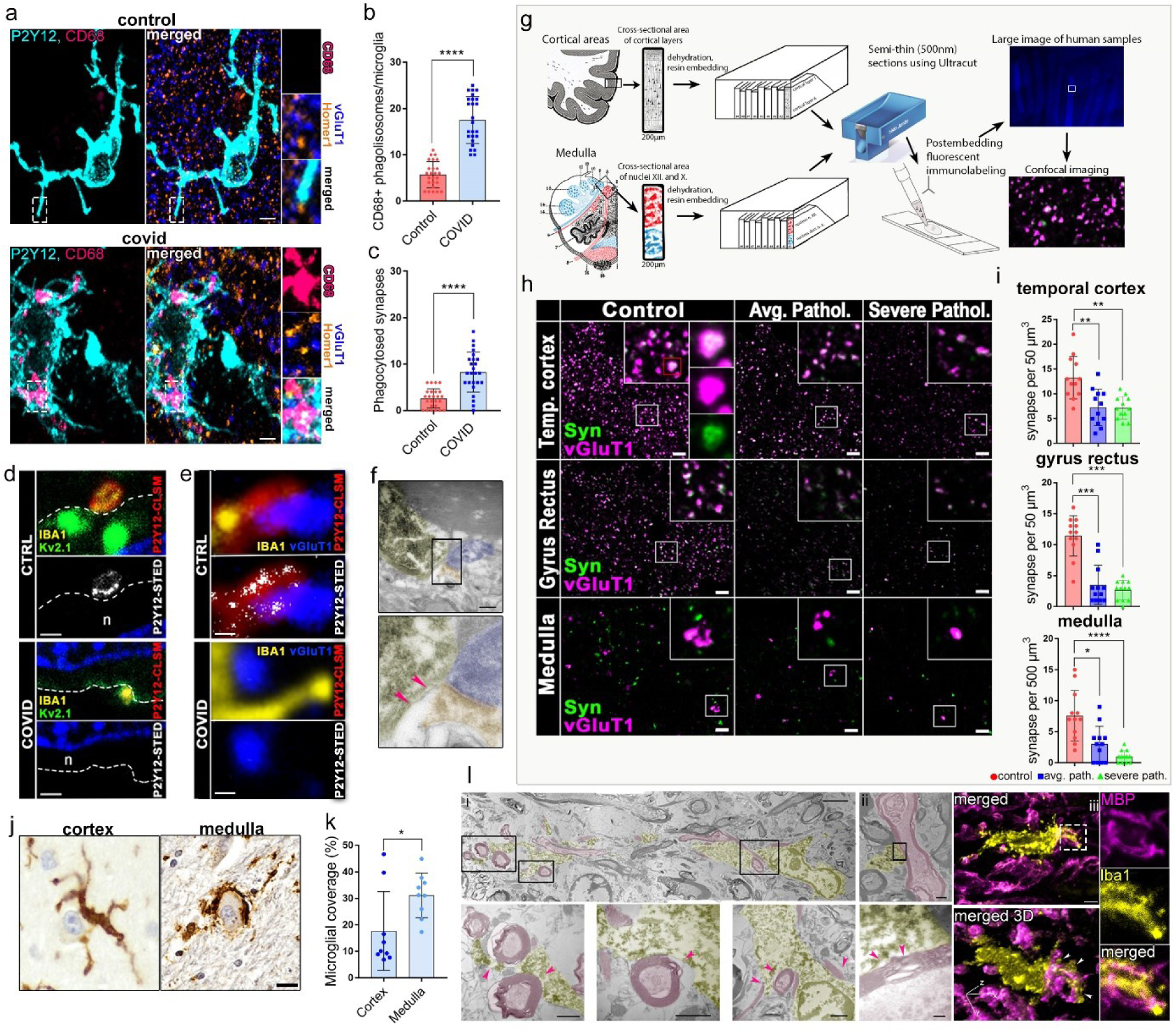
COVID-19 is associated with synapse loss and microglial phagocytosis of myelin and synapses. **a**. Fluorescent images show P2Y12 positive (cyan) CD68 negative microglia (pink) contacting vGluT1 (blue) Homer1 (orange) synapses in a non-COVID brain (top panel), while in COVID-19 brains P2Y12, CD68 positive microglia internalize vGluT1 / Homer1 synapses (arrowhead shows on bottom panel). **b**-**c**. Graphs showing significant increase in microglial CD68-positive phagolysosome numbers and increased synaptic phagocytosis by microglia in COVID-19 cerebral cortex. **d**. STED superresolution microscopy shows microglial P2Y12 receptors enriched at Kv2.1 positive neuronal somatic contact site (somatic junction) in the gyrus rectus of non-COVID cases (top panel), while in COVID-19 brains P2Y12R expression is downregulated at somatic contact sites (bottom panel, n: neuron). **e**. STED superresolution microscopy reveals microglial P2Y12R-positive microglial process contact at vGluT1 positive synaptic terminals in control brains (top lanel) while in COVID-19 brains P2Y12 receptors are lost at these synaptic contacts in severely affected areas of the gyrus rectus. **f**. TEM image showing Iba1 DAB immunoperoxidase-positive microglia process contacting a synapse in COVID-19 brain. Pink arrowheads show tissue loss around the contact site. **g**. 200 µm thick sections were cut from the gyrus rectus, temporal cortex and medulla, then sections were dehydrated and embedded into resin. One resin block contained samples from all patients from one brain region. Then ultracut sectioning was done on the cross-sectional surface of the sections and mounted onto glass slides. Fluorescent immunolabeling was performed on slides followed by light and confocal laser scanning microscopy. **h.** Confocal fluorescent panels show post-embedding immunolabelling of vGluT1 (pink) and Synapsin-positive (green) synapses for quantitative assessment. Note the loss of synapses in both average and severe COVID-19 pathology compared to control brains **i.** Quantification of post embedding immunolabelling reveals significant synapse loss in COVID-19 temporal cortex, gyrus rectus and medulla compared to the same brain areas in non-COVID cases. **j.** Iba1 DAB immunoperoxidase labelling with cresyl violet counterstain shows microglial engulfment of degenerating neuronal cell bodies in the cerebral cortex and medulla of COVID-19 cases. **k.** Average microglial somatic coverage of degenerating neuronal cell bodies is significantly increased in the medulla compared to the gyrus rectus in the same COVID-19 cases. **l**. TEM (left panels) and fluorescent confocal pictures (right panel) showing frequent associations of microglia with disintegrated myelin sheath, while myelin engulfment by microglia is also observed in COVID-19 medulla samples. Pink (TEM panel) and white (confocal 3D panel) arrowheads show sites of microglia-myelin interactions and phagocytosis. Scale bars : a : 5 µm, d, e : 500 nm, f. h. 5 µm, j. 5 µm, l. (i) 3 µm, 1 µm (ii) 1 µm, 250 nm (iii) 1 µm.

## Discussion

The mechanisms through which central and systemic effects of SARS-CoV-2 infection contribute to diverse neurological abnormalities in functionally and anatomically distinct sites of the CNS have remained enigmatic to date, despite intense research efforts due to the enormous health-related and socioeconomical burden of COVID-19. By using a unique autopsy platform and multimodal characterisation of inflammatory states, we show that in individual COVID-19 cases, viral load and the associated proinflammatory response in peripheral organs are strongly linked with viral load and inflammation, characterized by IL-1-and IL-6-related vascular pathologies and specific virus-sensing PRR fingerprints in different brain areas. Dysfunction of microglia, the main immune cells of the CNS parenchyma, characterized by their morphological transformation, spatial dislocation, loss of P2Y12R, impaired CX3CR1-fractalkine communication and mitochondrial failure occurs at sites of viral antigen-associated vascular injury, marking areas of microglia-related synapse- and myelin injury. These pathologies are more severe in the hindbrain than in the cerebral cortex, particularly in the medulla, where key central autonomic centers are localized.

The mechanisms whereby SARS-CoV-2 could drive inflammation and microglial changes in the brain have remained unclear to date. While 10 of the 11 systematically examined cases were unvaccinated allowing us to reliably characterize COVID-19 -related inflammatory states, it is generally difficult to control for the broad effects of critical illness and intensive care (e.g. mechanical ventillation, medication, etc) on the CNS. Thus, in addition to non-COVID cases, we compared different brain areas and peripheral organs within the COVID cohort for key readouts, to ensure appropriate assessment of viral load, inflammation and microglial phenotype changes across the brain. Beyond the olfactory bulb and cranial nerves containing high SARS-CoV-2 nc levels, viral proteins in the brain were mostly localized to blood vessels and not found inside neurons. This argues against a primarily neurotropic spread of SARS-CoV-2, in line with most neuropathological findings (*47*). Instead, viral antigens in vascular structures and intra- or perivascular immune cells were found near microglial processes or internalized by microglia. While microglia may not directly restrict SARS-CoV-2 replication (*48*), they are equipped with an arsenal of PRRs that recognize viral antigens inducing inflammatory signaling and cell death to clear infected cells (*42, 49*). In fact, while viral RNA levels in the brain were proportional to anti-viral type I IFN responses (a characteristic signature against RNA virus infection), viral load related microglial P2Y12R downregulation, microglial dyslocation and cell loss were most apparent in the severely affected medulla. Here, vascular inflammation could be triggered by IL-1, IL-6 and other mediators, while IL-8 and MCP-1 could provide a chemokine milieu for the recruitment of neutrophils and monocytes containing MMP-9 and MPO, to promote vascular pathologies and BBB injury. Microglial phenotypes are strongly modulated by systemic inflammation (*50*). To our surprise, despite the relatively small number of cases and their heterogenous clinical background, similar inflammatory fingerprints were identified in each brain area studied with strong links to CSF patterns and systemic inflammatory states. In line with this, systemic viral load was not only linked with brain viral load in individual cases, but negatively correlated with a different set of mediators (RANTES/CD62P/fractalkine) in the brain. These observations may suggest that while chronic inflammatory comorbidities promote the risk of poor clinical outcome in COVID-19 cases (*51*), SARS-CoV-2 infection markedly reshapes proinflammatory fingerprints across the body. Supporting our findings, circulating IL-6, IL-8 and MCP-1 levels are strongly associated with disease severity (*52–54*) in COVID-19 patients, while high viral load and low CCL5 expression levels were associated with intensive care unit admission or death (*36*). However, these inflammatory mediators have only been characterized in the CSF previously in COVID-19 cases and central-systemic links have not been established. Our data also suggest that the proposed links between circulating- and CSF IL-6 levels in COVID-19 patients are not explained by inflammatory mediators reaching the CSF from the systemic circulation (*55*), rather by a brain-body-wide induction of similar proinflammatory signatures as promoted by multiorgan SARS-CoV-2 infection and related vascular pathologies.

The exposure of microglia and vascular structures to viral antigens and the development of an IL-1 related proinflammatory response suggested the involvement of PRRs, including inflammasome forming NLRs that regulate IL-1β production (*41*). Cytosolic virus sensors NOD1, RIG1, MDA5, TLR3, TLR7,TLR9 and NLRC5 recognize SARS-CoV-2 infection among which RIG1 and MDA5 are key drivers of anti-viral type I IFN responses, while production of IL-1β could be induced by activation of TLRs 3-7-9, and the inflammasomes NLRP3 and AIM2, as supported by observations in monocytes- or monocyte-derived macrophages upon exposure to SARS-CoV-2 or viral antigens (*23, 42, 56, 57*). Of note, in the human brain, different PRRs not only showed strong association with microglial states, but correlation among PRRs and characteristic PRR fingerprints were also found in the cerebral cortex and the hindbrain (the latter showing more severe vascular microglia- and neuropathologies overall). For example, upregulation of TLR3 with other PRRs in the severely affected medulla paralleled a negative correlation with proinflammatory mediators and severe vascular / microglial pathologies. This might suggest an insufficient coping with infection, similarly to the lung, where endothelial TLR3 insufficiency in mice contributes to vascular remodeling by SARS-CoV-2 (*58*).

Microglia emerge as key modulators of neuronal- and vascular responses and gatekeepers against central neurotropic virus infection including coronaviruses (*59, 60*). However, the mechanisms behind their marked phenotypic transformation in COVID-19 and its consequences for disease outcome have remained largely unclear to date. Focal loss of P2Y12R across numerous brain areas, a key receptor mediating microglial responses to injury or infection is expected to augment neuronal and vascular pathologies in COVID-19 brains. Importantly, superresolution microscopy showed that P2Y12R loss occurs at key vascular and neuronal contact sites for intercellular communication with microglia. Absence of microglia or microglial P2Y12R leads to neuronal hyperactivity, and impairs hypoperfusion, vascular injury, neuronal death and neurological outcome in experimental models of acute brain injury or neuronal hyperexcitability (*26-28, 61-64*). Correspondingly, meta-analysis of EEG findings has revealed high proportion of abnormal background activity in COVID-19 patients (96%), with epileptiform discharges present in 20% of cases (*65*). Long-term microstructure and cerebral blood flow changes have also been observed in patients recovered from COVID-19, which correlated with C-reactive protein (CRP) and interleukin 6 (IL-6) levels even in the absence of neurological manifestations (*66*). We also observed marked impairments concerning the CX3CR1-fractalkine axis in COVID-19. Deficiencies in microglia-neuro-vascular communication due to impaired CX3CR1-fractalkine interactions are known to influence synapse- and neuron loss as well as proinflammatory respones in rodents (*67*). According to our data, focal microglial pathologies may arise from vessel-associated virus induced inflammation in multiple foci across the brain, where dysrupted vascular basement membranes and BBB injury was also observed. Supporting this, our single cell RNA seq studies revealed diverse metabolic pathways abundantly dysregulated in COVID-19 in microglia, astrocytes and endothelial cells, while neurons and T cells were barely affected. Thus, impaired microglial interaction with neurons, oligodendrocytes and OPCs in part via failures in core microglial regulator proteins (P2Y12R, CX3CR1) could precede the excessive synapse / myelin loss observed. While dysfunctional microglia may directly contribute to synapse- and myelin phagocytosis, diverse pathologies in severely affected brain sites may also involve secondary effects due to microglial death, as it was observed in the medulla. In fact, SARS-CoV-2 promotes microglial synapse elimination in human brain organoids, while functional microglia appear to be important to attenuate demyelination in neurotropic coronavirus infection in mice (*68, 69*).

We suggest that the development of the largely heterogenous neuropathologies seen in multiple foci across the brain in given patients are likely to be strongly influenced by deficient gliovascular communication due to vessel-associated SARS-CoV-2 infection and related microglial dysfunction. These changes are markedly modulated by systemic viral load and inflammatory states. Such inflammatory „hot spots” might render given brain areas alongside with body-wide effects of tissue hypoxia, multiorgan failure and systemic inflammatory burden particularly fragile, augmenting focal microcapillary coagulation, perfusion deficits and vascular- or neuronal injury. In turn, affected medullary- and hypothalamic sites could contribute to impaired neuroendocrine- and autonomic nervous system function, neuropathies or cardiovascular/respiratory abnormalities as parts of a vicious circle, while inflammation at other, functionally distinct areas, such as the thalamus or the cerebral cortex may contribute to sleep disturbances or cognitive deficits. Microglial disfunction may have long-lasting effects in patients who recover after SARS-CoV-2 infection. Changes in microglial activity may last for extended periods (months to years) after various brain injuries. Similarly, in patients with long-COVID syndromes, TSPO PET imaging for microglial reactivity has been shown to be strongly associated with depressive symptoms and cognitive dysfunction (*70*). This suggests that microglial dysfunction and related inflammatory states might be a cornerstone of both acute- and long-COVID neurological mantifestation. Our data therefore argue for the development of therapeutic tools targeting microglial dysfunction to alleviate acute- and long-COVID neuropathologies.

## Supporting information

Supplemental figure 6 (interactive)

## Acknowledgements

We thank the Microscopy Center, the Cell Biology Center and the Human Brain Research Laboratory at the Institute of Experimental Medicine (IEM) for kindly providing support in microscopy, measurement of inflammatory mediators, and for human brain samples, respectively. We are also grateful to Miklós Palkovits for his help in isolating medullary autonomic nuclei.

## Author contributions

Experimental design A.D.; Methodology R.F., A.D.S., E.B., C.C., A.S., B.P., Z.K., E.S., K.T., A.K., C.D.,L.K, T.H., A.L., S.B., A.D.; Formal Analysis R.F., A.S., E.B., C.C., A. D.S., B.P., Z.K., E.S., K.T., A.K., C.D., L.K., A.D.; Investigation R.F., A.S., E.B., C.C., A. D. S., B.P., Z.K., E.S., K.T., A.K., C.D., A.L., L.K., J.M., J.F., T.H., A.L., S.B., A.D.; Resources, A.D., S.B., A.L.; Writing – Original Draft A.D., R.F.; Editing R.F., A.S., E.B., C.C., A.S., B.P., Z.K., E.S., K.T., A.K., C.D., A.L., L.K., J.M., J.F., T.H., A.L., S.B., A.D.; Visualization R.F., A.S., E.B., C.C., A.D.S., B.P., Z.K., E.S., K.T., A.K., C.D., L.K., A.L., S.B., A.D.;. Supervision A.D., S.B., A.L., and A.L.; Project Administration A.D.; Funding Acquisition, A.D., S.B., A.L.

## Ethics statement

Human brain samples were collected in accordance with the Ethical Rules for Using Human Tissues for Medical Research in Hungary (HM 34/1999) and the Code of Ethics of the World Medical Association (Declaration of Helsinki). All procedures were approved by the Regional Committee of Science and Research Ethics of Scientific Council of Health (ETT TUKEB IV/5187 -2/2020/EKU, ETT TUKEB 31443/2011/EKU, renewed: ETT TUKEB 15032/2019/EKU).

## Methods

### Patients and post mortem tissue collection

Thirteen deceased individuals with COVID-19 confirmed by PCR for SARS-CoV-2 and six non-COVID cases were included in the study (Tables S1 and S2). Autopsies were performed at the National Korányi Institute of Pulmonology (n=11 COVID cases) and Szent Borbála Hospital (*n*=2 COVID-19 cases and *n*=6 non-COVID-19 cases) and in the framework of the Human Brain Tissue Bank (HBTB), Budapest, in Hungary (*n*=2 non-COVID cases), under ethical licence ETT TUKEB IV/5178-2/2020/EKU and ETT TUKEB 31443/2011/EKU, renewed: ETT TUKEB 15032/2019/EKU. In the 11 COVID-19 cases, whole-body autopsy was performed in the National Korányi Institute of Pulmonology, which included a thorough tissue collection from different peripheral organs including the lung, the liver and the spleen with tissue blocks collected from several brain areas including the olfactory bulb, cranial nerves, choroid plexus, meninges, gyrus rectus, temporal cortex, frontal gyrus, motor cortex, hippocampus, striatum, occipital lobe (deep white matter), midline thalamus, mesencephalon, pons and medulla. CSF samples were also collected and were frozen immediately. Immediately after removal, all tissue blocks were divided to two parts. One part was frozen on dry ice immediately for molecular biology studies (Fig. S1). A mirror block for each frozen block was immersion fixed on site for 6 hours in Zamboni I fixative (4% paraformaldehyde, 0,05% glutaraldehyde, 15% picric acid), placed on a shaker at room temperature. The fixative was replaced with fresh Zamboni I fixative solution in every 30min and changed to Zamboni II (4% paraformaldehide, 15% picric acid) for overnight at 4 °C. Brains of the two COVID-19 cases and six non-COVID-19 cases were perfused with 4% paraformaldehide at the Szent Borbála Hospital. Control subjects had no history of psychiatric or neurologic deficits, and their deaths were not caused directly by any brain damage. Authopsy patients were unvaccinated against SARS-CoV-2, except one case (Table S1). From all cases, clinical records were assessed thoroughly for progression of COVID-19 related symptoms before death.

### Tissue processing and immunohistochemistry

To investigate SARS-CoV-2 infection, localization of virus antigens and microglia, a set of lung and brain tissues from COVID-19 cases were embedded into paraffin after fixation. 4-6 µm thick FFPE tissue sections were incubated at 45 °C for 2 hours, deparaffinized in xylene, and hydrated in ascending alcohol solutions. Peroxidases were blocked by 1% H2O2 for 15 min at room temperature while gentle agitation was applied. Heat antigene retrieval was carried out by submerging slides in 1x pH 6 citrate buffer for 20 min at 90 °C. Sections were allowed to cool down to 23-25 °C. Then slices were incubated with 1% human serum albumin for 1 hour at room temperature. Slices were then treated with anti-rabbit SARS CoV-2 or anti-guineapig Iba1 antibodies (Table S5) overnight at 4 °C. On subsequent slices negative SARS CoV-2 antibody controls were made following the same protocol. This was followed by incubation in with biotinylated goat anti-rabbit and donkey anti-guineapig antibodies (Table S5) for 4 h at room temperature then in avidin–biotinylated horseradish peroxidase complex (Elite ABC; 1:300; Vector Laboratories) diluted in PBS for 3 h at room temperature. The immunoperoxidase reaction was developed using 3,3-diaminobenzidine (DAB; Sigma-Aldrich) as chromogen. Sections were lightly counterstained with haematoxylin, dehydrated in graded alcohols, cleared in xylene then coverslipped with Depex (Sigma). Images were captured by using 20x and 60x objectives and a Nikon Ni-E C2+ microscope.

### Immunfluorescent labeling and confocal laser scanning microscopy

From immersion fixed- or perfusion-fixed tissue blocks 50 µm thick sections were cut with vibratome. Before immunostaining the sections were washed in 0.1 M PB and 1% Sodium borohydride (Sigma) for 5 minutes. Then sections were washed again in 0.1 MPB and Tris-buffered saline (TBS). This was followed by blocking for 1 hour in 1% human serum albumin (HSA; Sigma-Aldrich) and 0.1% Triton X-100 dissolved in TBS. After this, slices were incubated in mixtures of primary antibodies for 48 hours at 4 °C and washed in TBS, then were incubated in mixtures of secondary antibodies diluted in TBS at 4 C° overnight. Secondary incubation was followed by TBS washes, then sections were mounted on glass slides, and coverslipped with Aqua/Poly/Mount (Polysciences). Immunfluorescence was analyzed using a Nikon Eclipse Ti-E inverted microscope (Nikon Instruments Europe B.V., Amsterdam, The Netherlands), with a CFI Plan Apochromat VC 60X oil immersion objective (numerical aperture: 1.4) and an A1R laser confocal system. Image stacks were obtained with NIS-Elements AR software. For primary and secondary antibodies used in this study, please see Table S5.

### Automated morphological analysis of microglial cells

From gyrus rectus, temporal cortex, hipothalamus and medulla, 100 µm thick sections were cut with vibratome and were immunostained with guineapig anti-Iba1 (1:500 #234308, Synaptic Systems), Alexa 647 donkey anti-guineapig (1:500; #706-606-148, Jackson ImmunoResearch) chicken anti-Iba1 (1:500 #234009, Synaptic Systems), Alexa 647 donkey anti-chicken (1:500; #703-606-155, Jackson ImmunoResearch) antibodies and DAPI. Imaging was carried out in 0.1 M PB, using a Nikon Eclipse Ti-E inverted microscope (Nikon Instruments Europe B. V., Amsterdam, the Netherlands), with a CFI Plan Apochromat VC 60X water immersion objective (numerical aperture: 1.2) and an A1R laser confocal system. For 3D morphological analysis of microglial cells, the open-source MATLAB-based Microglia Morphology Quantification Tool was used (available at https://github.com/isdneuroimaging/mmqt). This method uses microglia and cell nuclei labeling to identify microglial cells. Briefly, 59 possible parameters describing microglial morphology are determined through the following automated steps: identification of microglia (nucleus, soma, branches) and background, creation of 3D skeletons, watershed segmentation, and segregation of individual cells (*71*).

### Microglia distribution analysis

Microglial distribution heterogeneity was measured on maximum intensity projection images of 35-40 µm, 50 µm thick sections, captured at 0.6 µm/px resolution with a Nikon A1R confocal system. The imaged area was divided into 16 equal territories (ROI), and Iba1 expressing cells were counted in each of these ROIs. Heterogeneity score was calculated for each image by dividing the range of measured cell counts (maximum number of cells/ROI - minimum number of cells/ROI) with the average number of cells/ROI. This way a heterogeneity score of 0 means completely homogenous distribution, and larger scores mean larger irregularities in distribution.

### Pre-embedding immunoelectron microscopy

The 0.1 M PB and 0.05 M PBS used for the experiments contained 0.05% sodium azide. After extensive washes in 0.1 M PB and 0.05 M PBS sections were incubated in 1% NA-borohydride for 5 minutes followed by thorough washing with PB and after PBS. Sections were blocked in 1% HSA in PBS (BS) followed by incubation in primary antibodies (Table S5) diluted in PBS, for 2-3 days. After repeated washes in PBS, the sections were incubated in blocking solution (Gel-BS) containing 0.2% cold water fish skin gelatine and 0.5% HSA in PBS for 1 h. Next, sections were incubated in gold-conjugated or biotinylated secondary antibodies (Supplementary Table 5) diluted in Gel-BS overnight. After extensive washes in PBS and PB the sections were treated with 2% glutaraldehyde in PB for 15 min, after which the sections were washed with PB and PBS. This was followed by incubation in avidin–biotinylated horseradish peroxidase complex (Elite ABC; 1:300; Vector Laboratories) diluted in PBS for 3 h at room temperature. The immunoperoxidase reaction was developed using 3,3- diaminobenzidine (DAB; Sigma-Aldrich) as chromogen. To enlarge immunogold particles, sections were incubated in silver enhancement solution (SE-EM; Aurion) for 40-60 min at room temperature. The sections were then treated with 0.5% OsO4 in 0.1 M PB at room temperature, dehydrated in ascending alcohol series and acetonitrile and embedded in Durcupan (ACM; Fluka). During dehydration, the sections were treated with 1% uranyl acetate in 70% ethanol for 20 min. For electron microscopic analysis, tissue samples were glued onto Durcupan blocks. Consecutive 70 nm thick sections were cut using an ultramicrotome (Leica EM UC6) and picked up on Formvar-coated single-slot grids. Ultrathin sections were examined in a Hitachi 7100 electron microscope equipped with a Veleta CCD camera (Olympus Soft Imaging Solutions, Germany).

### Post-embedding immunfluorescent labelling and quantitative analysis

The technique described by Holderith et al. (*46*) was used with slight modifications. 200 µm thick brain slices were washed in 0.1M PB and 0.1M Maleate Buffer (MB, pH: 6.0). Then slices were treated with 1% uranyl-acetate diluted in 0.1M MB for 40 minutes in dark. This was followed by several washes in 0.1M PB, then slices were dehydrated in ascending alcohol series, acetonitrile and finally embedded in Durcupan (Fluca). Each block contained sections of all patients (4 control, 4 covid). Ultrathin sections were cut using a Leica UC7 ultramicrotome at 500 nm thickness, and collected onto Superfrost Ultra plus slides and left on a hotplate at 80°C for 30 minutes then in oven at 80°C overnight. Sections were encircled with silicon polymer (Body Double standard kit, Smooth-On, Inc.) to keep incubating solutions on the slides. The resin was etched with saturated Na-ethanolate for 7 minutes at room temperature. Then sections were rinsed three times with absolute ethanol, followed by 70% ethanol and then DW. Retrieval of the proteins was carried out in 0.02M Tris Base (pH = 9) containing 0.5% sodium dodecyl sulfate (SDS) at 80°C for 80 min. After several washes in TBS containing 0.1% Triton X-100 (TBST, pH = 7.6), sections were blocked in TBST containing 6% BlottoA (Santa Cruz Biotechnology), 10% normal goat serum (NGS, Vector Laboratories) and 1% BSA (Sigma) for 1 hour then incubated in the primary antibodies diluted in blocking solution at room temperature overnight with gentle agitation. After several washes in TBST the secondary Abs were applied in TBST containing 25% of blocking solution for 3 hours. After several washes in TBST, slides were rinsed in DW then sections were mounted in Slowfade Diamond (Invitrogen) and coverslipped. Immunofluorescence was analyzed using a Nikon Eclipse Ti-E inverted microscope (Nikon Instruments Europe B.V., Amsterdam, The Netherlands), with a CFI Plan Apochromat VC 60X oil immersion objective (numerical aperture: 1.4) and an A1R laser confocal system. We used 488 and 647 nm lasers (CVI Melles Griot), and scanning was done in line serial mode, pixel size was 50×50 nm. Image stacks were taken with NIS-Elements AR.

### STED superresolution imaging

50 µm thick, free-floating brain sections were washed in PB and Tris-buffered saline (TBS). This was followed by blocking for 1 hour in 1% human serum albumin (HSA; Sigma-Aldrich) and 0.03-0.1% Triton X-100 dissolved in TBS. After this, sections were incubated in mixtures of primary antibodies overnight at room temperature. After incubation, sections were washed in TBS and were incubated overnight at 4 °C in the mixture of secondary antibodies, all diluted in TBS. For CLSM channels Alexa dye conjugated antibodies were used, for the STED channel we used Abberior Star 635P conjugated antibodies. Secondary antibody incubation was followed by washes in TBS, PB, and sections were mounted in Slowfade Diamond (Invitrogen) and coverslipped. Immunofluorescence was analyzed using an Abberior Instruments Facility Line STED Microscope system built on an Olympus IX83 fully motorized inverted microscope base (Olympus), equipped with a ZDC-830 TrueFocus Z-drift compensator system, an IX3-SSU ultrasonic stage, a QUADScan Beam Scanner scanning head, APD detectors and an UPLXAPO60XO 60X oil immersion objective (numerical aperture: 1.42). We used 405, 488, 561 and 640 nm solid state lasers for imaging, and a 775 nm solid state laser for STED depletion. Image acquisition was performed using the Imspector data acqusition software (version: 16.3.14278-w2129-win64).

### Cytometric bead array (CBA)

CBA measurements were performed on frozen brain tissue homogenates (gyrus rectus, temporal cortex, medulla, hypothalamus), lung, liver, spleen tissue homogenates and CSF samples. Tissue samples were homogenized in TritonX-100 and protease inhibitor-containing (1:100, #539131, Calbiochem) Tris-HCl buffer (TBS, pH 7.4) and centrifuged at 17000 g, for 20 min at 4 °C. Protein level was quantified for every sample using BCA Protein Assay Kit (#23225, ThermoFisher Scientific). Cytokine levels in brain homogenates were normalized for total protein concentrations and expressed as pg/ml values in the CSF. Concentrations of TNFα, IFNγ, IL-17A, IL-6, IL-1α, IL-1β, IL-8, MCP-1, RANTES (CCL5), CD54 (ICAM-1), CD62P (P-selectin), CD121b, Angiogenin and Fractalkine were measured by BD CBA flex sets (BD Biosciences, US) according to the manufacturer’s guidelines. Measurements were performed by using a BD FACSVerse flow cytometer and analyzed by FCAP array software (BD Biosciences, US).

### Sample homogenization and qPCR

100 mg tissue was homogenized in 1 ml TRIzol (Thermo Fisher Scientific, Waltham, MA, USA) using a Beadbug 3 microtube homogenizer (Benchmark Scientific, Sayreville, NJ, USA). Samples were added to a microtube filled with 3-4 pieces of 2.8mm stainless steel beads and were shaken for 120-180 seconds on 300 RPM. Following homogenization, samples were centrifuged for 10 seconds to remove any undissolved particles. Supernatant was stored at -80C° until further use.

#### RNA isolation

Total RNA was isolated from tissue samples using TRIzol reagent following the manufacturer’s protocol. The RNA concentration was measured by a spectrophotometer (NanoDrop ND1000; Promega Biosciences, Madison, WI, USA). RNA integrity was assessed with Agilent 2100 Bioanalyzer (Agilent Technologies, Santa Clara, CA, USA), which evaluate the RNA integrity number (RIN) and the 28S/18S (rRNA) ratio. Samples were stored at -80C° in nuclease free water.

#### Determination of viral load

The viral load in different tissue samples was determined as described previously (*72*) TaqMan™ 2019nCoV Assay Kit v1 (Thermo Fisher Scientific, Waltham, MA, USA).

#### RT-PCR and Quantitative Real-Time PCR

Same amount of RNA was treated with DNase and RNAse inhibitor (Ambion, Austin, TX, USA), then subjected to reverse-transcription using SuperScript II first strand reverse transcriptase and oligo dT primers (Thermo Fisher Scientific, Waltham, MA, USA).

Gene expression was measured by PCR using a QuantStudio12K Flex qPCR instrument (Applied Biosystems, Foster City, CA, USA). The program setting consisted of a 95 °C 10 min denaturation step, and 40 two-stage PCR cycles (95 °C for 12 s and 60 °C for 1 min). Taqman Gene Expression Assays were used with the Taqman™ Gene Expression Master Mix (Applied Biosystems, Foster City, CA, USA). Comparative Ct method was used to calculate the relative gene expression of each target normalized to the ACTB gene.

|  |  |
| --- | --- |
| ACTB | Hs01060665_g1 |
| IFNA1 | Hs00256882_s1 |
| IFNB1 | Hs01077958_s1 |
| IL-1B | Hs01555410_m1 |
| IL-6 | Hs00174131_m1 |
| P2YR12 | Hs00375457_m1 |
| TLR3 | Hs01551078_m1 |
| TLR7 | Hs00152971_m1 |
| TLR9 | Hs00152973_m1 |
| NLRP2 | Hs01546932_m1 |
| NLRP3 | Hs00918082_m1 |
| NOD1 | Hs00196075_m1 |
| NLRC5 | Hs00260008_m1 |
| RIG-I (Ddx58) | Hs00204833_m1 |
| MDA5 (Ifih1) | Hs00223420_m1 |
| AIM2 | Hs00915710_m1 |

### Single Nuclei mRNA –seq

Single nuclei suspensions were prepared from flash-frozen human brain tissues using the Chromium Nuclei Isolation Kit with RNase Inhibitor (PN-1000494, 10x Genomics). Single nuclei suspensions were then submitted to a microscopic inspection of integrity and further processed according to manufacturer’s (10x) protocols at a concentration of 4.000-8.000 nuclei/µl. Single nuclei mRNA libraries were prepared using the 10x Chromium Next GEM Single Cell 3’ v.3.1. Solution. Quality control of all cDNA samples was performed with a Bioanalyzer 2100 (Agilent Technologies) and libraries were quantified with the Qubit dsDNA HS kit (ThermoFisher). Libraries were sequenced on an Illumina NextSeq 2000, aiming for 25,000 reads per nuclei.

### Single Nuclei mRNA-seq data processing and analysis

Cell Ranger software was used to process raw data, align reads to the human hs38 reference genome and summarize unique molecular identifier (UMI) counts. Filtered gene-barcode matrices containing only barcodes with UMI counts that passed the threshold for nuclei detection were used for further analysis. Filtered UMI count matrices were processed using R and the R package Seurat. As quality control steps, the following nuclei were filtered out for further analysis: (1) nuclei with a number of detected genes <200 or >4000, and (2) nucleiwith >3% of counts that belonged to mitochondrial genes. Raw gene counts in high-quality singlets were log normalized and submitted to the identification of high variable genes by MeanVarPlot method. Data was scaled and regressed against the number of UMIs and mitochondrial RNA content per cell. Data was subjected to principal component analysis and unsupervised clustering by the Louvain clustering method. Clusters were visualized using Uniform Manifold Approximation and Projection (UMAP) representations. Clusters were manually annotated using the top upregulated genes for each cluster and the expression of key previously described markers. Differentially expressed genes between conditions were calculated using the FindMarkers function. Volcano plots were generated using EnhancedVolcano in R (Bioconductor EnhancedVolcano v.1.6.0). Cell-to-cell communication network analysis was performed using R toolkit CellChat. In brief, probable interactions at the cell-to-cell level were calculated via computeCommunProb function using the 10% truncated mean method to compute average gene expression (parameter trim = 0.1). Communication probabilities were then calculated by invoking the computeCommunProbPathway function. Ligand receptor interaction probabilities were visualized via netVisual functions. Measures in weighted directed networks, including out-degree, in-degree, flow betweenesss and information centrality, were used to respectively identify dominant senders, receivers, mediators and influencers for the intercellular communications, followed by the netAnalysis_signalingRole_network for visualization. CellChat objects were compared via compareInteractions function. A gene signature enrichment analysis using the ‘AUCell’ method (*73*) was also applied to identify cells with active metabolic signatures. In brief, gene signatures representing different metabolic pathways were manually annotated based on datasets from Gene Ontology and KEGG pathways resources (glycolisis (mmu70171 + GO 0006096), pentose phosphate pathway (GO 0006098), TCA cycle (GO 0006099), oxidative phosphorylation (GO 000619), electron transport chain (GO 0022900), gluconeogenesis (GO:0006094), glutamine metabolism (GO 0006541), glucagon signaling (GO 0071377), fatty acid biosynthesis (mmu00061 + GO 0006633), fatty acid oxidation (GO 0019395)). Mouse gene IDs were converted to Human gene IDs using the g:Convert function from g:Profiler (*74*). The area under the curve (AUC) was then calculated to determine the proportion of genes from each metabolic gene signature that were highly expressed in each nucleus. Statistical differences of the resulting AUC values between nuclei from control and COVID samples were calculated using Student’s t-test with Benjamini-Hochberg correction.

### Correlation and network analysis

Protein, qPCR and complete blood count measurements were Z-score normalized. Pearson correlation in all possible combination was calculated, p-values were calculated and corrected by Benjamini-Hochberg method. Cystoscape was used to visualize *p*<0.001 connections.

### Quantification and statistical analysis

All statistical analysis was performed using Graphpad Prism 7 software (GraphPad Software, Inc). Based on the type and distribution of data we applied appropriate statistical tests: in case of two independent groups of data unpaired t-test or Mann Whitney U-test, for multiple comparisons one-way ANOVA with Kruskal-Wallis test was used. Correlation matrix was based on Pearson’s correlation test. Data were presented as means ± SEM.

### Data availability

Raw mRNA sequencing data generated in this study is available in the NCBI Gene Expression Omnibus (GEO) database repository (GSE234720). The used scripts for bioinformatic analyses of sequencing data are also available at https://github.com/Lieszlab.

## Supplementary Figures

**Figure S1.**
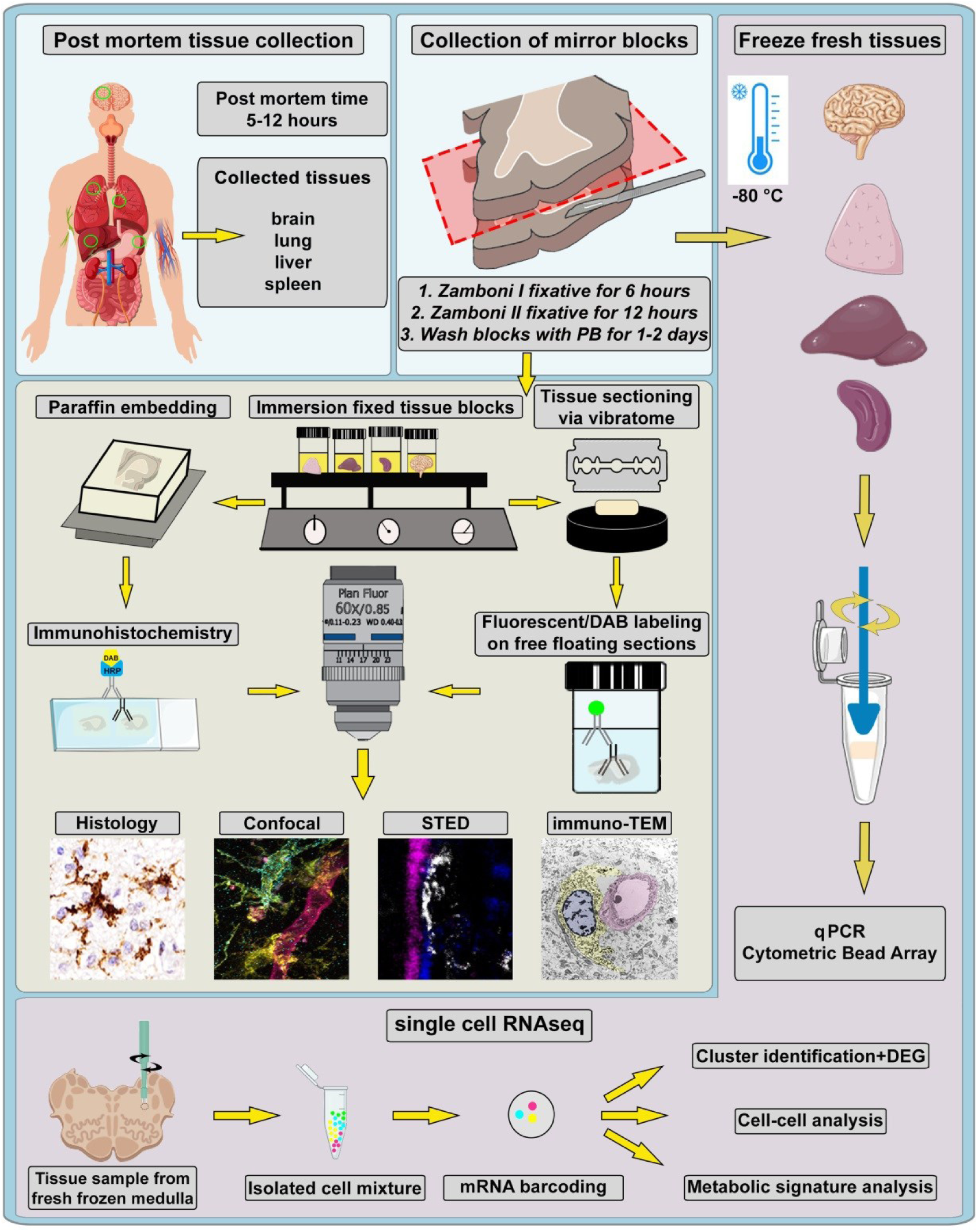
Post mortem tissue collection platform for correlated studies of inflammatory processes in COVID-19. Tissue samples were collected from several brain areas and peripheral organs (lung, liver and spleen tissues analysed) of 11 COVID-19 cases with post-mortem CSF samples from all patients. Autopsies were performed 5-12 hours after death. After removal, all tissue blocks were immediately dissected into two parts, one frozen on dry ice for molecular biology studies and the other immersion fixed with Zamboni fixative. Through the whole fixation process tissue blocks were kept on a shaker and the fixative solution was changed in every hour for 6 hours at room temperature. Then, tissue blocks were postfixed for 12 hours, washed with 0.1M PB for 1-2 days and were cut with vibratome or embedded into paraffin for histological assessment. Free floating sections were examined by immunfluorescent labeling and confocal fluorescent laser scanning/superresolution microscopy and EM. FFPE sections were labelled with DAB immunoperoxidase. Sets of frozen tissue samples were processed for qPCR, cytometric bead array and single nuclear RNAseq studies.

**Figure S2.**
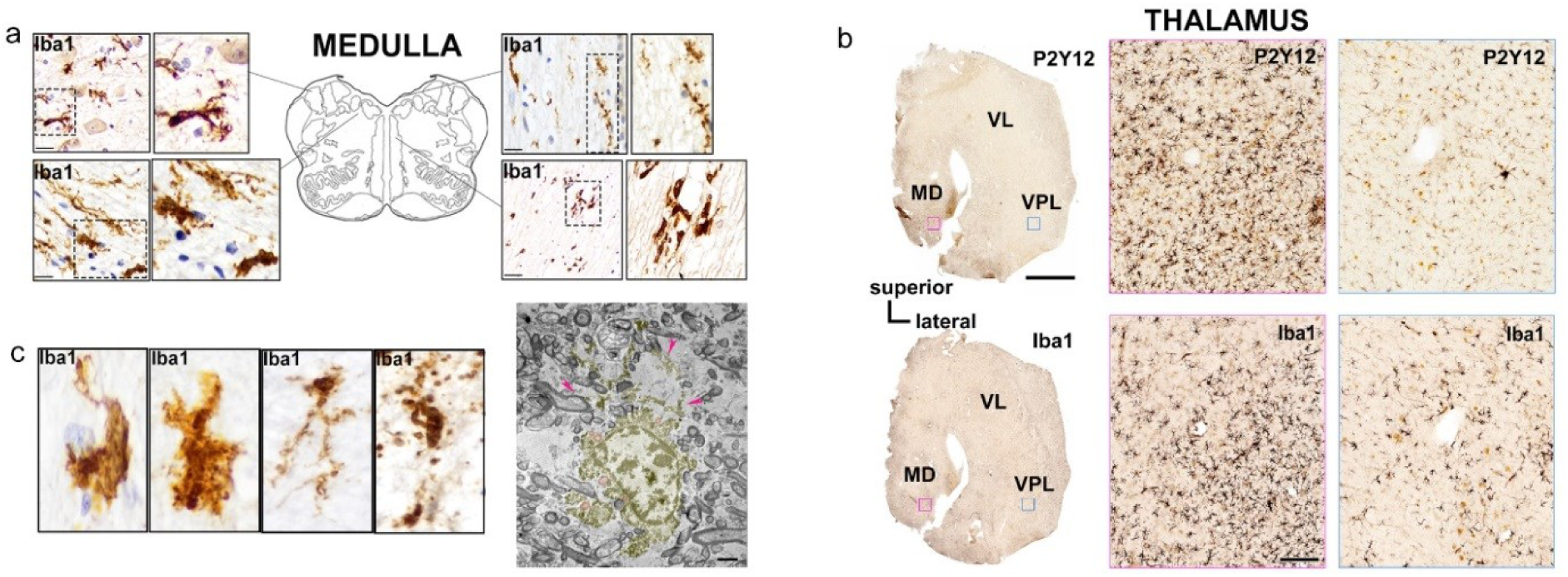
Microglial phenotype transformation and degenerative changes in the brain. **a.** Variable levels of microglia pathology and morphological changes are seen in key central autonomic nuclei of the medulla. **b.** Focal loss of P2Y12 levels and marked distribution heterogeneity of microglia are present in the thalamus. Thalamic nuclei: MD mediodorsal, VL ventral lateral, VPL ventral posterolateral. **c.** In sites of severe microglial pathologies different microglial states ranging from marked morphological transformation to degeneration and cell loss are seen. Transmission electron microscopic (TEM) images show degenerating microglia in the medulla (yellow pseudocolor). Pink arrowheads show degenerating microglia processes. Scale bars: a. 10 µm, b. 5000 µm; 200 µm; c, 5 µm.

**Figure S3.**
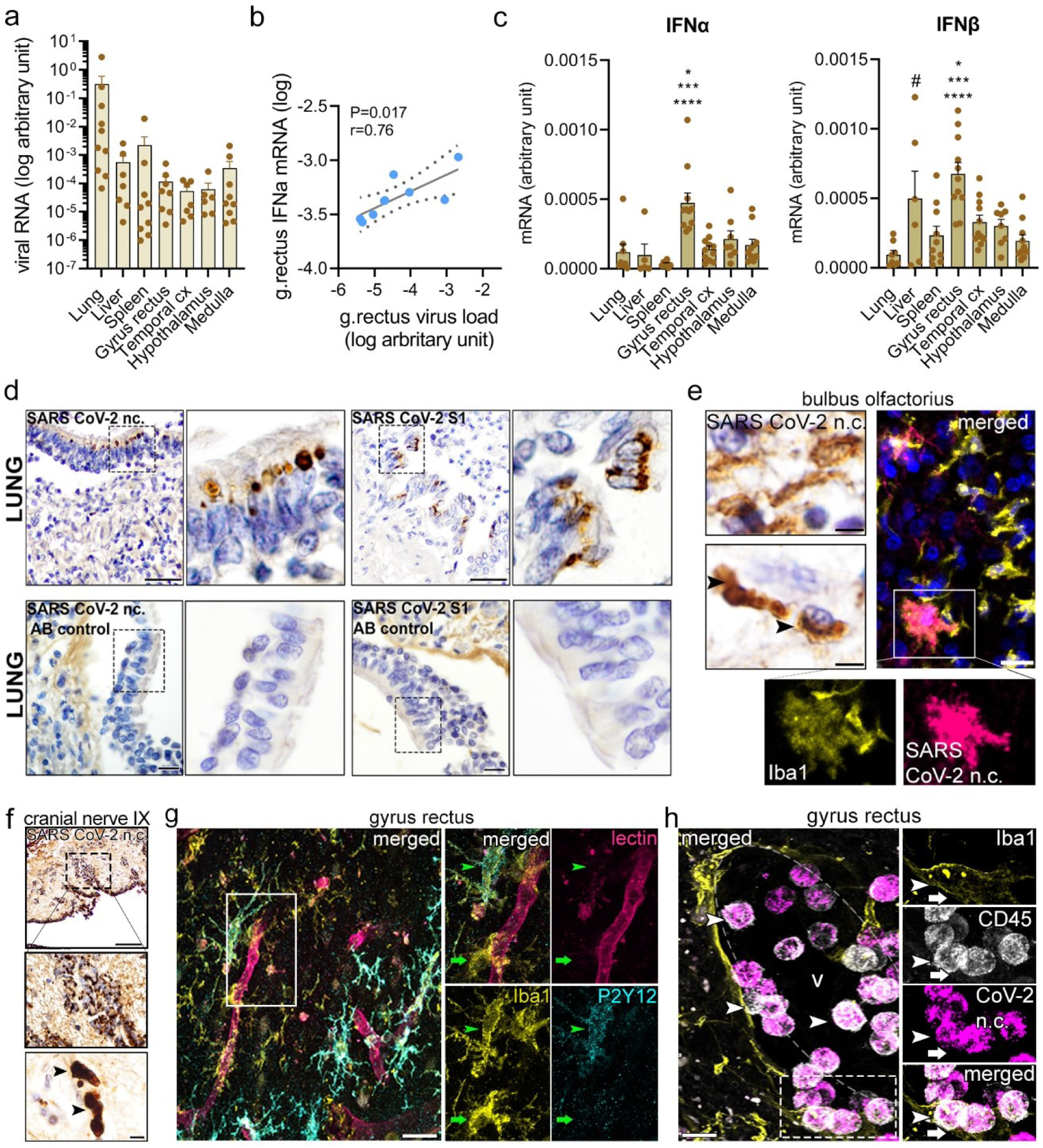
Accumulation of viral antigens in the brain parallels the development of a type I interferon response and downregulation of microglial P2Y12R. **a**. Viral mRNA levels were measured by qPCR and presented as log arbitrary units. **b**. Virus load shows a positive correlation with IFNα mRNA levels in the gyrus rectus (individual samples where SARS CoV-2 mRNA levels were detectable were included). **c**. IFNβ and IFNα mRNA levels in tissue homogenates as measured by qPCR. One-way ANOVA with Tukey’s post-hoc test. IFNβ: \*\*\*\**p*<0.0001 vs lung, \*\*\**p*<0.001 vs spleen and medulla, \**p*<0.05 vs temporal cx and hypothalamus. ^#^*p*<0.05 vs lung. IFNα: \*\*\*\**p*<0.0001 vs spleen, \*\*\**p*<0.001 vs lung, liver, temporal cx and medulla, \**p*<0.05 vs hypothalamus. **d**. SARS CoV-2 nuclecapsid and S1 protein labelings in lung epithelial cells. Lack of primary antibody eliminates specific immnopositivity for viral antigens on subsequent lung tissue sections. **e**. Immunodetection of SARS CoV-2 nucleocapsid (arrowheads) in paraffin embedded bulbus olfactorius with cresyl violet counterstain (left panels). A subset of microglia accumulate viral antigens as visualized by immunofluorescence at infected areas of the bulbus (right panels). **f**. Cranial nerve IX presents SARS CoV-2 nucleocapsid immunopositive profiles (arrowheads). **g**. Loss of P2Y12R from vessel-associated microglia and microglial processes (arrowhead) with some P2Y12R retained on microglia away from the vessel (arrow) is shown in the gyrus rectus in a COVID-19 case. **h**. Perivascular CD45-positive immune cells (arrowheads) containing SARS CoV-2 nucleocapsid are contacted and internalized by Iba positive microglia (arrows) in the gyrus rectus. Scale bars: d : 100 µm, 10 µm ; e : 10 µm ; f : 100 µm, 5 µm ; g-h : 10 µm.

**Figure S4.**
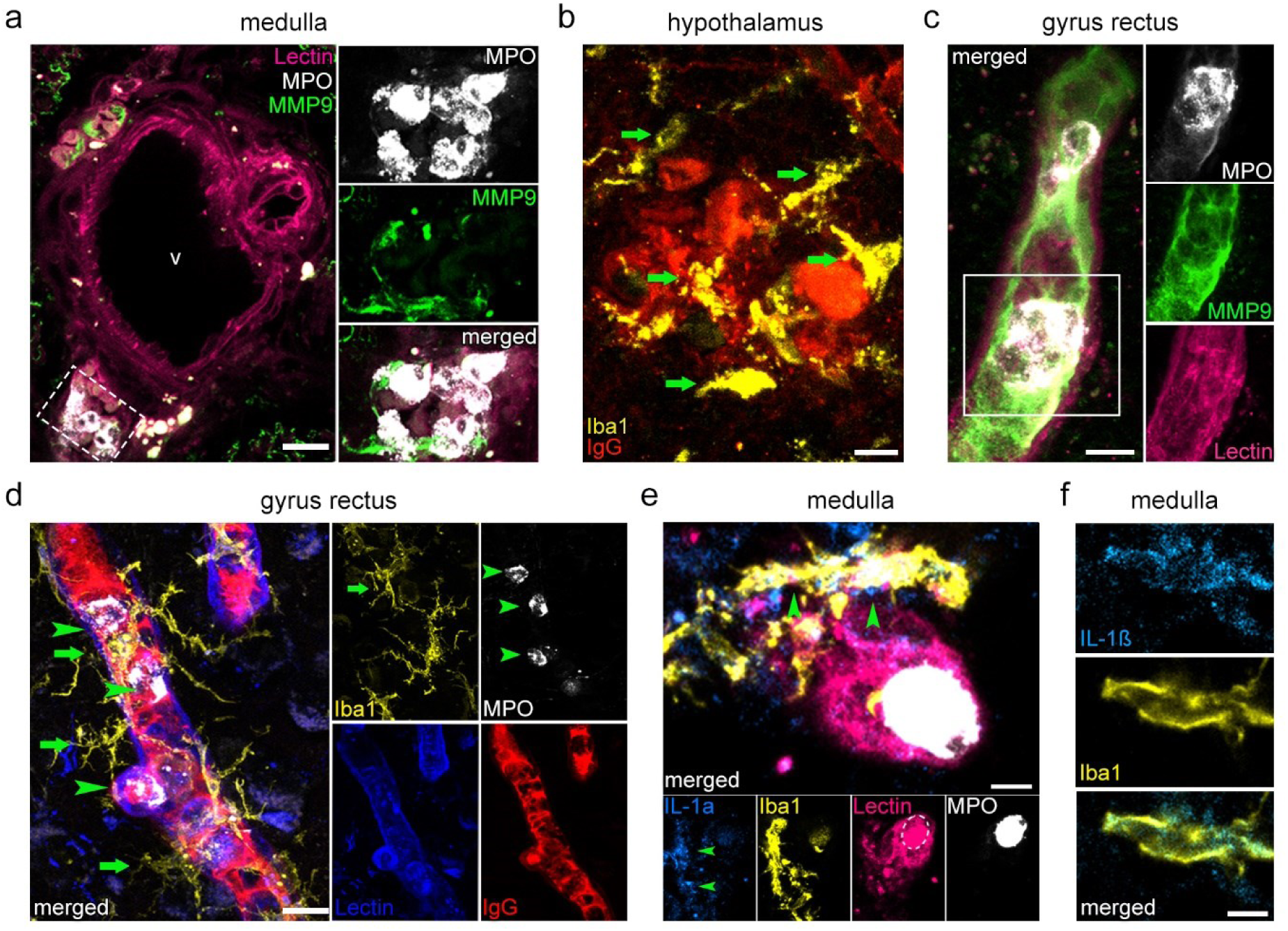
Vascular inflammation and BBB injury are associated with perivascular microglial pathologies in COVID-19. **a**. Iba1-positive microglia (yellow, green arrows) showing marked morphological transformation are recruited to blood vessels (lectin, pink) with intraluminal MPO positive cells (white, green arrowheads. Plasma leakage through disrupted blood vessels into the brain parenchyma is indicated by the presence of IgG (blue, white arrows). **b**. Microglia with markedly altered morphology (Iba1, yellow, green arrows) are recruited to sites of parenchymal IgG deposition (red) in the hypothalamus. **c**. Intravascular leukocytes in the gyrus rectus are associated with MMP9. **d**. Intravascular MPO-positive leukocytes (arrowheads) associate with IgG and vessel-associated microglia (arrows) in the gyrus rectus. **e**. Perivascular Iba-1 positive microglia (yellow, green arrowheads) expressing the pro-inflammatory cytokine IL-1α (blue) in close proximity to an intraluminal MPO positive leukocyte (white) in the medulla. **f**. The pro-inflammatory cytokine IL-1ß (blue) is expressed in Iba-1 positive microglia (yellow). Scale bars: a,b,c,d: 10μm; e,f: 5 μm.

**Figure S5.**
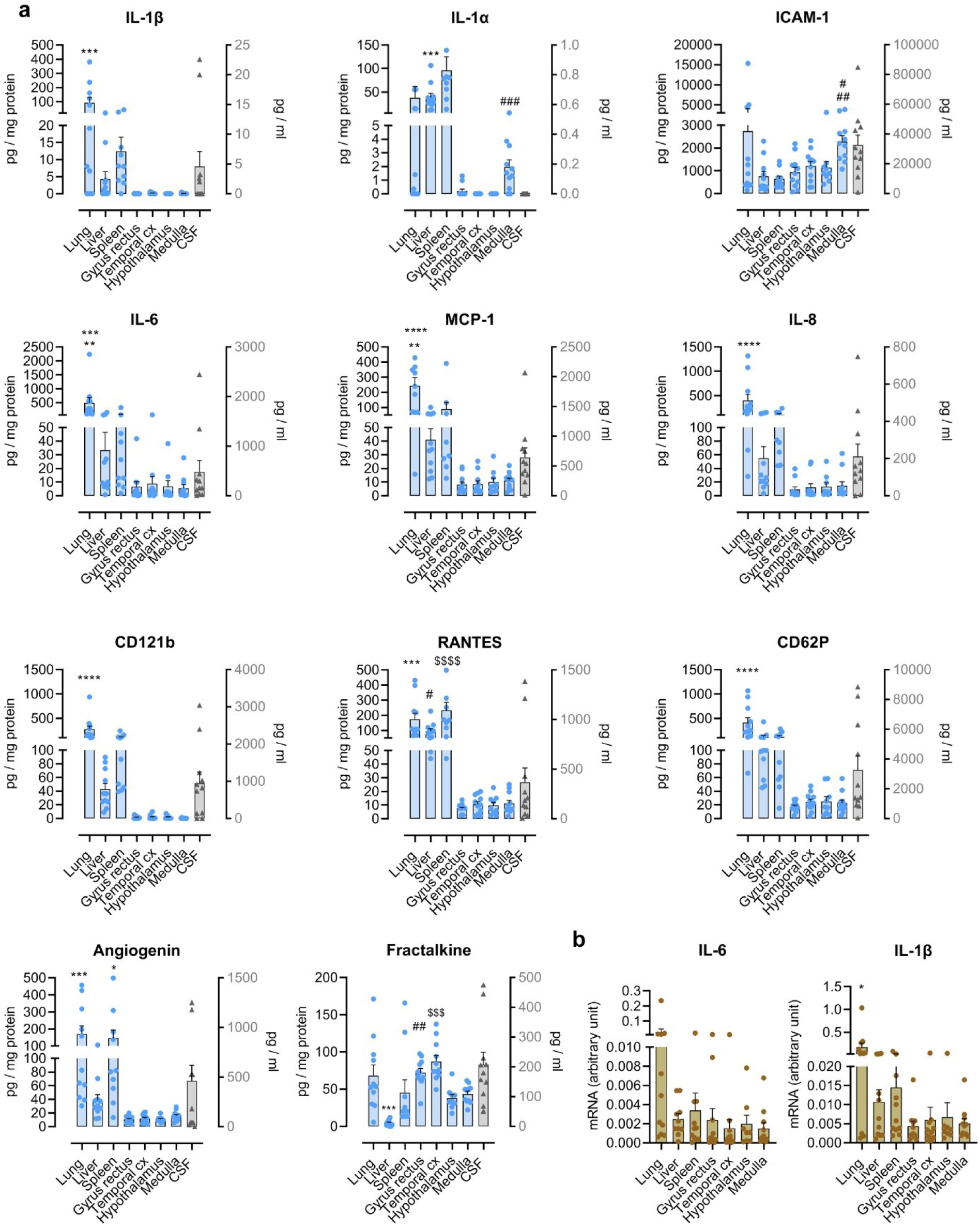
Inflammatory mediators in the brain and peripheral tissues in COVID-19 cases. **a.** Cytokine and chemokin levels were measured by cytometric bead array in tissue homogenates (blue bars, left Y axis, expressed as pg/mg protein) and in the CSF (gray bar, right Y axis, expressed as pg/ml). One-way ANOVA with Tukey’s post hoc test. IL-1β: \*\*\**p*<0.001 vs all areas, IL-6: \*\*\**p*<0.001 vs brain areas, \*\**p*<0.01 vs peripheral organs, MCP-1: \*\*\*\**p*<0.0001 vs brain areas, \*\**p*<0.01 vs peripheral organs, IL-8 \*\*\*\**p*<0.0001 vs all areas, CD62P \*\*\*\**p*<0.0001 vs all areas, Angiogenin: \*\*\**p*<0.001 vs brain areas, \**p*<0.05 vs brain areas, CD121b: \*\*\**p*<0.001 vs brain areas, RANTES: \*\*\**p*<0.001, ^#^*p*<.0.05, ^$$$$^P <0.0001 vs all brain areas. Fractalkine: \*\*\**p*<0.001 vs all tissues, ^##^*p*<.0.01 gyrus rectus vs hypothalamus and medulla, ^$$$^*p*<.0.001 temporal cx vs hypothalamus and medulla when brain areas are compared. Cytokines TNFα, IFNγ and IL-17A were below detection levels in most tissues (not shown). **b**. IL-6 and IL-1β mRNA levels were measured by qPCR in tissue homogenates (\**p*<0.05 vs all tissues).

**Figure S6.**
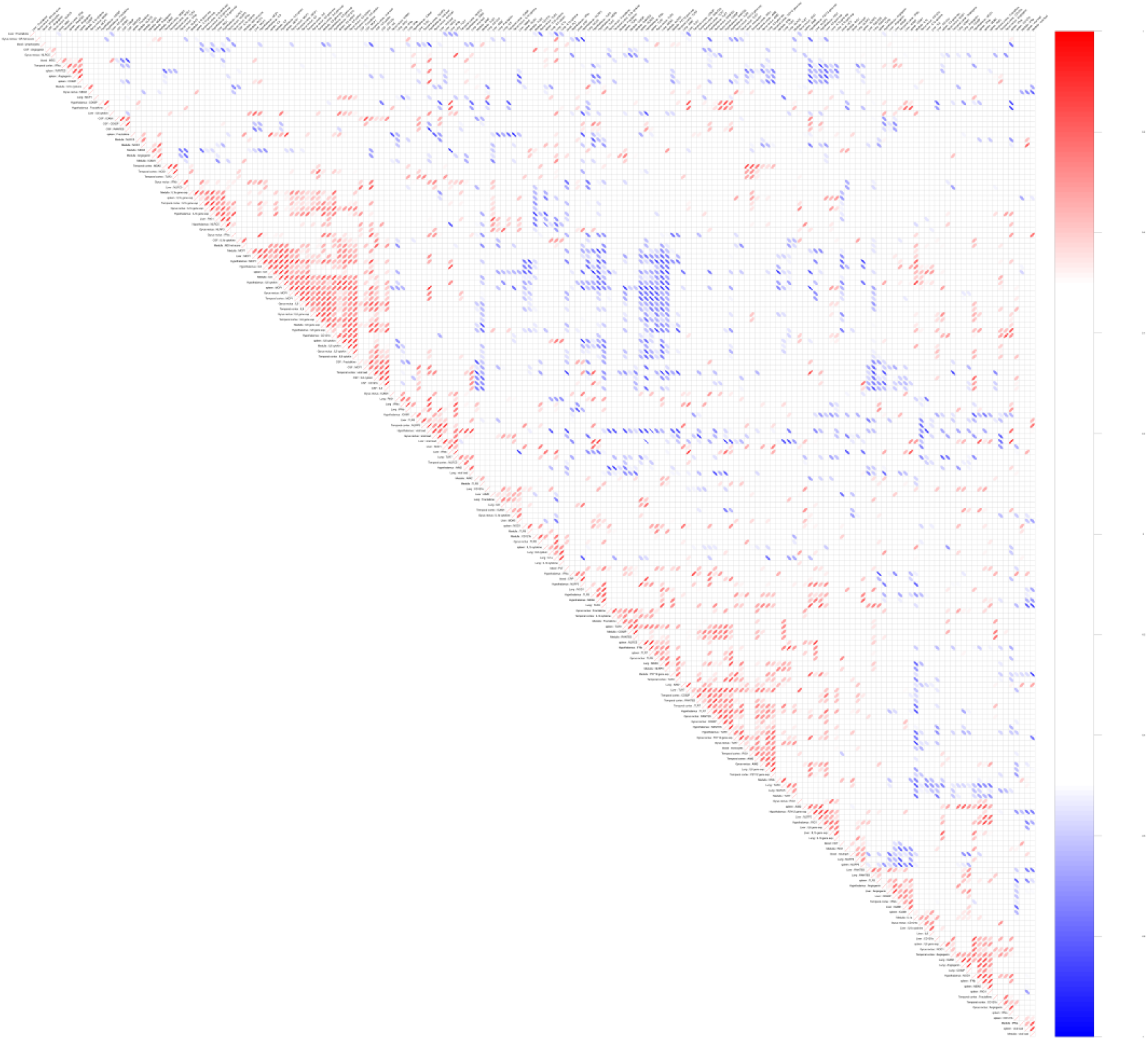
Correlation matrix showing associations between inflammatory mediators, microglial states and PRRs in COVID-19 cases. Pearson correlation was calculated in all combinations, *p*>0.05 or -0.5 <r>0.5 were masked out. Visualization was done with R package: corrplot, sing hclust on default. Red is for positive; blue for negative correlation, donut shape and color indicate p values and data spread, respectively. *Please download the folder containing Figure S6 from the drive below, and unzip to visualize html file*.

**Figure S7.**
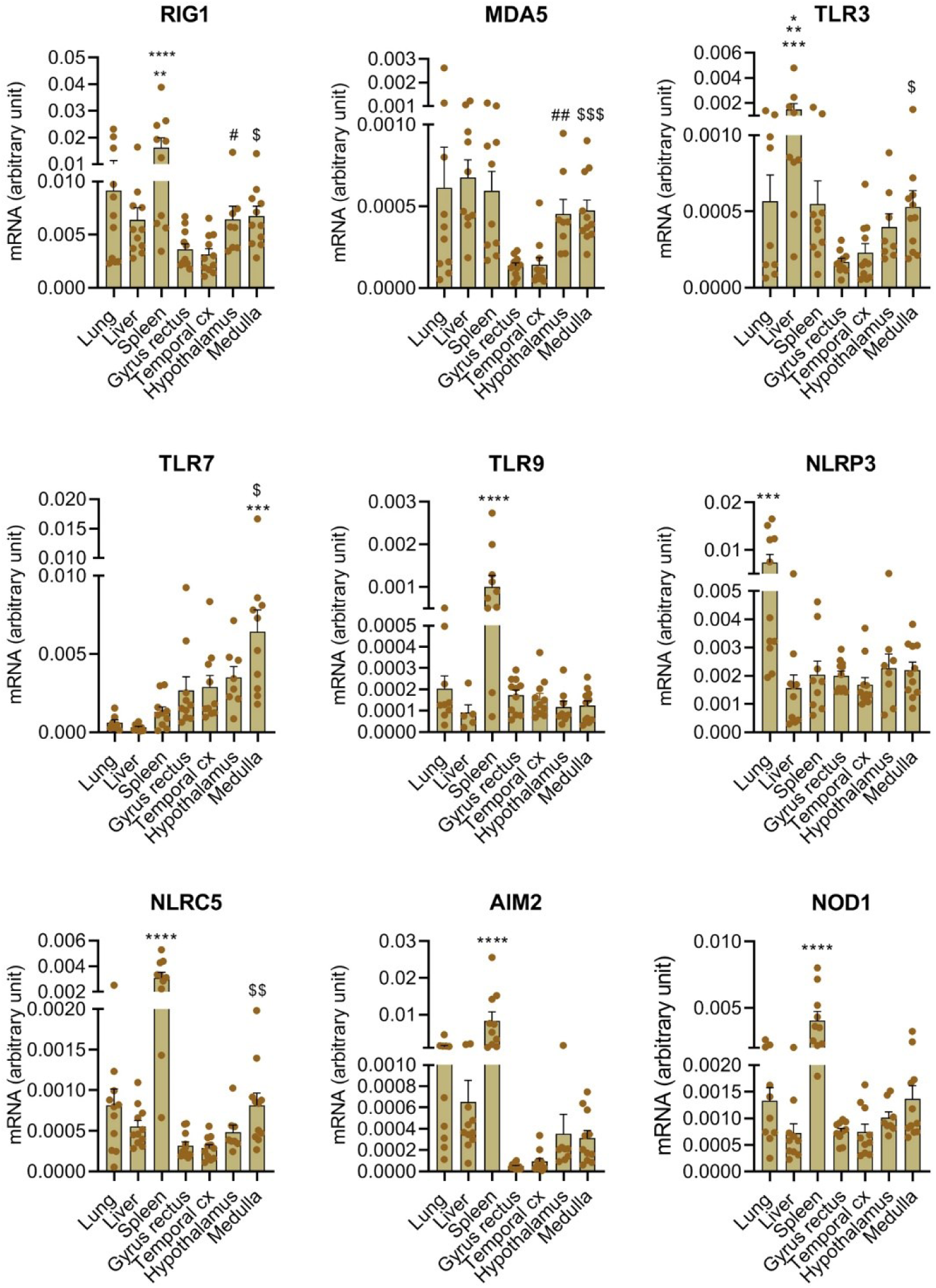
Expression of pattern recognition receptors in the brain and peripheral tissues in COVID-19 cases. mRNA levels of different PRRs were measured by qPCR in tissue homogenates. One-way ANOVA with Tukey’s post hoc test. RIG1: \*\**p*<0.01 vs spleen, hypothalamus and medulla, \*\*\*\**p*<0.0001 vs gyrus rectus and temporal cx. ^#, $^*p*<0.05 vs gyrus rectus and temporal cx when brain areas compared. MDA5: ^##^*p*<0.01, ^$$$^*p*<0.0001 vs gyrus rectus and temporal cx when brain areas compared. TLR3: \**p*<0.05 vs spleen, lung and medulla, \*\**p*<0.01 vs hypothalamus, \*\*\**p*<0.001 vs cortial areas. ^$^*p*<0.05 vs gyrus rectus and temporal cortex when brain areas compared. TLR7: \*\*\**p*<0.001 vs lung, liver and spleen, ^$^P<0.05 vs gyrus rectus and temporal cortex when brain areas compared. TLR9 \*\*\*\**p*<0.0001 vs all tissues; NLRP3: \*\*\**p*<0.001 vs all tissues; NLRC5: \*\*\*\**p*<0.0001 vs all tissues, ^$$^*p*<0.01 vs gyrus rectus and temporal cortex when brain areas compared; AIM2: \*\*\*\**p*<0.0001 vs all tissues; NOD1: \*\*\*\**p*<0.0001 vs all tissues.

**Figure S8.**
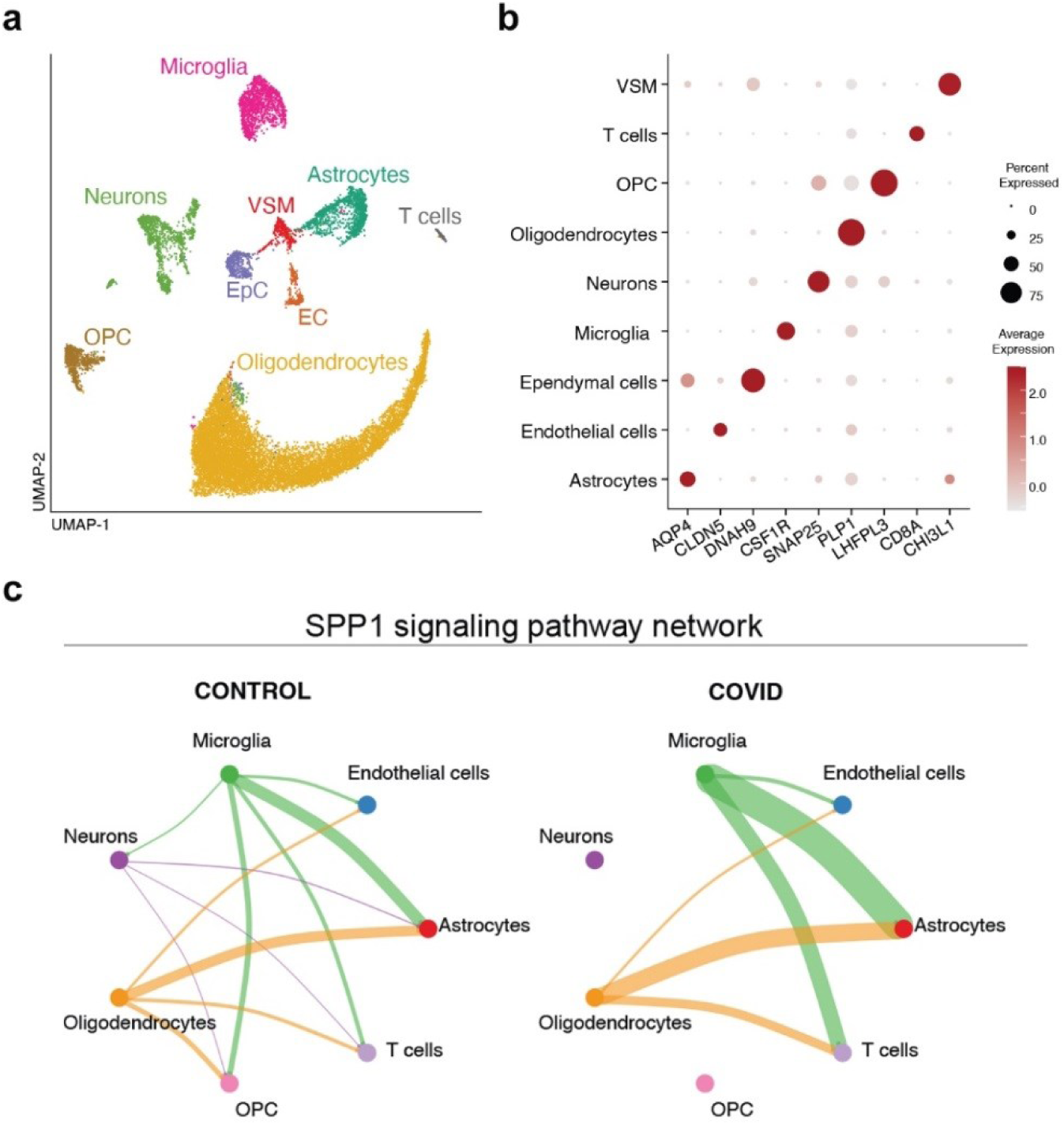
Single cell RNA sequencing in the medulla of control and COVID-19 cases. **a.** Uniform Manifold Approximation and Projection (UMAP) plot of a total of 16,260 cells, colored by identified populations (same as Fig.5a). **b.** Dot plot showing the expression profile of selected genes key for the identification of all brain cell populations. The dot size corresponds to the fraction of cells within each condition expressing the indicated transcript, and the color indicates average expression. **c**. SPP1 signaling pathway network analysis.

## Supplementary tables and table legends

**Table S1.** Clinical data of control cases involved in the study.

| **Control case No.** | **Age** | **Sex** | **Comorbidities** | **Cause of death** | **Post mortem delay (hour)** | **Fixation method and time** |
| --- | --- | --- | --- | --- | --- | --- |
| sko17 | 75 | female | atherosclerosis | acute transmural infarction of inferior wall | 4,5 hour | perfused plus post fixed 24 hours |
| sko16 | 72 | male | atherosclerosis, abscess of lung with pneumonia, acute bronchitis | respiratory arrest | 2,5 hour | perfused plus post fixed 24 hours |
| sko19 | 61 | female | atherosclerosis, diabetes, hypertension | acute transmural infarction of inferior wall | 3 hour | perfused plus post fixed 24 hours |
| sko24 | 79 | male | pulmonary heart disease, emphysema, calculus of gallbladder with other cholecystitis | left ventricular failure | 2,5 hour | perfused plus post fixed 24 hours |
| sko27 | 76 | female | atherosclerosis | small intestine death | 3 hour 15 mins | perfused plus post fixed 24 hours |
| sko28 | 93 | female | hypertension, atherosclerosis, diabetes mellitus, nephrosclerosis arteriosclerotica | left coronary artery occlusion | 4 hours | perfused plus post fixed 24 hours |
| #133 | 77 | male | long-term psychiatric treatment without previous suicide attempt arteriosclerosis, coronary sclerosis, renal arteriosclerosis | suicide (hanging) | 4 hours | frozen |
| #253 | 68 | male | acute pulmonary oedema serious arteriosclerosis (especially in the heart and kidney) peripheral arterial shunt cerebral sclerosis<br>left coronary occlusion | acute cardiac insufficiency | 10 hours | frozen |
| #256 | 67 | male | rapid pneumonia,<br>cardiopulmonary<br>insufficiency<br>duodenal ulcer<br>diabetes mellitus<br>hypertension<br>multiple abdominal<br>tumor metastasis,<br>duodenal ulcer | pancreas cancer | 10 hours | frozen |

**Table S2.** Clinical data of COVID-19 cases involved in the study.

| **Covid case No.** | **Age** | **Sex** | **Time of hospital admission until death (day)** | **Time spent at ICU (day)** | **Cause of death** | **Comorbidities** | **Vaccinated against COVID** | **COVID status** | **Post mortem delay (hour)** | **Fixation method and time** |
| --- | --- | --- | --- | --- | --- | --- | --- | --- | --- | --- |
| case1 | 87 | female | 55 days | 19 days | pneumonia | high blood pressure, vascular dementia | No | Positive | 7 h | Fresh-sampled, imersion fixed for 24 hours |
| case2 | 75 | female | 13 days | 13 days | ileus | diabetes, atherosclerosis | No | Positive | 17 h | Fresh-sampled, imersion fixed for 24 hours |
| case3 | 87 | female | 60 days | 33 days | pneumonia, respiratory arrest | high blood pressure | No | Positive | 15 h | Fresh-sampled, imersion fixed for 24 hours |
| case4 | 99 | female | 11 days | 3 days | pneumonia | atherosclerosis, Alzheimer disease | No | Positive | 9 h | Fresh-sampled, imersion fixed for 24 hours |
| case5 | 74 | male | 4 days | 4 days | pneumonia, congestive heart failure | hypertonia, hypothyreosis | No | Positive | 9,5 h | Fresh-sampled, imersion fixed for 24 hours |
| case6 | 88 | female | 1 day | 1 day | pneumonia, cardiac arrest | anaemia | No | Positive | 9,5 h | Fresh-sampled, imersion fixed for 24 hours |
| case7 | 66 | male | 19 days | 17 days | pneumonia, cardiac arrest | diabetes | No | Positive | 7 h | Fresh-sampled, imersion fixed for 24 hours |
| case8 | 66 | male | 5 days | 5 days | pneumonia,<br>cardiac arrest | Parkinson<br>disease | No | Positive | 7 h | Fresh-sampled,<br>imersion fixed<br>for 24 hours |
| case9 | 72 | male | 9 days | 9 days | pneumonia,<br>cardiac arrest | diabetes | No | Positive | 7 h | Fresh-sampled,<br>imersion fixed<br>for 24 hours |
| case10 | 66 | male | 5 days | 5 days | cardiac arrest | hypertonia | No | Positive | 5,5 h | Fresh-sampled,<br>imersion fixed<br>for 24 hours |
| case11 | 67 | male | 6 days | 6 days | pneumonia,<br>left<br>ventricular<br>failure | hypertonia,<br>diabetes | No | Positive | 4 h | Fresh-sampled,<br>imersion fixed<br>for 24 hours |
| case12 | 57 | male | 7 days | 7 days | respiratory<br>arrest | none | No | Positive | 3 h | Perfused plus post<br>fixed 24 hours |
| case13 | 63 | male | 14 days | 10 days | penumonia | atherosclerosis | Yes | Positive | 4 h | Perfused plus post<br>fixed 24 hours |

Table S3. Pearson correlation analysis results

Table S4. Differentially expressed genes, single nuclear RNA sequencing

**Table S5.** List of primary and secondary antibodies used.

| **Primary antibodies** | **host** | **source** | **catalog no.** |
| --- | --- | --- | --- |
| Aquaporin4 | guineapig | Synaptic Systems | 429 004 |
| CD68 | mouse | Leica | NCL-L-CD68 |
| CD45 | mouse | Leica | NCL-L-LCA_1 ml |
| Homer-1 | chicken | Synaptic Systems | 160 006 |
| Iba1 | guineapig | Synaptic Systems | 234 308 |
| Iba1 | chicken | Synaptic Systems | 234 009 |
| ICAM-1 | mouse | R&D Systems | BBA3 |
| IL-1a | mouse | R&D Systems | MAB-200-100 |
| IL-1β | mouse | R&D Systems | MAB-201-100 |
| Kv 2.1 clone K89/34 | mouse | NeuroMab | 75-014 |
| Lectin | tomato | Vectorlabs | B-1175 |
| MAP-2 | guineapig | Synaptic Systems | 188 004 |
| MBP | rat | Abcam | ab7349 |
| MMP-9 | mouse | StressMarq | SMC-396D |
| MPO | rabbit | BioCare Medical | PP023 AA |
| P2Y12 | rabbit | Anaspec | TL1403 |
| SARS CoV-2 nucleocapsid | rabbit | Novusbio | NB100-56576 |
| SARS CoV-2 Spike1 protein | rabbit | GeneTex | GTX635654-25 |
| Synapsin I | mouse | Synaptic Systems | 106 011 |
| P-Stat3 | rabbit | Cell Signaling | 9131S |
| vGlut1 | guineapig | Synaptic Systems | 135 304 |
| Tom20 | rabbit | Invitrogen | ST04-72 |
| **Secondary antibodies** |  |  |  |
| biotinilated anti-rabbit | goat | Vector | BA-1000 |
| biotinilated anti-guineapig | donkey | Jackson | 706-066-148 |
| DyLight 405 anti- | donkey | Jackson | 016-470-084 |
| Alexa 488 anti-rabbit | donkey | Jackson | 711-546-152 |
| Alexa 594 anti-guineapig | donkey | Jackson | 706-586-148 |
| Alexa 594 anti-mouse | donkey | Jackson | 715-586-151 |
| Alexa 594 anti-rat | goat | LifeTech | A11007 |
| Alexa 647 anti-guineapig | donkey | Jackson | 706-606-148 |
| Alexa 647 anti-rabbit | donkey | Jackson | 711-605-152 |
| Alexa 647 anti-chicken | donkey | Jackson | 703-606-155 |
| **Gold conjugated secondary antibodies** |  |  |  |
| Nanogold Fab' anti-rabbit | goat | Nanoprobes | 38C510 |

## References

1. Long-term neurological sequelae of SARS-CoV-2 infection. Nature medicine 28, 2269 (Nov, 2022).

2. P. Najt, H. L. Richards, D. G. Fortune, Brain imaging in patients with COVID-19: A systematic review. Brain, behavior, & immunity - health 16, 100290 (Oct, 2021).

3. A. Maiese et al., SARS-CoV-2 and the brain: A review of the current knowledge on neuropathology in COVID-19. Brain Pathol 31, e13013 (Nov, 2021).

4. E. M. Conway et al., Understanding COVID-19-associated coagulopathy. Nature reviews. Immunology 22, 639 (Oct, 2022).

5. G. Goshua et al., Endotheliopathy in COVID-19-associated coagulopathy: evidence from a single-centre, cross-sectional study. The Lancet. Haematology 7, e575 (Aug, 2020).

6. J. Meinhardt et al., Olfactory transmucosal SARS-CoV-2 invasion as a port of central nervous system entry in individuals with COVID-19. Nature neuroscience 24, 168 (Feb, 2021).

7. R. W. Paterson et al., The emerging spectrum of COVID-19 neurology: clinical, radiological and laboratory findings. Brain : a journal of neurology 143, 3104 (Oct 1, 2020).

8. G. Douaud et al., SARS-CoV-2 is associated with changes in brain structure in UK Biobank. Nature 604, 697 (Apr, 2022).

9. S. Spudich, A. Nath, Nervous system consequences of COVID-19. Science 375, 267 (Jan 21, 2022).

10. H. E. Davis, L. McCorkell, J. M. Vogel, E. J. Topol, Long COVID: major findings, mechanisms and recommendations. Nature reviews. Microbiology 21, 133 (Mar, 2023).

11. M. Ramos-Casals, P. Brito-Zeron, X. Mariette, Systemic and organ-specific immune-related manifestations of COVID-19. Nature reviews. Rheumatology 17, 315 (Jun, 2021).

12. Y. Wang, S. Perlman, COVID-19: Inflammatory Profile. Annual review of medicine 73, 65 (Jan 27, 2022).

13. M. Schwabenland et al., Deep spatial profiling of human COVID-19 brains reveals neuroinflammation with distinct microanatomical microglia-T-cell interactions. Immunity 54, 1594 (Jul 13, 2021).

14. A. C. Yang et al., Dysregulation of brain and choroid plexus cell types in severe COVID-19. Nature 595, 565 (Jul, 2021).

15. T. E. Poloni et al., COVID-19-related neuropathology and microglial activation in elderly with and without dementia. Brain Pathol 31, e12997 (Sep, 2021).

16. J. A. Hosp et al., Cognitive impairment and altered cerebral glucose metabolism in the subacute stage of COVID-19. Brain : a journal of neurology 144, 1263 (May 7, 2021).

17. N. Deigendesch et al., Correlates of critical illness-related encephalopathy predominate postmortem COVID-19 neuropathology. Acta neuropathologica 140, 583 (Oct, 2020).

18. K. T. Thakur et al., COVID-19 neuropathology at Columbia University Irving Medical Center/New York Presbyterian Hospital. Brain : a journal of neurology 144, 2696 (Oct 22, 2021).

19. J. Matschke et al., Neuropathology of patients with COVID-19 in Germany: a post-mortem case series. The Lancet. Neurology 19, 919 (Nov, 2020).

20. A. Vanderheiden, R. S. Klein, Neuroinflammation and COVID-19. Current opinion in neurobiology 76, 102608 (Oct, 2022).

21. R. Butowt, N. Meunier, B. Bryche, C. S. von Bartheld, The olfactory nerve is not a likely route to brain infection in COVID-19: a critical review of data from humans and animal models. Acta neuropathologica 141, 809 (Jun, 2021).

22. S. R. Stein et al., SARS-CoV-2 infection and persistence in the human body and brain at autopsy. Nature 612, 758 (Dec, 2022).

23. E. A. Albornoz et al., SARS-CoV-2 drives NLRP3 inflammasome activation in human microglia through spike protein. Molecular psychiatry, (Nov 1, 2022).

24. A. Denes, S. M. Allan, T. Hortobagyi, C. J. Smith, Studies on inflammation and stroke provide clues to pathomechanism of central nervous system involvement in COVID-19. Free Neuropathol 1, (Jan, 2020).

25. B. Martinez-Salazar et al., COVID-19 and the Vasculature: Current Aspects and Long-Term Consequences. Frontiers in cell and developmental biology 10, 824851 (2022).

26. S. E. Haynes et al., The P2Y12 receptor regulates microglial activation by extracellular nucleotides. Nature neuroscience 9, 1512 (Dec, 2006).

27. E. Csaszar et al., Microglia modulate blood flow, neurovascular coupling, and hypoperfusion via purinergic actions. The Journal of experimental medicine 219, (Mar 7, 2022).

28. K. Bisht et al., Capillary-associated microglia regulate vascular structure and function through PANX1-P2RY12 coupling in mice. Nature communications 12, 5289 (Sep 6, 2021).

29. B. Rossi, S. Angiari, E. Zenaro, S. L. Budui, G. Constantin, Vascular inflammation in central nervous system diseases: adhesion receptors controlling leukocyte-endothelial interactions. Journal of leukocyte biology 89, 539 (Apr, 2011).

30. A. Tello-Montoliu, J. V. Patel, G. Y. Lip, Angiogenin: a review of the pathophysiology and potential clinical applications. Journal of thrombosis and haemostasis : JTH 4, 1864 (Sep, 2006).

31. B. Vafadari, A. Salamian, L. Kaczmarek, MMP-9 in translation: from molecule to brain physiology, pathology, and therapy. Journal of neurochemistry 139 Suppl 2, 91 (Oct, 2016).

32. A. Denes, P. Thornton, N. J. Rothwell, S. M. Allan, Inflammation and brain injury: acute cerebral ischaemia, peripheral and central inflammation. Brain, behavior, and immunity 24, 708 (Jul, 2010).

33. C. A. Dinarello, Interleukin-1 and interleukin-1 antagonism. Blood 77, 1627 (Apr 15, 1991).

34. S. M. Allan, P. J. Tyrrell, N. J. Rothwell, Interleukin-1 and neuronal injury. Nature reviews. Immunology 5, 629 (Aug, 2005).

35. D. M. Del Valle et al., An inflammatory cytokine signature predicts COVID-19 severity and survival. Nature medicine 26, 1636 (Oct, 2020).

36. F. Perez-Garcia et al., High SARS-CoV-2 Viral Load and Low CCL5 Expression Levels in the Upper Respiratory Tract Are Associated With COVID-19 Severity. The Journal of infectious diseases 225, 977 (Mar 15, 2022).

37. M. Eldahshan et al., Prognostic Significance of Platelet Activation Marker CD62P in Hospitalized Covid-19 Patients. Clinical laboratory 68, (Sep 1, 2022).

38. B. K. Patterson et al., Persistence of SARS CoV-2 S1 Protein in CD16+ Monocytes in Post Acute Sequelae of COVID-19 (PASC) up to 15 Months Post-Infection. Frontiers in immunology 12, 746021 (2021).

39. A. E. Cardona et al., Control of microglial neurotoxicity by the fractalkine receptor. Nature neuroscience 9, 917 (Jul, 2006).

40. A. S. Mendiola et al., Fractalkine Signaling Attenuates Perivascular Clustering of Microglia and Fibrinogen Leakage during Systemic Inflammation in Mouse Models of Diabetic Retinopathy. Frontiers in cellular neuroscience 10, 303 (2016).

41. P. Broz, V. M. Dixit, Inflammasomes: mechanism of assembly, regulation and signalling. Nature reviews. Immunology 16, 407 (Jul, 2016).

42. M. S. Diamond, T. D. Kanneganti, Innate immunity: the first line of defense against SARS-CoV-2. Nature immunology 23, 165 (Feb, 2022).

43. S. Jin et al., Inference and analysis of cell-cell communication using CellChat. Nature communications 12, 1088 (Feb 17, 2021).

44. L. A. O’Neill, R. J. Kishton, J. Rathmell, A guide to immunometabolism for immunologists. Nature reviews. Immunology 16, 553 (Sep, 2016).

45. C. Cserep, B. Posfai, A. Denes, Shaping Neuronal Fate: Functional Heterogeneity of Direct Microglia-Neuron Interactions. Neuron, (Nov 20, 2020).

46. N. Holderith, J. Heredi, V. Kis, Z. Nusser, A High-Resolution Method for Quantitative Molecular Analysis of Functionally Characterized Individual Synapses. Cell reports 32, 107968 (Jul 28, 2020).

47. G. Cosentino et al., Neuropathological findings from COVID-19 patients with neurological symptoms argue against a direct brain invasion of SARS-CoV-2: A critical systematic review. European journal of neurology 28, 3856 (Nov, 2021).

48. G. M. Olivarria et al., Microglia Do Not Restrict SARS-CoV-2 Replication following Infection of the Central Nervous System of K18-Human ACE2 Transgenic Mice. Journal of virology 96, e0196921 (Feb 23, 2022).

49. K. Saijo, A. Crotti, C. K. Glass, Regulation of microglia activation and deactivation by nuclear receptors. Glia 61, 104 (Jan, 2013).

50. J. J. Neher, C. Cunningham, Priming Microglia for Innate Immune Memory in the Brain. Trends in immunology 40, 358 (Apr, 2019).

51. C. D. Russell, N. I. Lone, J. K. Baillie, Comorbidities, multimorbidity and COVID-19. Nature medicine 29, 334 (Feb, 2023).

52. A. G. Laing et al., A dynamic COVID-19 immune signature includes associations with poor prognosis. Nature medicine 26, 1623 (Oct, 2020).

53. C. Qin et al., Dysregulation of Immune Response in Patients With Coronavirus 2019 (COVID-19) in Wuhan, China. Clinical infectious diseases : an official publication of the Infectious Diseases Society of America 71, 762 (Jul 28, 2020).

54. Y. Chen et al., IP-10 and MCP-1 as biomarkers associated with disease severity of COVID-19. Mol Med 26, 97 (Oct 29, 2020).

55. D. Reinhold et al., The brain reacting to COVID-19: analysis of the cerebrospinal fluid proteome, RNA and inflammation. Journal of neuroinflammation 20, 30 (Feb 9, 2023).

56. V. Salvi, et al., SARS-CoV-2-associated ssRNAs activate inflammation and immunity via TLR7/8. JCI insight 6, (Sep 22, 2021).

57. X. Yin et al., MDA5 Governs the Innate Immune Response to SARS-CoV-2 in Lung Epithelial Cells. Cell reports 34, 108628 (Jan 12, 2021).

58. D. Farkas, et al., A role for Toll-like receptor 3 in lung vascular remodeling associated with SARS-CoV-2 infection. bioRxiv, (Jan 25, 2023).

59. R. Fekete et al., Microglia control the spread of neurotropic virus infection via P2Y12 signalling and recruit monocytes through P2Y12-independent mechanisms. Acta neuropathologica 136, 461 (Sep, 2018).

60. D. L. Wheeler, A. Sariol, D. K. Meyerholz, S. Perlman, Microglia are required for protection against lethal coronavirus encephalitis in mice. The Journal of clinical investigation 128, 931 (Mar 1, 2018).

61. C. Cserep et al., Microglia monitor and protect neuronal function through specialized somatic purinergic junctions. Science 367, 528 (Jan 31, 2020).

62. W. Wu et al., Microglial depletion aggravates the severity of acute and chronic seizures in mice. Brain, behavior, and immunity 89, 245 (Oct, 2020).

63. P. Mastorakos et al., Temporally distinct myeloid cell responses mediate damage and repair after cerebrovascular injury. Nature neuroscience 24, 245 (Feb, 2021).

64. G. Szalay et al., Microglia protect against brain injury and their selective elimination dysregulates neuronal network activity after stroke. Nature communications 7, 11499 (May 3, 2016).

65. T. Kubota, P. K. Gajera, N. Kuroda, Meta-analysis of EEG findings in patients with COVID-19. Epilepsy & behavior : E&B 115, 107682 (Feb, 2021).

66. Y. Qin et al., Long-term microstructure and cerebral blood flow changes in patients recovered from COVID-19 without neurological manifestations. The Journal of clinical investigation 131, (Apr 15, 2021).

67. P. Pawelec, M. Ziemka-Nalecz, J. Sypecka, T. Zalewska, The Impact of the CX3CL1/CX3CR1 Axis in Neurological Disorders. Cells 9, (Oct 13, 2020).

68. Samudyata, et al., SARS-CoV-2 promotes microglial synapse elimination in human brain organoids. Molecular psychiatry 27, 3939 (Oct, 2022).

69. A. Sariol et al., Microglia depletion exacerbates demyelination and impairs remyelination in a neurotropic coronavirus infection. Proceedings of the National Academy of Sciences of the United States of America 117, 24464 (Sep 29, 2020).

70. J. Braga et al., Neuroinflammation After COVID-19 With Persistent Depressive and Cognitive Symptoms. JAMA psychiatry, (May 31, 2023).

71. S. Heindl et al., Automated Morphological Analysis of Microglia After Stroke. Frontiers in cellular neuroscience 12, 106 (2018).

72. P. Bustos et al., Quantitative detection of SARS-CoV-2 RNA in nasopharyngeal samples from infected patients with mild disease. Journal of medical virology 93, 2439 (Apr, 2021).

73. S. Aibar et al., SCENIC: single-cell regulatory network inference and clustering. Nature methods 14, 1083 (Nov, 2017).

74. U. Raudvere et al., g:Profiler: a web server for functional enrichment analysis and conversions of gene lists (2019 update). Nucleic acids research 47, W191 (Jul 2, 2019).

