## Supplemental figure 6 (interactive) for "Infection-induced vascular inflammation in COVID-19 links focal microglial dysfunction with neuropathologies through IL-1/IL-6-related systemic inflammatory states": jquery.maphilight.min.js

```
/*! maphilight 07-02-2018 */
!function(a,b){"function"==typeof define&&define.amd?define(["jquery"],b):b(a.jQuery)}(this,function(a){var b,c,d,e,f,g,h,j,k,l,m;if(c=!!document.createElement("canvas").getContext,b=function(){var a=document.createElement("div");a.innerHTML='<v:shape id="vml_flag1" adj="1" />';var b=a.firstChild;return b.style.behavior="url(#default#VML)",b?"object"==typeof b.adj:!0}(),!c&&!b)return void(a.fn.maphilight=function(){return this});if(c){j=function(a){return Math.max(0,Math.min(parseInt(a,16),255))},k=function(a,b){return"rgba("+j(a.substr(0,2))+","+j(a.substr(2,2))+","+j(a.substr(4,2))+","+b+")"},d=function(b){var c=a('<canvas style="width:'+a(b).width()+"px;height:"+a(b).height()+'px;"></canvas>').get(0);return c.getContext("2d").clearRect(0,0,a(b).width(),a(b).height()),c};var n=function(a,b,c,d,e){if(d=d||0,e=e||0,a.beginPath(),"rect"==b)a.rect(c[0]+d,c[1]+e,c[2]-c[0],c[3]-c[1]);else if("poly"==b)for(a.moveTo(c[0]+d,c[1]+e),i=2;i<c.length;i+=2)a.lineTo(c[i]+d,c[i+1]+e);else"circ"==b&&a.arc(c[0]+d,c[1]+e,c[2],0,2*Math.PI,!1);a.closePath()};e=function(b,c,d,e,f){var g=b.getContext("2d");if(e.shadow){g.save(),"inside"==e.shadowPosition&&(n(g,c,d),g.clip());var h=100*b.width,i=100*b.height;n(g,c,d,h,i),g.shadowOffsetX=e.shadowX-h,g.shadowOffsetY=e.shadowY-i,g.shadowBlur=e.shadowRadius,g.shadowColor=k(e.shadowColor,e.shadowOpacity);var j=e.shadowFrom;j||(j="outside"==e.shadowPosition?"fill":"stroke"),"stroke"==j?(g.strokeStyle="rgba(0,0,0,1)",g.stroke()):"fill"==j&&(g.fillStyle="rgba(0,0,0,1)",g.fill()),g.restore(),"outside"==e.shadowPosition&&(g.save(),n(g,c,d),g.globalCompositeOperation="destination-out",g.fillStyle="rgba(0,0,0,1);",g.fill(),g.restore())}g.save(),n(g,c,d),e.fill&&(g.fillStyle=k(e.fillColor,e.fillOpacity),g.fill()),e.stroke&&(g.strokeStyle=k(e.strokeColor,e.strokeOpacity),g.lineWidth=e.strokeWidth,g.stroke()),g.restore(),e.fade&&a(b).css("opacity",0).animate({opacity:1},100)},f=function(a){a.getContext("2d").clearRect(0,0,a.width,a.height)}}else d=function(b){return a('<var style="zoom:1;overflow:hidden;display:block;width:'+b.width+"px;height:"+b.height+'px;"></var>').get(0)},e=function(b,c,d,e,f){var g,h,i,j;for(var k in d)d[k]=parseInt(d[k],10);g='<v:fill color="#'+e.fillColor+'" opacity="'+(e.fill?e.fillOpacity:0)+'" />',h=e.stroke?'strokeweight="'+e.strokeWidth+'" stroked="t" strokecolor="#'+e.strokeColor+'"':'stroked="f"',i='<v:stroke opacity="'+e.strokeOpacity+'"/>',"rect"==c?j=a('<v:rect name="'+f+'" filled="t" '+h+' style="zoom:1;margin:0;padding:0;display:block;position:absolute;left:'+d[0]+"px;top:"+d[1]+"px;width:"+(d[2]-d[0])+"px;height:"+(d[3]-d[1])+'px;"></v:rect>'):"poly"==c?j=a('<v:shape name="'+f+'" filled="t" '+h+' coordorigin="0,0" coordsize="'+b.width+","+b.height+'" path="m '+d[0]+","+d[1]+" l "+d.join(",")+' x e" style="zoom:1;margin:0;padding:0;display:block;position:absolute;top:0px;left:0px;width:'+b.width+"px;height:"+b.height+'px;"></v:shape>'):"circ"==c&&(j=a('<v:oval name="'+f+'" filled="t" '+h+' style="zoom:1;margin:0;padding:0;display:block;position:absolute;left:'+(d[0]-d[2])+"px;top:"+(d[1]-d[2])+"px;width:"+2*d[2]+"px;height:"+2*d[2]+'px;"></v:oval>')),j.get(0).innerHTML=g+i,a(b).append(j)},f=function(b){var c=a("<div>"+b.innerHTML+"</div>");c.children("[name=highlighted]").remove(),b.innerHTML=c.html()};g=function(a){var b,c=a.getAttribute("coords").split(",");for(b=0;b<c.length;b++)c[b]=parseFloat(c[b]);return[a.getAttribute("shape").toLowerCase().substr(0,4),c]},m=function(b,c){var d=a(b);return a.extend({},c,a.metadata?d.metadata():!1,d.data("maphilight"))},l=function(a){return a.complete?"undefined"!=typeof a.naturalWidth&&0===a.naturalWidth?!1:!0:!1},h={position:"absolute",left:0,top:0,padding:0,border:0};var o=!1;a.fn.maphilight=function(i){return i=a.extend({},a.fn.maphilight.defaults,i),c||o||(a(window).ready(function(){document.namespaces.add("v","urn:schemas-microsoft-com:vml");var b=document.createStyleSheet(),c=["shape","rect","oval","circ","fill","stroke","imagedata","group","textbox"];a.each(c,function(){b.addRule("v\\:"+this,"behavior: url(#default#VML); antialias:true")})}),o=!0),this.each(function(){var j,k,n,o,p,q,r;if(j=a(this),!l(this))return window.setTimeout(function(){j.maphilight(i)},200);if(n=a.extend({},i,a.metadata?j.metadata():!1,j.data("maphilight")),r=j.get(0).getAttribute("usemap"),r&&(o=a('map[name="'+r.substr(1)+'"]'),j.is('img,input[type="image"]')&&r&&o.length>0)){if(j.hasClass("maphilighted")){var s=j.parent();j.insertBefore(s),s.remove(),a(o).unbind(".maphilight")}k=a("<div></div>").css({display:"block",backgroundImage:'url("'+this.src+'")',backgroundSize:"contain",position:"relative",padding:0,width:this.width,height:this.height}),n.wrapClass&&(n.wrapClass===!0?k.addClass(a(this).attr("class")):k.addClass(n.wrapClass)),j.before(k).css("opacity",1e-10).css(h).remove(),b&&j.css("filter","Alpha(opacity=0)"),k.append(j),p=d(this),a(p).css(h),p.height=this.height,p.width=this.width,a(o).bind("alwaysOn.maphilight",function(){q&&f(q),c||a(p).empty(),a(o).find("area[coords]").each(function(){var b,f;f=m(this,n),f.alwaysOn&&(!q&&c&&(q=d(j[0]),a(q).css(h),q.width=j[0].width,q.height=j[0].height,j.before(q)),f.fade=f.alwaysOnFade,b=g(this),c?e(q,b[0],b[1],f,""):e(p,b[0],b[1],f,""))})}).trigger("alwaysOn.maphilight").bind("mouseover.maphilight, focusin.maphilight",function(b){var d,f,h=b.target;if(f=m(h,n),!f.neverOn&&!f.alwaysOn){if(d=g(h),e(p,d[0],d[1],f,"highlighted"),f.groupBy){var i;i=/^[a-zA-Z][\-a-zA-Z]+$/.test(f.groupBy)?o.find("area["+f.groupBy+'="'+a(h).attr(f.groupBy)+'"]'):o.find(f.groupBy);var j=h;i.each(function(){if(this!=j){var a=m(this,n);if(!a.neverOn&&!a.alwaysOn){var b=g(this);e(p,b[0],b[1],a,"highlighted")}}})}c||a(p).append("<v:rect></v:rect>")}}).bind("mouseout.maphilight, focusout.maphilight",function(a){f(p)}),j.before(p),j.addClass("maphilighted")}})},a.fn.maphilight.defaults={fill:!0,fillColor:"000000",fillOpacity:.2,stroke:!0,strokeColor:"ff0000",strokeOpacity:1,strokeWidth:1,fade:!0,alwaysOn:!1,neverOn:!1,groupBy:!1,wrapClass:!0,shadow:!1,shadowX:0,shadowY:0,shadowRadius:6,shadowColor:"000000",shadowOpacity:.8,shadowPosition:"outside",shadowFrom:!1}});
```
