## Supplemental figure 6 (interactive) for "Infection-induced vascular inflammation in COVID-19 links focal microglial dysfunction with neuropathologies through IL-1/IL-6-related systemic inflammatory states": jquery.min.js

```
/*! jQuery v3.3.1 | (c) JS Foundation and other contributors | jquery.org/license */
!function(e,t){"use strict";"object"==typeof module&&"object"==typeof module.exports?module.exports=e.document?t(e,!0):function(e){if(!e.document)throw new Error("jQuery requires a window with a document");return t(e)}:t(e)}("undefined"!=typeof window?window:this,function(e,t){"use strict";var n=[],r=e.document,i=Object.getPrototypeOf,o=n.slice,a=n.concat,s=n.push,u=n.indexOf,l={},c=l.toString,f=l.hasOwnProperty,p=f.toString,d=p.call(Object),h={},g=function e(t){return"function"==typeof t&&"number"!=typeof t.nodeType},y=function e(t){return null!=t&&t===t.window},v={type:!0,src:!0,noModule:!0};function m(e,t,n){var i,o=(t=t||r).createElement("script");if(o.text=e,n)for(i in v)n[i]&&(o[i]=n[i]);t.head.appendChild(o).parentNode.removeChild(o)}function x(e){return null==e?e+"":"object"==typeof e||"function"==typeof e?l[c.call(e)]||"object":typeof e}var b="3.3.1",w=function(e,t){return new w.fn.init(e,t)},T=/^[\s\uFEFF\xA0]+|[\s\uFEFF\xA0]+$/g;w.fn=w.prototype={jquery:"3.3.1",constructor:w,length:0,toArray:function(){return o.call(this)},get:function(e){return null==e?o.call(this):e<0?this[e+this.length]:this[e]},pushStack:function(e){var t=w.merge(this.constructor(),e);return t.prevObject=this,t},each:function(e){return w.each(this,e)},map:function(e){return this.pushStack(w.map(this,function(t,n){return e.call(t,n,t)}))},slice:function(){return this.pushStack(o.apply(this,arguments))},first:function(){return this.eq(0)},last:function(){return this.eq(-1)},eq:function(e){var t=this.length,n=+e+(e<0?t:0);return this.pushStack(n>=0&&n<t?[this[n]]:[])},end:function(){return this.prevObject||this.constructor()},push:s,sort:n.sort,splice:n.splice},w.extend=w.fn.extend=function(){var e,t,n,r,i,o,a=arguments[0]||{},s=1,u=arguments.length,l=!1;for("boolean"==typeof a&&(l=a,a=arguments[s]||{},s++),"object"==typeof a||g(a)||(a={}),s===u&&(a=this,s--);s<u;s++)if(null!=(e=arguments[s]))for(t in e)n=a[t],a!==(r=e[t])&&(l&&r&&(w.isPlainObject(r)||(i=Array.isArray(r)))?(i?(i=!1,o=n&&Array.isArray(n)?n:[]):o=n&&w.isPlainObject(n)?n:{},a[t]=w.extend(l,o,r)):void 0!==r&&(a[t]=r));return a},w.extend({expando:"jQuery"+("3.3.1"+Math.random()).replace(/\D/g,""),isReady:!0,error:function(e){throw new Error(e)},noop:function(){},isPlainObject:function(e){var t,n;return!(!e||"[object Object]"!==c.call(e))&&(!(t=i(e))||"function"==typeof(n=f.call(t,"constructor")&&t.constructor)&&p.call(n)===d)},isEmptyObject:function(e){var t;for(t in e)return!1;return!0},globalEval:function(e){m(e)},each:function(e,t){var n,r=0;if(C(e)){for(n=e.length;r<n;r++)if(!1===t.call(e[r],r,e[r]))break}else for(r in e)if(!1===t.call(e[r],r,e[r]))break;return e},trim:function(e){return null==e?"":(e+"").replace(T,"")},makeArray:function(e,t){var n=t||[];return null!=e&&(C(Object(e))?w.merge(n,"string"==typeof e?[e]:e):s.call(n,e)),n},inArray:function(e,t,n){return null==t?-1:u.call(t,e,n)},merge:function(e,t){for(var n=+t.length,r=0,i=e.length;r<n;r++)e[i++]=t[r];return e.length=i,e},grep:function(e,t,n){for(var r,i=[],o=0,a=e.length,s=!n;o<a;o++)(r=!t(e[o],o))!==s&&i.push(e[o]);return i},map:function(e,t,n){var r,i,o=0,s=[];if(C(e))for(r=e.length;o<r;o++)null!=(i=t(e[o],o,n))&&s.push(i);else for(o in e)null!=(i=t(e[o],o,n))&&s.push(i);return a.apply([],s)},guid:1,support:h}),"function"==typeof Symbol&&(w.fn[Symbol.iterator]=n[Symbol.iterator]),w.each("Boolean Number String Function Array Date RegExp Object Error Symbol".split(" "),function(e,t){l["[object "+t+"]"]=t.toLowerCase()});function C(e){var t=!!e&&"length"in e&&e.length,n=x(e);return!g(e)&&!y(e)&&("array"===n||0===t||"number"==typeof t&&t>0&&t-1 in e)}var E=function(e){var t,n,r,i,o,a,s,u,l,c,f,p,d,h,g,y,v,m,x,b="sizzle"+1*new Date,w=e.document,T=0,C=0,E=ae(),k=ae(),S=ae(),D=function(e,t){return e===t&&(f=!0),0},N={}.hasOwnProperty,A=[],j=A.pop,q=A.push,L=A.push,H=A.slice,O=function(e,t){for(var n=0,r=e.length;n<r;n++)if(e[n]===t)return n;return-1},P="checked|selected|async|autofocus|autoplay|controls|defer|disabled|hidden|ismap|loop|multiple|open|readonly|required|scoped",M="[\\x20\\t\\r\\n\\f]",R="(?:\\\\.|[\\w-]|[^\0-\\xa0])+",I="\\["+M+"*("+R+")(?:"+M+"*([*^$|!~]?=)"+M+"*(?:'((?:\\\\.|[^\\\\'])*)'|\"((?:\\\\.|[^\\\\\"])*)\"|("+R+"))|)"+M+"*\\]",W=":("+R+")(?:\\((('((?:\\\\.|[^\\\\'])*)'|\"((?:\\\\.|[^\\\\\"])*)\")|((?:\\\\.|[^\\\\()[\\]]|"+I+")*)|.*)\\)|)",$=new RegExp(M+"+","g"),B=new RegExp("^"+M+"+|((?:^|[^\\\\])(?:\\\\.)*)"+M+"+$","g"),F=new RegExp("^"+M+"*,"+M+"*"),_=new RegExp("^"+M+"*([>+~]|"+M+")"+M+"*"),z=new RegExp("="+M+"*([^\\]'\"]*?)"+M+"*\\]","g"),X=new RegExp(W),U=new RegExp("^"+R+"$"),V={ID:new RegExp("^#("+R+")"),CLASS:new RegExp("^\\.("+R+")"),TAG:new RegExp("^("+R+"|[*])"),ATTR:new RegExp("^"+I),PSEUDO:new RegExp("^"+W),CHILD:new RegExp("^:(only|first|last|nth|nth-last)-(child|of-type)(?:\\("+M+"*(even|odd|(([+-]|)(\\d*)n|)"+M+"*(?:([+-]|)"+M+"*(\\d+)|))"+M+"*\\)|)","i"),bool:new RegExp("^(?:"+P+")$","i"),needsContext:new RegExp("^"+M+"*[>+~]|:(even|odd|eq|gt|lt|nth|first|last)(?:\\("+M+"*((?:-\\d)?\\d*)"+M+"*\\)|)(?=[^-]|$)","i")},G=/^(?:input|select|textarea|button)$/i,Y=/^h\d$/i,Q=/^[^{]+\{\s*\[native \w/,J=/^(?:#([\w-]+)|(\w+)|\.([\w-]+))$/,K=/[+~]/,Z=new RegExp("\\\\([\\da-f]{1,6}"+M+"?|("+M+")|.)","ig"),ee=function(e,t,n){var r="0x"+t-65536;return r!==r||n?t:r<0?String.fromCharCode(r+65536):String.fromCharCode(r>>10|55296,1023&r|56320)},te=/([\0-\x1f\x7f]|^-?\d)|^-$|[^\0-\x1f\x7f-\uFFFF\w-]/g,ne=function(e,t){return t?"\0"===e?"\ufffd":e.slice(0,-1)+"\\"+e.charCodeAt(e.length-1).toString(16)+" ":"\\"+e},re=function(){p()},ie=me(function(e){return!0===e.disabled&&("form"in e||"label"in e)},{dir:"parentNode",next:"legend"});try{L.apply(A=H.call(w.childNodes),w.childNodes),A[w.childNodes.length].nodeType}catch(e){L={apply:A.length?function(e,t){q.apply(e,H.call(t))}:function(e,t){var n=e.length,r=0;while(e[n++]=t[r++]);e.length=n-1}}}function oe(e,t,r,i){var o,s,l,c,f,h,v,m=t&&t.ownerDocument,T=t?t.nodeType:9;if(r=r||[],"string"!=typeof e||!e||1!==T&&9!==T&&11!==T)return r;if(!i&&((t?t.ownerDocument||t:w)!==d&&p(t),t=t||d,g)){if(11!==T&&(f=J.exec(e)))if(o=f[1]){if(9===T){if(!(l=t.getElementById(o)))return r;if(l.id===o)return r.push(l),r}else if(m&&(l=m.getElementById(o))&&x(t,l)&&l.id===o)return r.push(l),r}else{if(f[2])return L.apply(r,t.getElementsByTagName(e)),r;if((o=f[3])&&n.getElementsByClassName&&t.getElementsByClassName)return L.apply(r,t.getElementsByClassName(o)),r}if(n.qsa&&!S[e+" "]&&(!y||!y.test(e))){if(1!==T)m=t,v=e;else if("object"!==t.nodeName.toLowerCase()){(c=t.getAttribute("id"))?c=c.replace(te,ne):t.setAttribute("id",c=b),s=(h=a(e)).length;while(s--)h[s]="#"+c+" "+ve(h[s]);v=h.join(","),m=K.test(e)&&ge(t.parentNode)||t}if(v)try{return L.apply(r,m.querySelectorAll(v)),r}catch(e){}finally{c===b&&t.removeAttribute("id")}}}return u(e.replace(B,"$1"),t,r,i)}function ae(){var e=[];function t(n,i){return e.push(n+" ")>r.cacheLength&&delete t[e.shift()],t[n+" "]=i}return t}function se(e){return e[b]=!0,e}function ue(e){var t=d.createElement("fieldset");try{return!!e(t)}catch(e){return!1}finally{t.parentNode&&t.parentNode.removeChild(t),t=null}}function le(e,t){var n=e.split("|"),i=n.length;while(i--)r.attrHandle[n[i]]=t}function ce(e,t){var n=t&&e,r=n&&1===e.nodeType&&1===t.nodeType&&e.sourceIndex-t.sourceIndex;if(r)return r;if(n)while(n=n.nextSibling)if(n===t)return-1;return e?1:-1}function fe(e){return function(t){return"input"===t.nodeName.toLowerCase()&&t.type===e}}function pe(e){return function(t){var n=t.nodeName.toLowerCase();return("input"===n||"button"===n)&&t.type===e}}function de(e){return function(t){return"form"in t?t.parentNode&&!1===t.disabled?"label"in t?"label"in t.parentNode?t.parentNode.disabled===e:t.disabled===e:t.isDisabled===e||t.isDisabled!==!e&&ie(t)===e:t.disabled===e:"label"in t&&t.disabled===e}}function he(e){return se(function(t){return t=+t,se(function(n,r){var i,o=e([],n.length,t),a=o.length;while(a--)n[i=o[a]]&&(n[i]=!(r[i]=n[i]))})})}function ge(e){return e&&"undefined"!=typeof e.getElementsByTagName&&e}n=oe.support={},o=oe.isXML=function(e){var t=e&&(e.ownerDocument||e).documentElement;return!!t&&"HTML"!==t.nodeName},p=oe.setDocument=function(e){var t,i,a=e?e.ownerDocument||e:w;return a!==d&&9===a.nodeType&&a.documentElement?(d=a,h=d.documentElement,g=!o(d),w!==d&&(i=d.defaultView)&&i.top!==i&&(i.addEventListener?i.addEventListener("unload",re,!1):i.attachEvent&&i.attachEvent("onunload",re)),n.attributes=ue(function(e){return e.className="i",!e.getAttribute("className")}),n.getElementsByTagName=ue(function(e){return e.appendChild(d.createComment("")),!e.getElementsByTagName("*").length}),n.getElementsByClassName=Q.test(d.getElementsByClassName),n.getById=ue(function(e){return h.appendChild(e).id=b,!d.getElementsByName||!d.getElementsByName(b).length}),n.getById?(r.filter.ID=function(e){var t=e.replace(Z,ee);return function(e){return e.getAttribute("id")===t}},r.find.ID=function(e,t){if("undefined"!=typeof t.getElementById&&g){var n=t.getElementById(e);return n?[n]:[]}}):(r.filter.ID=function(e){var t=e.replace(Z,ee);return function(e){var n="undefined"!=typeof e.getAttributeNode&&e.getAttributeNode("id");return n&&n.value===t}},r.find.ID=function(e,t){if("undefined"!=typeof t.getElementById&&g){var n,r,i,o=t.getElementById(e);if(o){if((n=o.getAttributeNode("id"))&&n.value===e)return[o];i=t.getElementsByName(e),r=0;while(o=i[r++])if((n=o.getAttributeNode("id"))&&n.value===e)return[o]}return[]}}),r.find.TAG=n.getElementsByTagName?function(e,t){return"undefined"!=typeof t.getElementsByTagName?t.getElementsByTagName(e):n.qsa?t.querySelectorAll(e):void 0}:function(e,t){var n,r=[],i=0,o=t.getElementsByTagName(e);if("*"===e){while(n=o[i++])1===n.nodeType&&r.push(n);return r}return o},r.find.CLASS=n.getElementsByClassName&&function(e,t){if("undefined"!=typeof t.getElementsByClassName&&g)return t.getElementsByClassName(e)},v=[],y=[],(n.qsa=Q.test(d.querySelectorAll))&&(ue(function(e){h.appendChild(e).innerHTML="<a id='"+b+"'></a><select id='"+b+"-\r\\' msallowcapture=''><option selected=''></option></select>",e.querySelectorAll("[msallowcapture^='']").length&&y.push("[*^$]="+M+"*(?:''|\"\")"),e.querySelectorAll("[selected]").length||y.push("\\["+M+"*(?:value|"+P+")"),e.querySelectorAll("[id~="+b+"-]").length||y.push("~="),e.querySelectorAll(":checked").length||y.push(":checked"),e.querySelectorAll("a#"+b+"+*").length||y.push(".#.+[+~]")}),ue(function(e){e.innerHTML="<a href='' disabled='disabled'></a><select disabled='disabled'><option/></select>";var t=d.createElement("input");t.setAttribute("type","hidden"),e.appendChild(t).setAttribute("name","D"),e.querySelectorAll("[name=d]").length&&y.push("name"+M+"*[*^$|!~]?="),2!==e.querySelectorAll(":enabled").length&&y.push(":enabled",":disabled"),h.appendChild(e).disabled=!0,2!==e.querySelectorAll(":disabled").length&&y.push(":enabled",":disabled"),e.querySelectorAll("*,:x"),y.push(",.*:")})),(n.matchesSelector=Q.test(m=h.matches||h.webkitMatchesSelector||h.mozMatchesSelector||h.oMatchesSelector||h.msMatchesSelector))&&ue(function(e){n.disconnectedMatch=m.call(e,"*"),m.call(e,"[s!='']:x"),v.push("!=",W)}),y=y.length&&new RegExp(y.join("|")),v=v.length&&new RegExp(v.join("|")),t=Q.test(h.compareDocumentPosition),x=t||Q.test(h.contains)?function(e,t){var n=9===e.nodeType?e.documentElement:e,r=t&&t.parentNode;return e===r||!(!r||1!==r.nodeType||!(n.contains?n.contains(r):e.compareDocumentPosition&&16&e.compareDocumentPosition(r)))}:function(e,t){if(t)while(t=t.parentNode)if(t===e)return!0;return!1},D=t?function(e,t){if(e===t)return f=!0,0;var r=!e.compareDocumentPosition-!t.compareDocumentPosition;return r||(1&(r=(e.ownerDocument||e)===(t.ownerDocument||t)?e.compareDocumentPosition(t):1)||!n.sortDetached&&t.compareDocumentPosition(e)===r?e===d||e.ownerDocument===w&&x(w,e)?-1:t===d||t.ownerDocument===w&&x(w,t)?1:c?O(c,e)-O(c,t):0:4&r?-1:1)}:function(e,t){if(e===t)return f=!0,0;var n,r=0,i=e.parentNode,o=t.parentNode,a=[e],s=[t];if(!i||!o)return e===d?-1:t===d?1:i?-1:o?1:c?O(c,e)-O(c,t):0;if(i===o)return ce(e,t);n=e;while(n=n.parentNode)a.unshift(n);n=t;while(n=n.parentNode)s.unshift(n);while(a[r]===s[r])r++;return r?ce(a[r],s[r]):a[r]===w?-1:s[r]===w?1:0},d):d},oe.matches=function(e,t){return oe(e,null,null,t)},oe.matchesSelector=function(e,t){if((e.ownerDocument||e)!==d&&p(e),t=t.replace(z,"='$1']"),n.matchesSelector&&g&&!S[t+" "]&&(!v||!v.test(t))&&(!y||!y.test(t)))try{var r=m.call(e,t);if(r||n.disconnectedMatch||e.document&&11!==e.document.nodeType)return r}catch(e){}return oe(t,d,null,[e]).length>0},oe.contains=function(e,t){return(e.ownerDocument||e)!==d&&p(e),x(e,t)},oe.attr=function(e,t){(e.ownerDocument||e)!==d&&p(e);var i=r.attrHandle[t.toLowerCase()],o=i&&N.call(r.attrHandle,t.toLowerCase())?i(e,t,!g):void 0;return void 0!==o?o:n.attributes||!g?e.getAttribute(t):(o=e.getAttributeNode(t))&&o.specified?o.value:null},oe.escape=function(e){return(e+"").replace(te,ne)},oe.error=function(e){throw new Error("Syntax error, unrecognized expression: "+e)},oe.uniqueSort=function(e){var t,r=[],i=0,o=0;if(f=!n.detectDuplicates,c=!n.sortStable&&e.slice(0),e.sort(D),f){while(t=e[o++])t===e[o]&&(i=r.push(o));while(i--)e.splice(r[i],1)}return c=null,e},i=oe.getText=function(e){var t,n="",r=0,o=e.nodeType;if(o){if(1===o||9===o||11===o){if("string"==typeof e.textContent)return e.textContent;for(e=e.firstChild;e;e=e.nextSibling)n+=i(e)}else if(3===o||4===o)return e.nodeValue}else while(t=e[r++])n+=i(t);return n},(r=oe.selectors={cacheLength:50,createPseudo:se,match:V,attrHandle:{},find:{},relative:{">":{dir:"parentNode",first:!0}," ":{dir:"parentNode"},"+":{dir:"previousSibling",first:!0},"~":{dir:"previousSibling"}},preFilter:{ATTR:function(e){return e[1]=e[1].replace(Z,ee),e[3]=(e[3]||e[4]||e[5]||"").replace(Z,ee),"~="===e[2]&&(e[3]=" "+e[3]+" "),e.slice(0,4)},CHILD:function(e){return e[1]=e[1].toLowerCase(),"nth"===e[1].slice(0,3)?(e[3]||oe.error(e[0]),e[4]=+(e[4]?e[5]+(e[6]||1):2*("even"===e[3]||"odd"===e[3])),e[5]=+(e[7]+e[8]||"odd"===e[3])):e[3]&&oe.error(e[0]),e},PSEUDO:function(e){var t,n=!e[6]&&e[2];return V.CHILD.test(e[0])?null:(e[3]?e[2]=e[4]||e[5]||"":n&&X.test(n)&&(t=a(n,!0))&&(t=n.indexOf(")",n.length-t)-n.length)&&(e[0]=e[0].slice(0,t),e[2]=n.slice(0,t)),e.slice(0,3))}},filter:{TAG:function(e){var t=e.replace(Z,ee).toLowerCase();return"*"===e?function(){return!0}:function(e){return e.nodeName&&e.nodeName.toLowerCase()===t}},CLASS:function(e){var t=E[e+" "];return t||(t=new RegExp("(^|"+M+")"+e+"("+M+"|$)"))&&E(e,function(e){return t.test("string"==typeof e.className&&e.className||"undefined"!=typeof e.getAttribute&&e.getAttribute("class")||"")})},ATTR:function(e,t,n){return function(r){var i=oe.attr(r,e);return null==i?"!="===t:!t||(i+="","="===t?i===n:"!="===t?i!==n:"^="===t?n&&0===i.indexOf(n):"*="===t?n&&i.indexOf(n)>-1:"$="===t?n&&i.slice(-n.length)===n:"~="===t?(" "+i.replace($," ")+" ").indexOf(n)>-1:"|="===t&&(i===n||i.slice(0,n.length+1)===n+"-"))}},CHILD:function(e,t,n,r,i){var o="nth"!==e.slice(0,3),a="last"!==e.slice(-4),s="of-type"===t;return 1===r&&0===i?function(e){return!!e.parentNode}:function(t,n,u){var l,c,f,p,d,h,g=o!==a?"nextSibling":"previousSibling",y=t.parentNode,v=s&&t.nodeName.toLowerCase(),m=!u&&!s,x=!1;if(y){if(o){while(g){p=t;while(p=p[g])if(s?p.nodeName.toLowerCase()===v:1===p.nodeType)return!1;h=g="only"===e&&!h&&"nextSibling"}return!0}if(h=[a?y.firstChild:y.lastChild],a&&m){x=(d=(l=(c=(f=(p=y)[b]||(p[b]={}))[p.uniqueID]||(f[p.uniqueID]={}))[e]||[])[0]===T&&l[1])&&l[2],p=d&&y.childNodes[d];while(p=++d&&p&&p[g]||(x=d=0)||h.pop())if(1===p.nodeType&&++x&&p===t){c[e]=[T,d,x];break}}else if(m&&(x=d=(l=(c=(f=(p=t)[b]||(p[b]={}))[p.uniqueID]||(f[p.uniqueID]={}))[e]||[])[0]===T&&l[1]),!1===x)while(p=++d&&p&&p[g]||(x=d=0)||h.pop())if((s?p.nodeName.toLowerCase()===v:1===p.nodeType)&&++x&&(m&&((c=(f=p[b]||(p[b]={}))[p.uniqueID]||(f[p.uniqueID]={}))[e]=[T,x]),p===t))break;return(x-=i)===r||x%r==0&&x/r>=0}}},PSEUDO:function(e,t){var n,i=r.pseudos[e]||r.setFilters[e.toLowerCase()]||oe.error("unsupported pseudo: "+e);return i[b]?i(t):i.length>1?(n=[e,e,"",t],r.setFilters.hasOwnProperty(e.toLowerCase())?se(function(e,n){var r,o=i(e,t),a=o.length;while(a--)e[r=O(e,o[a])]=!(n[r]=o[a])}):function(e){return i(e,0,n)}):i}},pseudos:{not:se(function(e){var t=[],n=[],r=s(e.replace(B,"$1"));return r[b]?se(function(e,t,n,i){var o,a=r(e,null,i,[]),s=e.length;while(s--)(o=a[s])&&(e[s]=!(t[s]=o))}):function(e,i,o){return t[0]=e,r(t,null,o,n),t[0]=null,!n.pop()}}),has:se(function(e){return function(t){return oe(e,t).length>0}}),contains:se(function(e){return e=e.replace(Z,ee),function(t){return(t.textContent||t.innerText||i(t)).indexOf(e)>-1}}),lang:se(function(e){return U.test(e||"")||oe.error("unsupported lang: "+e),e=e.replace(Z,ee).toLowerCase(),function(t){var n;do{if(n=g?t.lang:t.getAttribute("xml:lang")||t.getAttribute("lang"))return(n=n.toLowerCase())===e||0===n.indexOf(e+"-")}while((t=t.parentNode)&&1===t.nodeType);return!1}}),target:function(t){var n=e.location&&e.location.hash;return n&&n.slice(1)===t.id},root:function(e){return e===h},focus:function(e){return e===d.activeElement&&(!d.hasFocus||d.hasFocus())&&!!(e.type||e.href||~e.tabIndex)},enabled:de(!1),disabled:de(!0),checked:function(e){var t=e.nodeName.toLowerCase();return"input"===t&&!!e.checked||"option"===t&&!!e.selected},selected:function(e){return e.parentNode&&e.parentNode.selectedIndex,!0===e.selected},empty:function(e){for(e=e.firstChild;e;e=e.nextSibling)if(e.nodeType<6)return!1;return!0},parent:function(e){return!r.pseudos.empty(e)},header:function(e){return Y.test(e.nodeName)},input:function(e){return G.test(e.nodeName)},button:function(e){var t=e.nodeName.toLowerCase();return"input"===t&&"button"===e.type||"button"===t},text:function(e){var t;return"input"===e.nodeName.toLowerCase()&&"text"===e.type&&(null==(t=e.getAttribute("type"))||"text"===t.toLowerCase())},first:he(function(){return[0]}),last:he(function(e,t){return[t-1]}),eq:he(function(e,t,n){return[n<0?n+t:n]}),even:he(function(e,t){for(var n=0;n<t;n+=2)e.push(n);return e}),odd:he(function(e,t){for(var n=1;n<t;n+=2)e.push(n);return e}),lt:he(function(e,t,n){for(var r=n<0?n+t:n;--r>=0;)e.push(r);return e}),gt:he(function(e,t,n){for(var r=n<0?n+t:n;++r<t;)e.push(r);return e})}}).pseudos.nth=r.pseudos.eq;for(t in{radio:!0,checkbox:!0,file:!0,password:!0,image:!0})r.pseudos[t]=fe(t);for(t in{submit:!0,reset:!0})r.pseudos[t]=pe(t);function ye(){}ye.prototype=r.filters=r.pseudos,r.setFilters=new ye,a=oe.tokenize=function(e,t){var n,i,o,a,s,u,l,c=k[e+" "];if(c)return t?0:c.slice(0);s=e,u=[],l=r.preFilter;while(s){n&&!(i=F.exec(s))||(i&&(s=s.slice(i[0].length)||s),u.push(o=[])),n=!1,(i=_.exec(s))&&(n=i.shift(),o.push({value:n,type:i[0].replace(B," ")}),s=s.slice(n.length));for(a in r.filter)!(i=V[a].exec(s))||l[a]&&!(i=l[a](i))||(n=i.shift(),o.push({value:n,type:a,matches:i}),s=s.slice(n.length));if(!n)break}return t?s.length:s?oe.error(e):k(e,u).slice(0)};function ve(e){for(var t=0,n=e.length,r="";t<n;t++)r+=e[t].value;return r}function me(e,t,n){var r=t.dir,i=t.next,o=i||r,a=n&&"parentNode"===o,s=C++;return t.first?function(t,n,i){while(t=t[r])if(1===t.nodeType||a)return e(t,n,i);return!1}:function(t,n,u){var l,c,f,p=[T,s];if(u){while(t=t[r])if((1===t.nodeType||a)&&e(t,n,u))return!0}else while(t=t[r])if(1===t.nodeType||a)if(f=t[b]||(t[b]={}),c=f[t.uniqueID]||(f[t.uniqueID]={}),i&&i===t.nodeName.toLowerCase())t=t[r]||t;else{if((l=c[o])&&l[0]===T&&l[1]===s)return p[2]=l[2];if(c[o]=p,p[2]=e(t,n,u))return!0}return!1}}function xe(e){return e.length>1?function(t,n,r){var i=e.length;while(i--)if(!e[i](t,n,r))return!1;return!0}:e[0]}function be(e,t,n){for(var r=0,i=t.length;r<i;r++)oe(e,t[r],n);return n}function we(e,t,n,r,i){for(var o,a=[],s=0,u=e.length,l=null!=t;s<u;s++)(o=e[s])&&(n&&!n(o,r,i)||(a.push(o),l&&t.push(s)));return a}function Te(e,t,n,r,i,o){return r&&!r[b]&&(r=Te(r)),i&&!i[b]&&(i=Te(i,o)),se(function(o,a,s,u){var l,c,f,p=[],d=[],h=a.length,g=o||be(t||"*",s.nodeType?[s]:s,[]),y=!e||!o&&t?g:we(g,p,e,s,u),v=n?i||(o?e:h||r)?[]:a:y;if(n&&n(y,v,s,u),r){l=we(v,d),r(l,[],s,u),c=l.length;while(c--)(f=l[c])&&(v[d[c]]=!(y[d[c]]=f))}if(o){if(i||e){if(i){l=[],c=v.length;while(c--)(f=v[c])&&l.push(y[c]=f);i(null,v=[],l,u)}c=v.length;while(c--)(f=v[c])&&(l=i?O(o,f):p[c])>-1&&(o[l]=!(a[l]=f))}}else v=we(v===a?v.splice(h,v.length):v),i?i(null,a,v,u):L.apply(a,v)})}function Ce(e){for(var t,n,i,o=e.length,a=r.relative[e[0].type],s=a||r.relative[" "],u=a?1:0,c=me(function(e){return e===t},s,!0),f=me(function(e){return O(t,e)>-1},s,!0),p=[function(e,n,r){var i=!a&&(r||n!==l)||((t=n).nodeType?c(e,n,r):f(e,n,r));return t=null,i}];u<o;u++)if(n=r.relative[e[u].type])p=[me(xe(p),n)];else{if((n=r.filter[e[u].type].apply(null,e[u].matches))[b]){for(i=++u;i<o;i++)if(r.relative[e[i].type])break;return Te(u>1&&xe(p),u>1&&ve(e.slice(0,u-1).concat({value:" "===e[u-2].type?"*":""})).replace(B,"$1"),n,u<i&&Ce(e.slice(u,i)),i<o&&Ce(e=e.slice(i)),i<o&&ve(e))}p.push(n)}return xe(p)}function Ee(e,t){var n=t.length>0,i=e.length>0,o=function(o,a,s,u,c){var f,h,y,v=0,m="0",x=o&&[],b=[],w=l,C=o||i&&r.find.TAG("*",c),E=T+=null==w?1:Math.random()||.1,k=C.length;for(c&&(l=a===d||a||c);m!==k&&null!=(f=C[m]);m++){if(i&&f){h=0,a||f.ownerDocument===d||(p(f),s=!g);while(y=e[h++])if(y(f,a||d,s)){u.push(f);break}c&&(T=E)}n&&((f=!y&&f)&&v--,o&&x.push(f))}if(v+=m,n&&m!==v){h=0;while(y=t[h++])y(x,b,a,s);if(o){if(v>0)while(m--)x[m]||b[m]||(b[m]=j.call(u));b=we(b)}L.apply(u,b),c&&!o&&b.length>0&&v+t.length>1&&oe.uniqueSort(u)}return c&&(T=E,l=w),x};return n?se(o):o}return s=oe.compile=function(e,t){var n,r=[],i=[],o=S[e+" "];if(!o){t||(t=a(e)),n=t.length;while(n--)(o=Ce(t[n]))[b]?r.push(o):i.push(o);(o=S(e,Ee(i,r))).selector=e}return o},u=oe.select=function(e,t,n,i){var o,u,l,c,f,p="function"==typeof e&&e,d=!i&&a(e=p.selector||e);if(n=n||[],1===d.length){if((u=d[0]=d[0].slice(0)).length>2&&"ID"===(l=u[0]).type&&9===t.nodeType&&g&&r.relative[u[1].type]){if(!(t=(r.find.ID(l.matches[0].replace(Z,ee),t)||[])[0]))return n;p&&(t=t.parentNode),e=e.slice(u.shift().value.length)}o=V.needsContext.test(e)?0:u.length;while(o--){if(l=u[o],r.relative[c=l.type])break;if((f=r.find[c])&&(i=f(l.matches[0].replace(Z,ee),K.test(u[0].type)&&ge(t.parentNode)||t))){if(u.splice(o,1),!(e=i.length&&ve(u)))return L.apply(n,i),n;break}}}return(p||s(e,d))(i,t,!g,n,!t||K.test(e)&&ge(t.parentNode)||t),n},n.sortStable=b.split("").sort(D).join("")===b,n.detectDuplicates=!!f,p(),n.sortDetached=ue(function(e){return 1&e.compareDocumentPosition(d.createElement("fieldset"))}),ue(function(e){return e.innerHTML="<a href='#'></a>","#"===e.firstChild.getAttribute("href")})||le("type|href|height|width",function(e,t,n){if(!n)return e.getAttribute(t,"type"===t.toLowerCase()?1:2)}),n.attributes&&ue(function(e){return e.innerHTML="<input/>",e.firstChild.setAttribute("value",""),""===e.firstChild.getAttribute("value")})||le("value",function(e,t,n){if(!n&&"input"===e.nodeName.toLowerCase())return e.defaultValue}),ue(function(e){return null==e.getAttribute("disabled")})||le(P,function(e,t,n){var r;if(!n)return!0===e[t]?t.toLowerCase():(r=e.getAttributeNode(t))&&r.specified?r.value:null}),oe}(e);w.find=E,w.expr=E.selectors,w.expr[":"]=w.expr.pseudos,w.uniqueSort=w.unique=E.uniqueSort,w.text=E.getText,w.isXMLDoc=E.isXML,w.contains=E.contains,w.escapeSelector=E.escape;var k=function(e,t,n){var r=[],i=void 0!==n;while((e=e[t])&&9!==e.nodeType)if(1===e.nodeType){if(i&&w(e).is(n))break;r.push(e)}return r},S=function(e,t){for(var n=[];e;e=e.nextSibling)1===e.nodeType&&e!==t&&n.push(e);return n},D=w.expr.match.needsContext;function N(e,t){return e.nodeName&&e.nodeName.toLowerCase()===t.toLowerCase()}var A=/^<([a-z][^\/\0>:\x20\t\r\n\f]*)[\x20\t\r\n\f]*\/?>(?:<\/\1>|)$/i;function j(e,t,n){return g(t)?w.grep(e,function(e,r){return!!t.call(e,r,e)!==n}):t.nodeType?w.grep(e,function(e){return e===t!==n}):"string"!=typeof t?w.grep(e,function(e){return u.call(t,e)>-1!==n}):w.filter(t,e,n)}w.filter=function(e,t,n){var r=t[0];return n&&(e=":not("+e+")"),1===t.length&&1===r.nodeType?w.find.matchesSelector(r,e)?[r]:[]:w.find.matches(e,w.grep(t,function(e){return 1===e.nodeType}))},w.fn.extend({find:function(e){var t,n,r=this.length,i=this;if("string"!=typeof e)return this.pushStack(w(e).filter(function(){for(t=0;t<r;t++)if(w.contains(i[t],this))return!0}));for(n=this.pushStack([]),t=0;t<r;t++)w.find(e,i[t],n);return r>1?w.uniqueSort(n):n},filter:function(e){return this.pushStack(j(this,e||[],!1))},not:function(e){return this.pushStack(j(this,e||[],!0))},is:function(e){return!!j(this,"string"==typeof e&&D.test(e)?w(e):e||[],!1).length}});var q,L=/^(?:\s*(<[\w\W]+>)[^>]*|#([\w-]+))$/;(w.fn.init=function(e,t,n){var i,o;if(!e)return this;if(n=n||q,"string"==typeof e){if(!(i="<"===e[0]&&">"===e[e.length-1]&&e.length>=3?[null,e,null]:L.exec(e))||!i[1]&&t)return!t||t.jquery?(t||n).find(e):this.constructor(t).find(e);if(i[1]){if(t=t instanceof w?t[0]:t,w.merge(this,w.parseHTML(i[1],t&&t.nodeType?t.ownerDocument||t:r,!0)),A.test(i[1])&&w.isPlainObject(t))for(i in t)g(this[i])?this[i](t[i]):this.attr(i,t[i]);return this}return(o=r.getElementById(i[2]))&&(this[0]=o,this.length=1),this}return e.nodeType?(this[0]=e,this.length=1,this):g(e)?void 0!==n.ready?n.ready(e):e(w):w.makeArray(e,this)}).prototype=w.fn,q=w(r);var H=/^(?:parents|prev(?:Until|All))/,O={children:!0,contents:!0,next:!0,prev:!0};w.fn.extend({has:function(e){var t=w(e,this),n=t.length;return this.filter(function(){for(var e=0;e<n;e++)if(w.contains(this,t[e]))return!0})},closest:function(e,t){var n,r=0,i=this.length,o=[],a="string"!=typeof e&&w(e);if(!D.test(e))for(;r<i;r++)for(n=this[r];n&&n!==t;n=n.parentNode)if(n.nodeType<11&&(a?a.index(n)>-1:1===n.nodeType&&w.find.matchesSelector(n,e))){o.push(n);break}return this.pushStack(o.length>1?w.uniqueSort(o):o)},index:function(e){return e?"string"==typeof e?u.call(w(e),this[0]):u.call(this,e.jquery?e[0]:e):this[0]&&this[0].parentNode?this.first().prevAll().length:-1},add:function(e,t){return this.pushStack(w.uniqueSort(w.merge(this.get(),w(e,t))))},addBack:function(e){return this.add(null==e?this.prevObject:this.prevObject.filter(e))}});function P(e,t){while((e=e[t])&&1!==e.nodeType);return e}w.each({parent:function(e){var t=e.parentNode;return t&&11!==t.nodeType?t:null},parents:function(e){return k(e,"parentNode")},parentsUntil:function(e,t,n){return k(e,"parentNode",n)},next:function(e){return P(e,"nextSibling")},prev:function(e){return P(e,"previousSibling")},nextAll:function(e){return k(e,"nextSibling")},prevAll:function(e){return k(e,"previousSibling")},nextUntil:function(e,t,n){return k(e,"nextSibling",n)},prevUntil:function(e,t,n){return k(e,"previousSibling",n)},siblings:function(e){return S((e.parentNode||{}).firstChild,e)},children:function(e){return S(e.firstChild)},contents:function(e){return N(e,"iframe")?e.contentDocument:(N(e,"template")&&(e=e.content||e),w.merge([],e.childNodes))}},function(e,t){w.fn[e]=function(n,r){var i=w.map(this,t,n);return"Until"!==e.slice(-5)&&(r=n),r&&"string"==typeof r&&(i=w.filter(r,i)),this.length>1&&(O[e]||w.uniqueSort(i),H.test(e)&&i.reverse()),this.pushStack(i)}});var M=/[^\x20\t\r\n\f]+/g;function R(e){var t={};return w.each(e.match(M)||[],function(e,n){t[n]=!0}),t}w.Callbacks=function(e){e="string"==typeof e?R(e):w.extend({},e);var t,n,r,i,o=[],a=[],s=-1,u=function(){for(i=i||e.once,r=t=!0;a.length;s=-1){n=a.shift();while(++s<o.length)!1===o[s].apply(n[0],n[1])&&e.stopOnFalse&&(s=o.length,n=!1)}e.memory||(n=!1),t=!1,i&&(o=n?[]:"")},l={add:function(){return o&&(n&&!t&&(s=o.length-1,a.push(n)),function t(n){w.each(n,function(n,r){g(r)?e.unique&&l.has(r)||o.push(r):r&&r.length&&"string"!==x(r)&&t(r)})}(arguments),n&&!t&&u()),this},remove:function(){return w.each(arguments,function(e,t){var n;while((n=w.inArray(t,o,n))>-1)o.splice(n,1),n<=s&&s--}),this},has:function(e){return e?w.inArray(e,o)>-1:o.length>0},empty:function(){return o&&(o=[]),this},disable:function(){return i=a=[],o=n="",this},disabled:function(){return!o},lock:function(){return i=a=[],n||t||(o=n=""),this},locked:function(){return!!i},fireWith:function(e,n){return i||(n=[e,(n=n||[]).slice?n.slice():n],a.push(n),t||u()),this},fire:function(){return l.fireWith(this,arguments),this},fired:function(){return!!r}};return l};function I(e){return e}function W(e){throw e}function $(e,t,n,r){var i;try{e&&g(i=e.promise)?i.call(e).done(t).fail(n):e&&g(i=e.then)?i.call(e,t,n):t.apply(void 0,[e].slice(r))}catch(e){n.apply(void 0,[e])}}w.extend({Deferred:function(t){var n=[["notify","progress",w.Callbacks("memory"),w.Callbacks("memory"),2],["resolve","done",w.Callbacks("once memory"),w.Callbacks("once memory"),0,"resolved"],["reject","fail",w.Callbacks("once memory"),w.Callbacks("once memory"),1,"rejected"]],r="pending",i={state:function(){return r},always:function(){return o.done(arguments).fail(arguments),this},"catch":function(e){return i.then(null,e)},pipe:function(){var e=arguments;return w.Deferred(function(t){w.each(n,function(n,r){var i=g(e[r[4]])&&e[r[4]];o[r[1]](function(){var e=i&&i.apply(this,arguments);e&&g(e.promise)?e.promise().progress(t.notify).done(t.resolve).fail(t.reject):t[r[0]+"With"](this,i?[e]:arguments)})}),e=null}).promise()},then:function(t,r,i){var o=0;function a(t,n,r,i){return function(){var s=this,u=arguments,l=function(){var e,l;if(!(t<o)){if((e=r.apply(s,u))===n.promise())throw new TypeError("Thenable self-resolution");l=e&&("object"==typeof e||"function"==typeof e)&&e.then,g(l)?i?l.call(e,a(o,n,I,i),a(o,n,W,i)):(o++,l.call(e,a(o,n,I,i),a(o,n,W,i),a(o,n,I,n.notifyWith))):(r!==I&&(s=void 0,u=[e]),(i||n.resolveWith)(s,u))}},c=i?l:function(){try{l()}catch(e){w.Deferred.exceptionHook&&w.Deferred.exceptionHook(e,c.stackTrace),t+1>=o&&(r!==W&&(s=void 0,u=[e]),n.rejectWith(s,u))}};t?c():(w.Deferred.getStackHook&&(c.stackTrace=w.Deferred.getStackHook()),e.setTimeout(c))}}return w.Deferred(function(e){n[0][3].add(a(0,e,g(i)?i:I,e.notifyWith)),n[1][3].add(a(0,e,g(t)?t:I)),n[2][3].add(a(0,e,g(r)?r:W))}).promise()},promise:function(e){return null!=e?w.extend(e,i):i}},o={};return w.each(n,function(e,t){var a=t[2],s=t[5];i[t[1]]=a.add,s&&a.add(function(){r=s},n[3-e][2].disable,n[3-e][3].disable,n[0][2].lock,n[0][3].lock),a.add(t[3].fire),o[t[0]]=function(){return o[t[0]+"With"](this===o?void 0:this,arguments),this},o[t[0]+"With"]=a.fireWith}),i.promise(o),t&&t.call(o,o),o},when:function(e){var t=arguments.length,n=t,r=Array(n),i=o.call(arguments),a=w.Deferred(),s=function(e){return function(n){r[e]=this,i[e]=arguments.length>1?o.call(arguments):n,--t||a.resolveWith(r,i)}};if(t<=1&&($(e,a.done(s(n)).resolve,a.reject,!t),"pending"===a.state()||g(i[n]&&i[n].then)))return a.then();while(n--)$(i[n],s(n),a.reject);return a.promise()}});var B=/^(Eval|Internal|Range|Reference|Syntax|Type|URI)Error$/;w.Deferred.exceptionHook=function(t,n){e.console&&e.console.warn&&t&&B.test(t.name)&&e.console.warn("jQuery.Deferred exception: "+t.message,t.stack,n)},w.readyException=function(t){e.setTimeout(function(){throw t})};var F=w.Deferred();w.fn.ready=function(e){return F.then(e)["catch"](function(e){w.readyException(e)}),this},w.extend({isReady:!1,readyWait:1,ready:function(e){(!0===e?--w.readyWait:w.isReady)||(w.isReady=!0,!0!==e&&--w.readyWait>0||F.resolveWith(r,[w]))}}),w.ready.then=F.then;function _(){r.removeEventListener("DOMContentLoaded",_),e.removeEventListener("load",_),w.ready()}"complete"===r.readyState||"loading"!==r.readyState&&!r.documentElement.doScroll?e.setTimeout(w.ready):(r.addEventListener("DOMContentLoaded",_),e.addEventListener("load",_));var z=function(e,t,n,r,i,o,a){var s=0,u=e.length,l=null==n;if("object"===x(n)){i=!0;for(s in n)z(e,t,s,n[s],!0,o,a)}else if(void 0!==r&&(i=!0,g(r)||(a=!0),l&&(a?(t.call(e,r),t=null):(l=t,t=function(e,t,n){return l.call(w(e),n)})),t))for(;s<u;s++)t(e[s],n,a?r:r.call(e[s],s,t(e[s],n)));return i?e:l?t.call(e):u?t(e[0],n):o},X=/^-ms-/,U=/-([a-z])/g;function V(e,t){return t.toUpperCase()}function G(e){return e.replace(X,"ms-").replace(U,V)}var Y=function(e){return 1===e.nodeType||9===e.nodeType||!+e.nodeType};function Q(){this.expando=w.expando+Q.uid++}Q.uid=1,Q.prototype={cache:function(e){var t=e[this.expando];return t||(t={},Y(e)&&(e.nodeType?e[this.expando]=t:Object.defineProperty(e,this.expando,{value:t,configurable:!0}))),t},set:function(e,t,n){var r,i=this.cache(e);if("string"==typeof t)i[G(t)]=n;else for(r in t)i[G(r)]=t[r];return i},get:function(e,t){return void 0===t?this.cache(e):e[this.expando]&&e[this.expando][G(t)]},access:function(e,t,n){return void 0===t||t&&"string"==typeof t&&void 0===n?this.get(e,t):(this.set(e,t,n),void 0!==n?n:t)},remove:function(e,t){var n,r=e[this.expando];if(void 0!==r){if(void 0!==t){n=(t=Array.isArray(t)?t.map(G):(t=G(t))in r?[t]:t.match(M)||[]).length;while(n--)delete r[t[n]]}(void 0===t||w.isEmptyObject(r))&&(e.nodeType?e[this.expando]=void 0:delete e[this.expando])}},hasData:function(e){var t=e[this.expando];return void 0!==t&&!w.isEmptyObject(t)}};var J=new Q,K=new Q,Z=/^(?:\{[\w\W]*\}|\[[\w\W]*\])$/,ee=/[A-Z]/g;function te(e){return"true"===e||"false"!==e&&("null"===e?null:e===+e+""?+e:Z.test(e)?JSON.parse(e):e)}function ne(e,t,n){var r;if(void 0===n&&1===e.nodeType)if(r="data-"+t.replace(ee,"-$&").toLowerCase(),"string"==typeof(n=e.getAttribute(r))){try{n=te(n)}catch(e){}K.set(e,t,n)}else n=void 0;return n}w.extend({hasData:function(e){return K.hasData(e)||J.hasData(e)},data:function(e,t,n){return K.access(e,t,n)},removeData:function(e,t){K.remove(e,t)},_data:function(e,t,n){return J.access(e,t,n)},_removeData:function(e,t){J.remove(e,t)}}),w.fn.extend({data:function(e,t){var n,r,i,o=this[0],a=o&&o.attributes;if(void 0===e){if(this.length&&(i=K.get(o),1===o.nodeType&&!J.get(o,"hasDataAttrs"))){n=a.length;while(n--)a[n]&&0===(r=a[n].name).indexOf("data-")&&(r=G(r.slice(5)),ne(o,r,i[r]));J.set(o,"hasDataAttrs",!0)}return i}return"object"==typeof e?this.each(function(){K.set(this,e)}):z(this,function(t){var n;if(o&&void 0===t){if(void 0!==(n=K.get(o,e)))return n;if(void 0!==(n=ne(o,e)))return n}else this.each(function(){K.set(this,e,t)})},null,t,arguments.length>1,null,!0)},removeData:function(e){return this.each(function(){K.remove(this,e)})}}),w.extend({queue:function(e,t,n){var r;if(e)return t=(t||"fx")+"queue",r=J.get(e,t),n&&(!r||Array.isArray(n)?r=J.access(e,t,w.makeArray(n)):r.push(n)),r||[]},dequeue:function(e,t){t=t||"fx";var n=w.queue(e,t),r=n.length,i=n.shift(),o=w._queueHooks(e,t),a=function(){w.dequeue(e,t)};"inprogress"===i&&(i=n.shift(),r--),i&&("fx"===t&&n.unshift("inprogress"),delete o.stop,i.call(e,a,o)),!r&&o&&o.empty.fire()},_queueHooks:function(e,t){var n=t+"queueHooks";return J.get(e,n)||J.access(e,n,{empty:w.Callbacks("once memory").add(function(){J.remove(e,[t+"queue",n])})})}}),w.fn.extend({queue:function(e,t){var n=2;return"string"!=typeof e&&(t=e,e="fx",n--),arguments.length<n?w.queue(this[0],e):void 0===t?this:this.each(function(){var n=w.queue(this,e,t);w._queueHooks(this,e),"fx"===e&&"inprogress"!==n[0]&&w.dequeue(this,e)})},dequeue:function(e){return this.each(function(){w.dequeue(this,e)})},clearQueue:function(e){return this.queue(e||"fx",[])},promise:function(e,t){var n,r=1,i=w.Deferred(),o=this,a=this.length,s=function(){--r||i.resolveWith(o,[o])};"string"!=typeof e&&(t=e,e=void 0),e=e||"fx";while(a--)(n=J.get(o[a],e+"queueHooks"))&&n.empty&&(r++,n.empty.add(s));return s(),i.promise(t)}});var re=/[+-]?(?:\d*\.|)\d+(?:[eE][+-]?\d+|)/.source,ie=new RegExp("^(?:([+-])=|)("+re+")([a-z%]*)$","i"),oe=["Top","Right","Bottom","Left"],ae=function(e,t){return"none"===(e=t||e).style.display||""===e.style.display&&w.contains(e.ownerDocument,e)&&"none"===w.css(e,"display")},se=function(e,t,n,r){var i,o,a={};for(o in t)a[o]=e.style[o],e.style[o]=t[o];i=n.apply(e,r||[]);for(o in t)e.style[o]=a[o];return i};function ue(e,t,n,r){var i,o,a=20,s=r?function(){return r.cur()}:function(){return w.css(e,t,"")},u=s(),l=n&&n[3]||(w.cssNumber[t]?"":"px"),c=(w.cssNumber[t]||"px"!==l&&+u)&&ie.exec(w.css(e,t));if(c&&c[3]!==l){u/=2,l=l||c[3],c=+u||1;while(a--)w.style(e,t,c+l),(1-o)*(1-(o=s()/u||.5))<=0&&(a=0),c/=o;c*=2,w.style(e,t,c+l),n=n||[]}return n&&(c=+c||+u||0,i=n[1]?c+(n[1]+1)*n[2]:+n[2],r&&(r.unit=l,r.start=c,r.end=i)),i}var le={};function ce(e){var t,n=e.ownerDocument,r=e.nodeName,i=le[r];return i||(t=n.body.appendChild(n.createElement(r)),i=w.css(t,"display"),t.parentNode.removeChild(t),"none"===i&&(i="block"),le[r]=i,i)}function fe(e,t){for(var n,r,i=[],o=0,a=e.length;o<a;o++)(r=e[o]).style&&(n=r.style.display,t?("none"===n&&(i[o]=J.get(r,"display")||null,i[o]||(r.style.display="")),""===r.style.display&&ae(r)&&(i[o]=ce(r))):"none"!==n&&(i[o]="none",J.set(r,"display",n)));for(o=0;o<a;o++)null!=i[o]&&(e[o].style.display=i[o]);return e}w.fn.extend({show:function(){return fe(this,!0)},hide:function(){return fe(this)},toggle:function(e){return"boolean"==typeof e?e?this.show():this.hide():this.each(function(){ae(this)?w(this).show():w(this).hide()})}});var pe=/^(?:checkbox|radio)$/i,de=/<([a-z][^\/\0>\x20\t\r\n\f]+)/i,he=/^$|^module$|\/(?:java|ecma)script/i,ge={option:[1,"<select multiple='multiple'>","</select>"],thead:[1,"<table>","</table>"],col:[2,"<table><colgroup>","</colgroup></table>"],tr:[2,"<table><tbody>","</tbody></table>"],td:[3,"<table><tbody><tr>","</tr></tbody></table>"],_default:[0,"",""]};ge.optgroup=ge.option,ge.tbody=ge.tfoot=ge.colgroup=ge.caption=ge.thead,ge.th=ge.td;function ye(e,t){var n;return n="undefined"!=typeof e.getElementsByTagName?e.getElementsByTagName(t||"*"):"undefined"!=typeof e.querySelectorAll?e.querySelectorAll(t||"*"):[],void 0===t||t&&N(e,t)?w.merge([e],n):n}function ve(e,t){for(var n=0,r=e.length;n<r;n++)J.set(e[n],"globalEval",!t||J.get(t[n],"globalEval"))}var me=/<|&#?\w+;/;function xe(e,t,n,r,i){for(var o,a,s,u,l,c,f=t.createDocumentFragment(),p=[],d=0,h=e.length;d<h;d++)if((o=e[d])||0===o)if("object"===x(o))w.merge(p,o.nodeType?[o]:o);else if(me.test(o)){a=a||f.appendChild(t.createElement("div")),s=(de.exec(o)||["",""])[1].toLowerCase(),u=ge[s]||ge._default,a.innerHTML=u[1]+w.htmlPrefilter(o)+u[2],c=u[0];while(c--)a=a.lastChild;w.merge(p,a.childNodes),(a=f.firstChild).textContent=""}else p.push(t.createTextNode(o));f.textContent="",d=0;while(o=p[d++])if(r&&w.inArray(o,r)>-1)i&&i.push(o);else if(l=w.contains(o.ownerDocument,o),a=ye(f.appendChild(o),"script"),l&&ve(a),n){c=0;while(o=a[c++])he.test(o.type||"")&&n.push(o)}return f}!function(){var e=r.createDocumentFragment().appendChild(r.createElement("div")),t=r.createElement("input");t.setAttribute("type","radio"),t.setAttribute("checked","checked"),t.setAttribute("name","t"),e.appendChild(t),h.checkClone=e.cloneNode(!0).cloneNode(!0).lastChild.checked,e.innerHTML="<textarea>x</textarea>",h.noCloneChecked=!!e.cloneNode(!0).lastChild.defaultValue}();var be=r.documentElement,we=/^key/,Te=/^(?:mouse|pointer|contextmenu|drag|drop)|click/,Ce=/^([^.]*)(?:\.(.+)|)/;function Ee(){return!0}function ke(){return!1}function Se(){try{return r.activeElement}catch(e){}}function De(e,t,n,r,i,o){var a,s;if("object"==typeof t){"string"!=typeof n&&(r=r||n,n=void 0);for(s in t)De(e,s,n,r,t[s],o);return e}if(null==r&&null==i?(i=n,r=n=void 0):null==i&&("string"==typeof n?(i=r,r=void 0):(i=r,r=n,n=void 0)),!1===i)i=ke;else if(!i)return e;return 1===o&&(a=i,(i=function(e){return w().off(e),a.apply(this,arguments)}).guid=a.guid||(a.guid=w.guid++)),e.each(function(){w.event.add(this,t,i,r,n)})}w.event={global:{},add:function(e,t,n,r,i){var o,a,s,u,l,c,f,p,d,h,g,y=J.get(e);if(y){n.handler&&(n=(o=n).handler,i=o.selector),i&&w.find.matchesSelector(be,i),n.guid||(n.guid=w.guid++),(u=y.events)||(u=y.events={}),(a=y.handle)||(a=y.handle=function(t){return"undefined"!=typeof w&&w.event.triggered!==t.type?w.event.dispatch.apply(e,arguments):void 0}),l=(t=(t||"").match(M)||[""]).length;while(l--)d=g=(s=Ce.exec(t[l])||[])[1],h=(s[2]||"").split(".").sort(),d&&(f=w.event.special[d]||{},d=(i?f.delegateType:f.bindType)||d,f=w.event.special[d]||{},c=w.extend({type:d,origType:g,data:r,handler:n,guid:n.guid,selector:i,needsContext:i&&w.expr.match.needsContext.test(i),namespace:h.join(".")},o),(p=u[d])||((p=u[d]=[]).delegateCount=0,f.setup&&!1!==f.setup.call(e,r,h,a)||e.addEventListener&&e.addEventListener(d,a)),f.add&&(f.add.call(e,c),c.handler.guid||(c.handler.guid=n.guid)),i?p.splice(p.delegateCount++,0,c):p.push(c),w.event.global[d]=!0)}},remove:function(e,t,n,r,i){var o,a,s,u,l,c,f,p,d,h,g,y=J.hasData(e)&&J.get(e);if(y&&(u=y.events)){l=(t=(t||"").match(M)||[""]).length;while(l--)if(s=Ce.exec(t[l])||[],d=g=s[1],h=(s[2]||"").split(".").sort(),d){f=w.event.special[d]||{},p=u[d=(r?f.delegateType:f.bindType)||d]||[],s=s[2]&&new RegExp("(^|\\.)"+h.join("\\.(?:.*\\.|)")+"(\\.|$)"),a=o=p.length;while(o--)c=p[o],!i&&g!==c.origType||n&&n.guid!==c.guid||s&&!s.test(c.namespace)||r&&r!==c.selector&&("**"!==r||!c.selector)||(p.splice(o,1),c.selector&&p.delegateCount--,f.remove&&f.remove.call(e,c));a&&!p.length&&(f.teardown&&!1!==f.teardown.call(e,h,y.handle)||w.removeEvent(e,d,y.handle),delete u[d])}else for(d in u)w.event.remove(e,d+t[l],n,r,!0);w.isEmptyObject(u)&&J.remove(e,"handle events")}},dispatch:function(e){var t=w.event.fix(e),n,r,i,o,a,s,u=new Array(arguments.length),l=(J.get(this,"events")||{})[t.type]||[],c=w.event.special[t.type]||{};for(u[0]=t,n=1;n<arguments.length;n++)u[n]=arguments[n];if(t.delegateTarget=this,!c.preDispatch||!1!==c.preDispatch.call(this,t)){s=w.event.handlers.call(this,t,l),n=0;while((o=s[n++])&&!t.isPropagationStopped()){t.currentTarget=o.elem,r=0;while((a=o.handlers[r++])&&!t.isImmediatePropagationStopped())t.rnamespace&&!t.rnamespace.test(a.namespace)||(t.handleObj=a,t.data=a.data,void 0!==(i=((w.event.special[a.origType]||{}).handle||a.handler).apply(o.elem,u))&&!1===(t.result=i)&&(t.preventDefault(),t.stopPropagation()))}return c.postDispatch&&c.postDispatch.call(this,t),t.result}},handlers:function(e,t){var n,r,i,o,a,s=[],u=t.delegateCount,l=e.target;if(u&&l.nodeType&&!("click"===e.type&&e.button>=1))for(;l!==this;l=l.parentNode||this)if(1===l.nodeType&&("click"!==e.type||!0!==l.disabled)){for(o=[],a={},n=0;n<u;n++)void 0===a[i=(r=t[n]).selector+" "]&&(a[i]=r.needsContext?w(i,this).index(l)>-1:w.find(i,this,null,[l]).length),a[i]&&o.push(r);o.length&&s.push({elem:l,handlers:o})}return l=this,u<t.length&&s.push({elem:l,handlers:t.slice(u)}),s},addProp:function(e,t){Object.defineProperty(w.Event.prototype,e,{enumerable:!0,configurable:!0,get:g(t)?function(){if(this.originalEvent)return t(this.originalEvent)}:function(){if(this.originalEvent)return this.originalEvent[e]},set:function(t){Object.defineProperty(this,e,{enumerable:!0,configurable:!0,writable:!0,value:t})}})},fix:function(e){return e[w.expando]?e:new w.Event(e)},special:{load:{noBubble:!0},focus:{trigger:function(){if(this!==Se()&&this.focus)return this.focus(),!1},delegateType:"focusin"},blur:{trigger:function(){if(this===Se()&&this.blur)return this.blur(),!1},delegateType:"focusout"},click:{trigger:function(){if("checkbox"===this.type&&this.click&&N(this,"input"))return this.click(),!1},_default:function(e){return N(e.target,"a")}},beforeunload:{postDispatch:function(e){void 0!==e.result&&e.originalEvent&&(e.originalEvent.returnValue=e.result)}}}},w.removeEvent=function(e,t,n){e.removeEventListener&&e.removeEventListener(t,n)},w.Event=function(e,t){if(!(this instanceof w.Event))return new w.Event(e,t);e&&e.type?(this.originalEvent=e,this.type=e.type,this.isDefaultPrevented=e.defaultPrevented||void 0===e.defaultPrevented&&!1===e.returnValue?Ee:ke,this.target=e.target&&3===e.target.nodeType?e.target.parentNode:e.target,this.currentTarget=e.currentTarget,this.relatedTarget=e.relatedTarget):this.type=e,t&&w.extend(this,t),this.timeStamp=e&&e.timeStamp||Date.now(),this[w.expando]=!0},w.Event.prototype={constructor:w.Event,isDefaultPrevented:ke,isPropagationStopped:ke,isImmediatePropagationStopped:ke,isSimulated:!1,preventDefault:function(){var e=this.originalEvent;this.isDefaultPrevented=Ee,e&&!this.isSimulated&&e.preventDefault()},stopPropagation:function(){var e=this.originalEvent;this.isPropagationStopped=Ee,e&&!this.isSimulated&&e.stopPropagation()},stopImmediatePropagation:function(){var e=this.originalEvent;this.isImmediatePropagationStopped=Ee,e&&!this.isSimulated&&e.stopImmediatePropagation(),this.stopPropagation()}},w.each({altKey:!0,bubbles:!0,cancelable:!0,changedTouches:!0,ctrlKey:!0,detail:!0,eventPhase:!0,metaKey:!0,pageX:!0,pageY:!0,shiftKey:!0,view:!0,"char":!0,charCode:!0,key:!0,keyCode:!0,button:!0,buttons:!0,clientX:!0,clientY:!0,offsetX:!0,offsetY:!0,pointerId:!0,pointerType:!0,screenX:!0,screenY:!0,targetTouches:!0,toElement:!0,touches:!0,which:function(e){var t=e.button;return null==e.which&&we.test(e.type)?null!=e.charCode?e.charCode:e.keyCode:!e.which&&void 0!==t&&Te.test(e.type)?1&t?1:2&t?3:4&t?2:0:e.which}},w.event.addProp),w.each({mouseenter:"mouseover",mouseleave:"mouseout",pointerenter:"pointerover",pointerleave:"pointerout"},function(e,t){w.event.special[e]={delegateType:t,bindType:t,handle:function(e){var n,r=this,i=e.relatedTarget,o=e.handleObj;return i&&(i===r||w.contains(r,i))||(e.type=o.origType,n=o.handler.apply(this,arguments),e.type=t),n}}}),w.fn.extend({on:function(e,t,n,r){return De(this,e,t,n,r)},one:function(e,t,n,r){return De(this,e,t,n,r,1)},off:function(e,t,n){var r,i;if(e&&e.preventDefault&&e.handleObj)return r=e.handleObj,w(e.delegateTarget).off(r.namespace?r.origType+"."+r.namespace:r.origType,r.selector,r.handler),this;if("object"==typeof e){for(i in e)this.off(i,t,e[i]);return this}return!1!==t&&"function"!=typeof t||(n=t,t=void 0),!1===n&&(n=ke),this.each(function(){w.event.remove(this,e,n,t)})}});var Ne=/<(?!area|br|col|embed|hr|img|input|link|meta|param)(([a-z][^\/\0>\x20\t\r\n\f]*)[^>]*)\/>/gi,Ae=/<script|<style|<link/i,je=/checked\s*(?:[^=]|=\s*.checked.)/i,qe=/^\s*<!(?:\[CDATA\[|--)|(?:\]\]|--)>\s*$/g;function Le(e,t){return N(e,"table")&&N(11!==t.nodeType?t:t.firstChild,"tr")?w(e).children("tbody")[0]||e:e}function He(e){return e.type=(null!==e.getAttribute("type"))+"/"+e.type,e}function Oe(e){return"true/"===(e.type||"").slice(0,5)?e.type=e.type.slice(5):e.removeAttribute("type"),e}function Pe(e,t){var n,r,i,o,a,s,u,l;if(1===t.nodeType){if(J.hasData(e)&&(o=J.access(e),a=J.set(t,o),l=o.events)){delete a.handle,a.events={};for(i in l)for(n=0,r=l[i].length;n<r;n++)w.event.add(t,i,l[i][n])}K.hasData(e)&&(s=K.access(e),u=w.extend({},s),K.set(t,u))}}function Me(e,t){var n=t.nodeName.toLowerCase();"input"===n&&pe.test(e.type)?t.checked=e.checked:"input"!==n&&"textarea"!==n||(t.defaultValue=e.defaultValue)}function Re(e,t,n,r){t=a.apply([],t);var i,o,s,u,l,c,f=0,p=e.length,d=p-1,y=t[0],v=g(y);if(v||p>1&&"string"==typeof y&&!h.checkClone&&je.test(y))return e.each(function(i){var o=e.eq(i);v&&(t[0]=y.call(this,i,o.html())),Re(o,t,n,r)});if(p&&(i=xe(t,e[0].ownerDocument,!1,e,r),o=i.firstChild,1===i.childNodes.length&&(i=o),o||r)){for(u=(s=w.map(ye(i,"script"),He)).length;f<p;f++)l=i,f!==d&&(l=w.clone(l,!0,!0),u&&w.merge(s,ye(l,"script"))),n.call(e[f],l,f);if(u)for(c=s[s.length-1].ownerDocument,w.map(s,Oe),f=0;f<u;f++)l=s[f],he.test(l.type||"")&&!J.access(l,"globalEval")&&w.contains(c,l)&&(l.src&&"module"!==(l.type||"").toLowerCase()?w._evalUrl&&w._evalUrl(l.src):m(l.textContent.replace(qe,""),c,l))}return e}function Ie(e,t,n){for(var r,i=t?w.filter(t,e):e,o=0;null!=(r=i[o]);o++)n||1!==r.nodeType||w.cleanData(ye(r)),r.parentNode&&(n&&w.contains(r.ownerDocument,r)&&ve(ye(r,"script")),r.parentNode.removeChild(r));return e}w.extend({htmlPrefilter:function(e){return e.replace(Ne,"<$1></$2>")},clone:function(e,t,n){var r,i,o,a,s=e.cloneNode(!0),u=w.contains(e.ownerDocument,e);if(!(h.noCloneChecked||1!==e.nodeType&&11!==e.nodeType||w.isXMLDoc(e)))for(a=ye(s),r=0,i=(o=ye(e)).length;r<i;r++)Me(o[r],a[r]);if(t)if(n)for(o=o||ye(e),a=a||ye(s),r=0,i=o.length;r<i;r++)Pe(o[r],a[r]);else Pe(e,s);return(a=ye(s,"script")).length>0&&ve(a,!u&&ye(e,"script")),s},cleanData:function(e){for(var t,n,r,i=w.event.special,o=0;void 0!==(n=e[o]);o++)if(Y(n)){if(t=n[J.expando]){if(t.events)for(r in t.events)i[r]?w.event.remove(n,r):w.removeEvent(n,r,t.handle);n[J.expando]=void 0}n[K.expando]&&(n[K.expando]=void 0)}}}),w.fn.extend({detach:function(e){return Ie(this,e,!0)},remove:function(e){return Ie(this,e)},text:function(e){return z(this,function(e){return void 0===e?w.text(this):this.empty().each(function(){1!==this.nodeType&&11!==this.nodeType&&9!==this.nodeType||(this.textContent=e)})},null,e,arguments.length)},append:function(){return Re(this,arguments,function(e){1!==this.nodeType&&11!==this.nodeType&&9!==this.nodeType||Le(this,e).appendChild(e)})},prepend:function(){return Re(this,arguments,function(e){if(1===this.nodeType||11===this.nodeType||9===this.nodeType){var t=Le(this,e);t.insertBefore(e,t.firstChild)}})},before:function(){return Re(this,arguments,function(e){this.parentNode&&this.parentNode.insertBefore(e,this)})},after:function(){return Re(this,arguments,function(e){this.parentNode&&this.parentNode.insertBefore(e,this.nextSibling)})},empty:function(){for(var e,t=0;null!=(e=this[t]);t++)1===e.nodeType&&(w.cleanData(ye(e,!1)),e.textContent="");return this},clone:function(e,t){return e=null!=e&&e,t=null==t?e:t,this.map(function(){return w.clone(this,e,t)})},html:function(e){return z(this,function(e){var t=this[0]||{},n=0,r=this.length;if(void 0===e&&1===t.nodeType)return t.innerHTML;if("string"==typeof e&&!Ae.test(e)&&!ge[(de.exec(e)||["",""])[1].toLowerCase()]){e=w.htmlPrefilter(e);try{for(;n<r;n++)1===(t=this[n]||{}).nodeType&&(w.cleanData(ye(t,!1)),t.innerHTML=e);t=0}catch(e){}}t&&this.empty().append(e)},null,e,arguments.length)},replaceWith:function(){var e=[];return Re(this,arguments,function(t){var n=this.parentNode;w.inArray(this,e)<0&&(w.cleanData(ye(this)),n&&n.replaceChild(t,this))},e)}}),w.each({appendTo:"append",prependTo:"prepend",insertBefore:"before",insertAfter:"after",replaceAll:"replaceWith"},function(e,t){w.fn[e]=function(e){for(var n,r=[],i=w(e),o=i.length-1,a=0;a<=o;a++)n=a===o?this:this.clone(!0),w(i[a])[t](n),s.apply(r,n.get());return this.pushStack(r)}});var We=new RegExp("^("+re+")(?!px)[a-z%]+$","i"),$e=function(t){var n=t.ownerDocument.defaultView;return n&&n.opener||(n=e),n.getComputedStyle(t)},Be=new RegExp(oe.join("|"),"i");!function(){function t(){if(c){l.style.cssText="position:absolute;left:-11111px;width:60px;margin-top:1px;padding:0;border:0",c.style.cssText="position:relative;display:block;box-sizing:border-box;overflow:scroll;margin:auto;border:0px;padding:1px;width:60%;top:1%",be.appendChild(l).appendChild(c);var t=e.getComputedStyle(c);i="1%"!==t.top,u=12===n(t.marginLeft),c.style.right="60%",s=36===n(t.right),o=36===n(t.width),c.style.position="absolute",a=36===c.offsetWidth||"absolute",be.removeChild(l),c=null}}function n(e){return Math.round(parseFloat(e))}var i,o,a,s,u,l=r.createElement("div"),c=r.createElement("div");c.style&&(c.style.backgroundClip="content-box",c.cloneNode(!0).style.backgroundClip="",h.clearCloneStyle="content-box"===c.style.backgroundClip,w.extend(h,{boxSizingReliable:function(){return t(),o},pixelBoxStyles:function(){return t(),s},pixelPosition:function(){return t(),i},reliableMarginLeft:function(){return t(),u},scrollboxSize:function(){return t(),a}}))}();function Fe(e,t,n){var r,i,o,a,s=e.style;return(n=n||$e(e))&&(""!==(a=n.getPropertyValue(t)||n[t])||w.contains(e.ownerDocument,e)||(a=w.style(e,t)),!h.pixelBoxStyles()&&We.test(a)&&Be.test(t)&&(r=s.width,i=s.minWidth,o=s.maxWidth,s.minWidth=s.maxWidth=s.width=a,a=n.width,s.width=r,s.minWidth=i,s.maxWidth=o)),void 0!==a?a+"":a}function _e(e,t){return{get:function(){if(!e())return(this.get=t).apply(this,arguments);delete this.get}}}var ze=/^(none|table(?!-c[ea]).+)/,Xe=/^--/,Ue={position:"absolute",visibility:"hidden",display:"block"},Ve={letterSpacing:"0",fontWeight:"400"},Ge=["Webkit","Moz","ms"],Ye=r.createElement("div").style;function Qe(e){if(e in Ye)return e;var t=e[0].toUpperCase()+e.slice(1),n=Ge.length;while(n--)if((e=Ge[n]+t)in Ye)return e}function Je(e){var t=w.cssProps[e];return t||(t=w.cssProps[e]=Qe(e)||e),t}function Ke(e,t,n){var r=ie.exec(t);return r?Math.max(0,r[2]-(n||0))+(r[3]||"px"):t}function Ze(e,t,n,r,i,o){var a="width"===t?1:0,s=0,u=0;if(n===(r?"border":"content"))return 0;for(;a<4;a+=2)"margin"===n&&(u+=w.css(e,n+oe[a],!0,i)),r?("content"===n&&(u-=w.css(e,"padding"+oe[a],!0,i)),"margin"!==n&&(u-=w.css(e,"border"+oe[a]+"Width",!0,i))):(u+=w.css(e,"padding"+oe[a],!0,i),"padding"!==n?u+=w.css(e,"border"+oe[a]+"Width",!0,i):s+=w.css(e,"border"+oe[a]+"Width",!0,i));return!r&&o>=0&&(u+=Math.max(0,Math.ceil(e["offset"+t[0].toUpperCase()+t.slice(1)]-o-u-s-.5))),u}function et(e,t,n){var r=$e(e),i=Fe(e,t,r),o="border-box"===w.css(e,"boxSizing",!1,r),a=o;if(We.test(i)){if(!n)return i;i="auto"}return a=a&&(h.boxSizingReliable()||i===e.style[t]),("auto"===i||!parseFloat(i)&&"inline"===w.css(e,"display",!1,r))&&(i=e["offset"+t[0].toUpperCase()+t.slice(1)],a=!0),(i=parseFloat(i)||0)+Ze(e,t,n||(o?"border":"content"),a,r,i)+"px"}w.extend({cssHooks:{opacity:{get:function(e,t){if(t){var n=Fe(e,"opacity");return""===n?"1":n}}}},cssNumber:{animationIterationCount:!0,columnCount:!0,fillOpacity:!0,flexGrow:!0,flexShrink:!0,fontWeight:!0,lineHeight:!0,opacity:!0,order:!0,orphans:!0,widows:!0,zIndex:!0,zoom:!0},cssProps:{},style:function(e,t,n,r){if(e&&3!==e.nodeType&&8!==e.nodeType&&e.style){var i,o,a,s=G(t),u=Xe.test(t),l=e.style;if(u||(t=Je(s)),a=w.cssHooks[t]||w.cssHooks[s],void 0===n)return a&&"get"in a&&void 0!==(i=a.get(e,!1,r))?i:l[t];"string"==(o=typeof n)&&(i=ie.exec(n))&&i[1]&&(n=ue(e,t,i),o="number"),null!=n&&n===n&&("number"===o&&(n+=i&&i[3]||(w.cssNumber[s]?"":"px")),h.clearCloneStyle||""!==n||0!==t.indexOf("background")||(l[t]="inherit"),a&&"set"in a&&void 0===(n=a.set(e,n,r))||(u?l.setProperty(t,n):l[t]=n))}},css:function(e,t,n,r){var i,o,a,s=G(t);return Xe.test(t)||(t=Je(s)),(a=w.cssHooks[t]||w.cssHooks[s])&&"get"in a&&(i=a.get(e,!0,n)),void 0===i&&(i=Fe(e,t,r)),"normal"===i&&t in Ve&&(i=Ve[t]),""===n||n?(o=parseFloat(i),!0===n||isFinite(o)?o||0:i):i}}),w.each(["height","width"],function(e,t){w.cssHooks[t]={get:function(e,n,r){if(n)return!ze.test(w.css(e,"display"))||e.getClientRects().length&&e.getBoundingClientRect().width?et(e,t,r):se(e,Ue,function(){return et(e,t,r)})},set:function(e,n,r){var i,o=$e(e),a="border-box"===w.css(e,"boxSizing",!1,o),s=r&&Ze(e,t,r,a,o);return a&&h.scrollboxSize()===o.position&&(s-=Math.ceil(e["offset"+t[0].toUpperCase()+t.slice(1)]-parseFloat(o[t])-Ze(e,t,"border",!1,o)-.5)),s&&(i=ie.exec(n))&&"px"!==(i[3]||"px")&&(e.style[t]=n,n=w.css(e,t)),Ke(e,n,s)}}}),w.cssHooks.marginLeft=_e(h.reliableMarginLeft,function(e,t){if(t)return(parseFloat(Fe(e,"marginLeft"))||e.getBoundingClientRect().left-se(e,{marginLeft:0},function(){return e.getBoundingClientRect().left}))+"px"}),w.each({margin:"",padding:"",border:"Width"},function(e,t){w.cssHooks[e+t]={expand:function(n){for(var r=0,i={},o="string"==typeof n?n.split(" "):[n];r<4;r++)i[e+oe[r]+t]=o[r]||o[r-2]||o[0];return i}},"margin"!==e&&(w.cssHooks[e+t].set=Ke)}),w.fn.extend({css:function(e,t){return z(this,function(e,t,n){var r,i,o={},a=0;if(Array.isArray(t)){for(r=$e(e),i=t.length;a<i;a++)o[t[a]]=w.css(e,t[a],!1,r);return o}return void 0!==n?w.style(e,t,n):w.css(e,t)},e,t,arguments.length>1)}});function tt(e,t,n,r,i){return new tt.prototype.init(e,t,n,r,i)}w.Tween=tt,tt.prototype={constructor:tt,init:function(e,t,n,r,i,o){this.elem=e,this.prop=n,this.easing=i||w.easing._default,this.options=t,this.start=this.now=this.cur(),this.end=r,this.unit=o||(w.cssNumber[n]?"":"px")},cur:function(){var e=tt.propHooks[this.prop];return e&&e.get?e.get(this):tt.propHooks._default.get(this)},run:function(e){var t,n=tt.propHooks[this.prop];return this.options.duration?this.pos=t=w.easing[this.easing](e,this.options.duration*e,0,1,this.options.duration):this.pos=t=e,this.now=(this.end-this.start)*t+this.start,this.options.step&&this.options.step.call(this.elem,this.now,this),n&&n.set?n.set(this):tt.propHooks._default.set(this),this}},tt.prototype.init.prototype=tt.prototype,tt.propHooks={_default:{get:function(e){var t;return 1!==e.elem.nodeType||null!=e.elem[e.prop]&&null==e.elem.style[e.prop]?e.elem[e.prop]:(t=w.css(e.elem,e.prop,""))&&"auto"!==t?t:0},set:function(e){w.fx.step[e.prop]?w.fx.step[e.prop](e):1!==e.elem.nodeType||null==e.elem.style[w.cssProps[e.prop]]&&!w.cssHooks[e.prop]?e.elem[e.prop]=e.now:w.style(e.elem,e.prop,e.now+e.unit)}}},tt.propHooks.scrollTop=tt.propHooks.scrollLeft={set:function(e){e.elem.nodeType&&e.elem.parentNode&&(e.elem[e.prop]=e.now)}},w.easing={linear:function(e){return e},swing:function(e){return.5-Math.cos(e*Math.PI)/2},_default:"swing"},w.fx=tt.prototype.init,w.fx.step={};var nt,rt,it=/^(?:toggle|show|hide)$/,ot=/queueHooks$/;function at(){rt&&(!1===r.hidden&&e.requestAnimationFrame?e.requestAnimationFrame(at):e.setTimeout(at,w.fx.interval),w.fx.tick())}function st(){return e.setTimeout(function(){nt=void 0}),nt=Date.now()}function ut(e,t){var n,r=0,i={height:e};for(t=t?1:0;r<4;r+=2-t)i["margin"+(n=oe[r])]=i["padding"+n]=e;return t&&(i.opacity=i.width=e),i}function lt(e,t,n){for(var r,i=(pt.tweeners[t]||[]).concat(pt.tweeners["*"]),o=0,a=i.length;o<a;o++)if(r=i[o].call(n,t,e))return r}function ct(e,t,n){var r,i,o,a,s,u,l,c,f="width"in t||"height"in t,p=this,d={},h=e.style,g=e.nodeType&&ae(e),y=J.get(e,"fxshow");n.queue||(null==(a=w._queueHooks(e,"fx")).unqueued&&(a.unqueued=0,s=a.empty.fire,a.empty.fire=function(){a.unqueued||s()}),a.unqueued++,p.always(function(){p.always(function(){a.unqueued--,w.queue(e,"fx").length||a.empty.fire()})}));for(r in t)if(i=t[r],it.test(i)){if(delete t[r],o=o||"toggle"===i,i===(g?"hide":"show")){if("show"!==i||!y||void 0===y[r])continue;g=!0}d[r]=y&&y[r]||w.style(e,r)}if((u=!w.isEmptyObject(t))||!w.isEmptyObject(d)){f&&1===e.nodeType&&(n.overflow=[h.overflow,h.overflowX,h.overflowY],null==(l=y&&y.display)&&(l=J.get(e,"display")),"none"===(c=w.css(e,"display"))&&(l?c=l:(fe([e],!0),l=e.style.display||l,c=w.css(e,"display"),fe([e]))),("inline"===c||"inline-block"===c&&null!=l)&&"none"===w.css(e,"float")&&(u||(p.done(function(){h.display=l}),null==l&&(c=h.display,l="none"===c?"":c)),h.display="inline-block")),n.overflow&&(h.overflow="hidden",p.always(function(){h.overflow=n.overflow[0],h.overflowX=n.overflow[1],h.overflowY=n.overflow[2]})),u=!1;for(r in d)u||(y?"hidden"in y&&(g=y.hidden):y=J.access(e,"fxshow",{display:l}),o&&(y.hidden=!g),g&&fe([e],!0),p.done(function(){g||fe([e]),J.remove(e,"fxshow");for(r in d)w.style(e,r,d[r])})),u=lt(g?y[r]:0,r,p),r in y||(y[r]=u.start,g&&(u.end=u.start,u.start=0))}}function ft(e,t){var n,r,i,o,a;for(n in e)if(r=G(n),i=t[r],o=e[n],Array.isArray(o)&&(i=o[1],o=e[n]=o[0]),n!==r&&(e[r]=o,delete e[n]),(a=w.cssHooks[r])&&"expand"in a){o=a.expand(o),delete e[r];for(n in o)n in e||(e[n]=o[n],t[n]=i)}else t[r]=i}function pt(e,t,n){var r,i,o=0,a=pt.prefilters.length,s=w.Deferred().always(function(){delete u.elem}),u=function(){if(i)return!1;for(var t=nt||st(),n=Math.max(0,l.startTime+l.duration-t),r=1-(n/l.duration||0),o=0,a=l.tweens.length;o<a;o++)l.tweens[o].run(r);return s.notifyWith(e,[l,r,n]),r<1&&a?n:(a||s.notifyWith(e,[l,1,0]),s.resolveWith(e,[l]),!1)},l=s.promise({elem:e,props:w.extend({},t),opts:w.extend(!0,{specialEasing:{},easing:w.easing._default},n),originalProperties:t,originalOptions:n,startTime:nt||st(),duration:n.duration,tweens:[],createTween:function(t,n){var r=w.Tween(e,l.opts,t,n,l.opts.specialEasing[t]||l.opts.easing);return l.tweens.push(r),r},stop:function(t){var n=0,r=t?l.tweens.length:0;if(i)return this;for(i=!0;n<r;n++)l.tweens[n].run(1);return t?(s.notifyWith(e,[l,1,0]),s.resolveWith(e,[l,t])):s.rejectWith(e,[l,t]),this}}),c=l.props;for(ft(c,l.opts.specialEasing);o<a;o++)if(r=pt.prefilters[o].call(l,e,c,l.opts))return g(r.stop)&&(w._queueHooks(l.elem,l.opts.queue).stop=r.stop.bind(r)),r;return w.map(c,lt,l),g(l.opts.start)&&l.opts.start.call(e,l),l.progress(l.opts.progress).done(l.opts.done,l.opts.complete).fail(l.opts.fail).always(l.opts.always),w.fx.timer(w.extend(u,{elem:e,anim:l,queue:l.opts.queue})),l}w.Animation=w.extend(pt,{tweeners:{"*":[function(e,t){var n=this.createTween(e,t);return ue(n.elem,e,ie.exec(t),n),n}]},tweener:function(e,t){g(e)?(t=e,e=["*"]):e=e.match(M);for(var n,r=0,i=e.length;r<i;r++)n=e[r],pt.tweeners[n]=pt.tweeners[n]||[],pt.tweeners[n].unshift(t)},prefilters:[ct],prefilter:function(e,t){t?pt.prefilters.unshift(e):pt.prefilters.push(e)}}),w.speed=function(e,t,n){var r=e&&"object"==typeof e?w.extend({},e):{complete:n||!n&&t||g(e)&&e,duration:e,easing:n&&t||t&&!g(t)&&t};return w.fx.off?r.duration=0:"number"!=typeof r.duration&&(r.duration in w.fx.speeds?r.duration=w.fx.speeds[r.duration]:r.duration=w.fx.speeds._default),null!=r.queue&&!0!==r.queue||(r.queue="fx"),r.old=r.complete,r.complete=function(){g(r.old)&&r.old.call(this),r.queue&&w.dequeue(this,r.queue)},r},w.fn.extend({fadeTo:function(e,t,n,r){return this.filter(ae).css("opacity",0).show().end().animate({opacity:t},e,n,r)},animate:function(e,t,n,r){var i=w.isEmptyObject(e),o=w.speed(t,n,r),a=function(){var t=pt(this,w.extend({},e),o);(i||J.get(this,"finish"))&&t.stop(!0)};return a.finish=a,i||!1===o.queue?this.each(a):this.queue(o.queue,a)},stop:function(e,t,n){var r=function(e){var t=e.stop;delete e.stop,t(n)};return"string"!=typeof e&&(n=t,t=e,e=void 0),t&&!1!==e&&this.queue(e||"fx",[]),this.each(function(){var t=!0,i=null!=e&&e+"queueHooks",o=w.timers,a=J.get(this);if(i)a[i]&&a[i].stop&&r(a[i]);else for(i in a)a[i]&&a[i].stop&&ot.test(i)&&r(a[i]);for(i=o.length;i--;)o[i].elem!==this||null!=e&&o[i].queue!==e||(o[i].anim.stop(n),t=!1,o.splice(i,1));!t&&n||w.dequeue(this,e)})},finish:function(e){return!1!==e&&(e=e||"fx"),this.each(function(){var t,n=J.get(this),r=n[e+"queue"],i=n[e+"queueHooks"],o=w.timers,a=r?r.length:0;for(n.finish=!0,w.queue(this,e,[]),i&&i.stop&&i.stop.call(this,!0),t=o.length;t--;)o[t].elem===this&&o[t].queue===e&&(o[t].anim.stop(!0),o.splice(t,1));for(t=0;t<a;t++)r[t]&&r[t].finish&&r[t].finish.call(this);delete n.finish})}}),w.each(["toggle","show","hide"],function(e,t){var n=w.fn[t];w.fn[t]=function(e,r,i){return null==e||"boolean"==typeof e?n.apply(this,arguments):this.animate(ut(t,!0),e,r,i)}}),w.each({slideDown:ut("show"),slideUp:ut("hide"),slideToggle:ut("toggle"),fadeIn:{opacity:"show"},fadeOut:{opacity:"hide"},fadeToggle:{opacity:"toggle"}},function(e,t){w.fn[e]=function(e,n,r){return this.animate(t,e,n,r)}}),w.timers=[],w.fx.tick=function(){var e,t=0,n=w.timers;for(nt=Date.now();t<n.length;t++)(e=n[t])()||n[t]!==e||n.splice(t--,1);n.length||w.fx.stop(),nt=void 0},w.fx.timer=function(e){w.timers.push(e),w.fx.start()},w.fx.interval=13,w.fx.start=function(){rt||(rt=!0,at())},w.fx.stop=function(){rt=null},w.fx.speeds={slow:600,fast:200,_default:400},w.fn.delay=function(t,n){return t=w.fx?w.fx.speeds[t]||t:t,n=n||"fx",this.queue(n,function(n,r){var i=e.setTimeout(n,t);r.stop=function(){e.clearTimeout(i)}})},function(){var e=r.createElement("input"),t=r.createElement("select").appendChild(r.createElement("option"));e.type="checkbox",h.checkOn=""!==e.value,h.optSelected=t.selected,(e=r.createElement("input")).value="t",e.type="radio",h.radioValue="t"===e.value}();var dt,ht=w.expr.attrHandle;w.fn.extend({attr:function(e,t){return z(this,w.attr,e,t,arguments.length>1)},removeAttr:function(e){return this.each(function(){w.removeAttr(this,e)})}}),w.extend({attr:function(e,t,n){var r,i,o=e.nodeType;if(3!==o&&8!==o&&2!==o)return"undefined"==typeof e.getAttribute?w.prop(e,t,n):(1===o&&w.isXMLDoc(e)||(i=w.attrHooks[t.toLowerCase()]||(w.expr.match.bool.test(t)?dt:void 0)),void 0!==n?null===n?void w.removeAttr(e,t):i&&"set"in i&&void 0!==(r=i.set(e,n,t))?r:(e.setAttribute(t,n+""),n):i&&"get"in i&&null!==(r=i.get(e,t))?r:null==(r=w.find.attr(e,t))?void 0:r)},attrHooks:{type:{set:function(e,t){if(!h.radioValue&&"radio"===t&&N(e,"input")){var n=e.value;return e.setAttribute("type",t),n&&(e.value=n),t}}}},removeAttr:function(e,t){var n,r=0,i=t&&t.match(M);if(i&&1===e.nodeType)while(n=i[r++])e.removeAttribute(n)}}),dt={set:function(e,t,n){return!1===t?w.removeAttr(e,n):e.setAttribute(n,n),n}},w.each(w.expr.match.bool.source.match(/\w+/g),function(e,t){var n=ht[t]||w.find.attr;ht[t]=function(e,t,r){var i,o,a=t.toLowerCase();return r||(o=ht[a],ht[a]=i,i=null!=n(e,t,r)?a:null,ht[a]=o),i}});var gt=/^(?:input|select|textarea|button)$/i,yt=/^(?:a|area)$/i;w.fn.extend({prop:function(e,t){return z(this,w.prop,e,t,arguments.length>1)},removeProp:function(e){return this.each(function(){delete this[w.propFix[e]||e]})}}),w.extend({prop:function(e,t,n){var r,i,o=e.nodeType;if(3!==o&&8!==o&&2!==o)return 1===o&&w.isXMLDoc(e)||(t=w.propFix[t]||t,i=w.propHooks[t]),void 0!==n?i&&"set"in i&&void 0!==(r=i.set(e,n,t))?r:e[t]=n:i&&"get"in i&&null!==(r=i.get(e,t))?r:e[t]},propHooks:{tabIndex:{get:function(e){var t=w.find.attr(e,"tabindex");return t?parseInt(t,10):gt.test(e.nodeName)||yt.test(e.nodeName)&&e.href?0:-1}}},propFix:{"for":"htmlFor","class":"className"}}),h.optSelected||(w.propHooks.selected={get:function(e){var t=e.parentNode;return t&&t.parentNode&&t.parentNode.selectedIndex,null},set:function(e){var t=e.parentNode;t&&(t.selectedIndex,t.parentNode&&t.parentNode.selectedIndex)}}),w.each(["tabIndex","readOnly","maxLength","cellSpacing","cellPadding","rowSpan","colSpan","useMap","frameBorder","contentEditable"],function(){w.propFix[this.toLowerCase()]=this});function vt(e){return(e.match(M)||[]).join(" ")}function mt(e){return e.getAttribute&&e.getAttribute("class")||""}function xt(e){return Array.isArray(e)?e:"string"==typeof e?e.match(M)||[]:[]}w.fn.extend({addClass:function(e){var t,n,r,i,o,a,s,u=0;if(g(e))return this.each(function(t){w(this).addClass(e.call(this,t,mt(this)))});if((t=xt(e)).length)while(n=this[u++])if(i=mt(n),r=1===n.nodeType&&" "+vt(i)+" "){a=0;while(o=t[a++])r.indexOf(" "+o+" ")<0&&(r+=o+" ");i!==(s=vt(r))&&n.setAttribute("class",s)}return this},removeClass:function(e){var t,n,r,i,o,a,s,u=0;if(g(e))return this.each(function(t){w(this).removeClass(e.call(this,t,mt(this)))});if(!arguments.length)return this.attr("class","");if((t=xt(e)).length)while(n=this[u++])if(i=mt(n),r=1===n.nodeType&&" "+vt(i)+" "){a=0;while(o=t[a++])while(r.indexOf(" "+o+" ")>-1)r=r.replace(" "+o+" "," ");i!==(s=vt(r))&&n.setAttribute("class",s)}return this},toggleClass:function(e,t){var n=typeof e,r="string"===n||Array.isArray(e);return"boolean"==typeof t&&r?t?this.addClass(e):this.removeClass(e):g(e)?this.each(function(n){w(this).toggleClass(e.call(this,n,mt(this),t),t)}):this.each(function(){var t,i,o,a;if(r){i=0,o=w(this),a=xt(e);while(t=a[i++])o.hasClass(t)?o.removeClass(t):o.addClass(t)}else void 0!==e&&"boolean"!==n||((t=mt(this))&&J.set(this,"__className__",t),this.setAttribute&&this.setAttribute("class",t||!1===e?"":J.get(this,"__className__")||""))})},hasClass:function(e){var t,n,r=0;t=" "+e+" ";while(n=this[r++])if(1===n.nodeType&&(" "+vt(mt(n))+" ").indexOf(t)>-1)return!0;return!1}});var bt=/\r/g;w.fn.extend({val:function(e){var t,n,r,i=this[0];{if(arguments.length)return r=g(e),this.each(function(n){var i;1===this.nodeType&&(null==(i=r?e.call(this,n,w(this).val()):e)?i="":"number"==typeof i?i+="":Array.isArray(i)&&(i=w.map(i,function(e){return null==e?"":e+""})),(t=w.valHooks[this.type]||w.valHooks[this.nodeName.toLowerCase()])&&"set"in t&&void 0!==t.set(this,i,"value")||(this.value=i))});if(i)return(t=w.valHooks[i.type]||w.valHooks[i.nodeName.toLowerCase()])&&"get"in t&&void 0!==(n=t.get(i,"value"))?n:"string"==typeof(n=i.value)?n.replace(bt,""):null==n?"":n}}}),w.extend({valHooks:{option:{get:function(e){var t=w.find.attr(e,"value");return null!=t?t:vt(w.text(e))}},select:{get:function(e){var t,n,r,i=e.options,o=e.selectedIndex,a="select-one"===e.type,s=a?null:[],u=a?o+1:i.length;for(r=o<0?u:a?o:0;r<u;r++)if(((n=i[r]).selected||r===o)&&!n.disabled&&(!n.parentNode.disabled||!N(n.parentNode,"optgroup"))){if(t=w(n).val(),a)return t;s.push(t)}return s},set:function(e,t){var n,r,i=e.options,o=w.makeArray(t),a=i.length;while(a--)((r=i[a]).selected=w.inArray(w.valHooks.option.get(r),o)>-1)&&(n=!0);return n||(e.selectedIndex=-1),o}}}}),w.each(["radio","checkbox"],function(){w.valHooks[this]={set:function(e,t){if(Array.isArray(t))return e.checked=w.inArray(w(e).val(),t)>-1}},h.checkOn||(w.valHooks[this].get=function(e){return null===e.getAttribute("value")?"on":e.value})}),h.focusin="onfocusin"in e;var wt=/^(?:focusinfocus|focusoutblur)$/,Tt=function(e){e.stopPropagation()};w.extend(w.event,{trigger:function(t,n,i,o){var a,s,u,l,c,p,d,h,v=[i||r],m=f.call(t,"type")?t.type:t,x=f.call(t,"namespace")?t.namespace.split("."):[];if(s=h=u=i=i||r,3!==i.nodeType&&8!==i.nodeType&&!wt.test(m+w.event.triggered)&&(m.indexOf(".")>-1&&(m=(x=m.split(".")).shift(),x.sort()),c=m.indexOf(":")<0&&"on"+m,t=t[w.expando]?t:new w.Event(m,"object"==typeof t&&t),t.isTrigger=o?2:3,t.namespace=x.join("."),t.rnamespace=t.namespace?new RegExp("(^|\\.)"+x.join("\\.(?:.*\\.|)")+"(\\.|$)"):null,t.result=void 0,t.target||(t.target=i),n=null==n?[t]:w.makeArray(n,[t]),d=w.event.special[m]||{},o||!d.trigger||!1!==d.trigger.apply(i,n))){if(!o&&!d.noBubble&&!y(i)){for(l=d.delegateType||m,wt.test(l+m)||(s=s.parentNode);s;s=s.parentNode)v.push(s),u=s;u===(i.ownerDocument||r)&&v.push(u.defaultView||u.parentWindow||e)}a=0;while((s=v[a++])&&!t.isPropagationStopped())h=s,t.type=a>1?l:d.bindType||m,(p=(J.get(s,"events")||{})[t.type]&&J.get(s,"handle"))&&p.apply(s,n),(p=c&&s[c])&&p.apply&&Y(s)&&(t.result=p.apply(s,n),!1===t.result&&t.preventDefault());return t.type=m,o||t.isDefaultPrevented()||d._default&&!1!==d._default.apply(v.pop(),n)||!Y(i)||c&&g(i[m])&&!y(i)&&((u=i[c])&&(i[c]=null),w.event.triggered=m,t.isPropagationStopped()&&h.addEventListener(m,Tt),i[m](),t.isPropagationStopped()&&h.removeEventListener(m,Tt),w.event.triggered=void 0,u&&(i[c]=u)),t.result}},simulate:function(e,t,n){var r=w.extend(new w.Event,n,{type:e,isSimulated:!0});w.event.trigger(r,null,t)}}),w.fn.extend({trigger:function(e,t){return this.each(function(){w.event.trigger(e,t,this)})},triggerHandler:function(e,t){var n=this[0];if(n)return w.event.trigger(e,t,n,!0)}}),h.focusin||w.each({focus:"focusin",blur:"focusout"},function(e,t){var n=function(e){w.event.simulate(t,e.target,w.event.fix(e))};w.event.special[t]={setup:function(){var r=this.ownerDocument||this,i=J.access(r,t);i||r.addEventListener(e,n,!0),J.access(r,t,(i||0)+1)},teardown:function(){var r=this.ownerDocument||this,i=J.access(r,t)-1;i?J.access(r,t,i):(r.removeEventListener(e,n,!0),J.remove(r,t))}}});var Ct=e.location,Et=Date.now(),kt=/\?/;w.parseXML=function(t){var n;if(!t||"string"!=typeof t)return null;try{n=(new e.DOMParser).parseFromString(t,"text/xml")}catch(e){n=void 0}return n&&!n.getElementsByTagName("parsererror").length||w.error("Invalid XML: "+t),n};var St=/\[\]$/,Dt=/\r?\n/g,Nt=/^(?:submit|button|image|reset|file)$/i,At=/^(?:input|select|textarea|keygen)/i;function jt(e,t,n,r){var i;if(Array.isArray(t))w.each(t,function(t,i){n||St.test(e)?r(e,i):jt(e+"["+("object"==typeof i&&null!=i?t:"")+"]",i,n,r)});else if(n||"object"!==x(t))r(e,t);else for(i in t)jt(e+"["+i+"]",t[i],n,r)}w.param=function(e,t){var n,r=[],i=function(e,t){var n=g(t)?t():t;r[r.length]=encodeURIComponent(e)+"="+encodeURIComponent(null==n?"":n)};if(Array.isArray(e)||e.jquery&&!w.isPlainObject(e))w.each(e,function(){i(this.name,this.value)});else for(n in e)jt(n,e[n],t,i);return r.join("&")},w.fn.extend({serialize:function(){return w.param(this.serializeArray())},serializeArray:function(){return this.map(function(){var e=w.prop(this,"elements");return e?w.makeArray(e):this}).filter(function(){var e=this.type;return this.name&&!w(this).is(":disabled")&&At.test(this.nodeName)&&!Nt.test(e)&&(this.checked||!pe.test(e))}).map(function(e,t){var n=w(this).val();return null==n?null:Array.isArray(n)?w.map(n,function(e){return{name:t.name,value:e.replace(Dt,"\r\n")}}):{name:t.name,value:n.replace(Dt,"\r\n")}}).get()}});var qt=/%20/g,Lt=/#.*$/,Ht=/([?&])_=[^&]*/,Ot=/^(.*?):[ \t]*([^\r\n]*)$/gm,Pt=/^(?:about|app|app-storage|.+-extension|file|res|widget):$/,Mt=/^(?:GET|HEAD)$/,Rt=/^\/\//,It={},Wt={},$t="*/".concat("*"),Bt=r.createElement("a");Bt.href=Ct.href;function Ft(e){return function(t,n){"string"!=typeof t&&(n=t,t="*");var r,i=0,o=t.toLowerCase().match(M)||[];if(g(n))while(r=o[i++])"+"===r[0]?(r=r.slice(1)||"*",(e[r]=e[r]||[]).unshift(n)):(e[r]=e[r]||[]).push(n)}}function _t(e,t,n,r){var i={},o=e===Wt;function a(s){var u;return i[s]=!0,w.each(e[s]||[],function(e,s){var l=s(t,n,r);return"string"!=typeof l||o||i[l]?o?!(u=l):void 0:(t.dataTypes.unshift(l),a(l),!1)}),u}return a(t.dataTypes[0])||!i["*"]&&a("*")}function zt(e,t){var n,r,i=w.ajaxSettings.flatOptions||{};for(n in t)void 0!==t[n]&&((i[n]?e:r||(r={}))[n]=t[n]);return r&&w.extend(!0,e,r),e}function Xt(e,t,n){var r,i,o,a,s=e.contents,u=e.dataTypes;while("*"===u[0])u.shift(),void 0===r&&(r=e.mimeType||t.getResponseHeader("Content-Type"));if(r)for(i in s)if(s[i]&&s[i].test(r)){u.unshift(i);break}if(u[0]in n)o=u[0];else{for(i in n){if(!u[0]||e.converters[i+" "+u[0]]){o=i;break}a||(a=i)}o=o||a}if(o)return o!==u[0]&&u.unshift(o),n[o]}function Ut(e,t,n,r){var i,o,a,s,u,l={},c=e.dataTypes.slice();if(c[1])for(a in e.converters)l[a.toLowerCase()]=e.converters[a];o=c.shift();while(o)if(e.responseFields[o]&&(n[e.responseFields[o]]=t),!u&&r&&e.dataFilter&&(t=e.dataFilter(t,e.dataType)),u=o,o=c.shift())if("*"===o)o=u;else if("*"!==u&&u!==o){if(!(a=l[u+" "+o]||l["* "+o]))for(i in l)if((s=i.split(" "))[1]===o&&(a=l[u+" "+s[0]]||l["* "+s[0]])){!0===a?a=l[i]:!0!==l[i]&&(o=s[0],c.unshift(s[1]));break}if(!0!==a)if(a&&e["throws"])t=a(t);else try{t=a(t)}catch(e){return{state:"parsererror",error:a?e:"No conversion from "+u+" to "+o}}}return{state:"success",data:t}}w.extend({active:0,lastModified:{},etag:{},ajaxSettings:{url:Ct.href,type:"GET",isLocal:Pt.test(Ct.protocol),global:!0,processData:!0,async:!0,contentType:"application/x-www-form-urlencoded; charset=UTF-8",accepts:{"*":$t,text:"text/plain",html:"text/html",xml:"application/xml, text/xml",json:"application/json, text/javascript"},contents:{xml:/\bxml\b/,html:/\bhtml/,json:/\bjson\b/},responseFields:{xml:"responseXML",text:"responseText",json:"responseJSON"},converters:{"* text":String,"text html":!0,"text json":JSON.parse,"text xml":w.parseXML},flatOptions:{url:!0,context:!0}},ajaxSetup:function(e,t){return t?zt(zt(e,w.ajaxSettings),t):zt(w.ajaxSettings,e)},ajaxPrefilter:Ft(It),ajaxTransport:Ft(Wt),ajax:function(t,n){"object"==typeof t&&(n=t,t=void 0),n=n||{};var i,o,a,s,u,l,c,f,p,d,h=w.ajaxSetup({},n),g=h.context||h,y=h.context&&(g.nodeType||g.jquery)?w(g):w.event,v=w.Deferred(),m=w.Callbacks("once memory"),x=h.statusCode||{},b={},T={},C="canceled",E={readyState:0,getResponseHeader:function(e){var t;if(c){if(!s){s={};while(t=Ot.exec(a))s[t[1].toLowerCase()]=t[2]}t=s[e.toLowerCase()]}return null==t?null:t},getAllResponseHeaders:function(){return c?a:null},setRequestHeader:function(e,t){return null==c&&(e=T[e.toLowerCase()]=T[e.toLowerCase()]||e,b[e]=t),this},overrideMimeType:function(e){return null==c&&(h.mimeType=e),this},statusCode:function(e){var t;if(e)if(c)E.always(e[E.status]);else for(t in e)x[t]=[x[t],e[t]];return this},abort:function(e){var t=e||C;return i&&i.abort(t),k(0,t),this}};if(v.promise(E),h.url=((t||h.url||Ct.href)+"").replace(Rt,Ct.protocol+"//"),h.type=n.method||n.type||h.method||h.type,h.dataTypes=(h.dataType||"*").toLowerCase().match(M)||[""],null==h.crossDomain){l=r.createElement("a");try{l.href=h.url,l.href=l.href,h.crossDomain=Bt.protocol+"//"+Bt.host!=l.protocol+"//"+l.host}catch(e){h.crossDomain=!0}}if(h.data&&h.processData&&"string"!=typeof h.data&&(h.data=w.param(h.data,h.traditional)),_t(It,h,n,E),c)return E;(f=w.event&&h.global)&&0==w.active++&&w.event.trigger("ajaxStart"),h.type=h.type.toUpperCase(),h.hasContent=!Mt.test(h.type),o=h.url.replace(Lt,""),h.hasContent?h.data&&h.processData&&0===(h.contentType||"").indexOf("application/x-www-form-urlencoded")&&(h.data=h.data.replace(qt,"+")):(d=h.url.slice(o.length),h.data&&(h.processData||"string"==typeof h.data)&&(o+=(kt.test(o)?"&":"?")+h.data,delete h.data),!1===h.cache&&(o=o.replace(Ht,"$1"),d=(kt.test(o)?"&":"?")+"_="+Et+++d),h.url=o+d),h.ifModified&&(w.lastModified[o]&&E.setRequestHeader("If-Modified-Since",w.lastModified[o]),w.etag[o]&&E.setRequestHeader("If-None-Match",w.etag[o])),(h.data&&h.hasContent&&!1!==h.contentType||n.contentType)&&E.setRequestHeader("Content-Type",h.contentType),E.setRequestHeader("Accept",h.dataTypes[0]&&h.accepts[h.dataTypes[0]]?h.accepts[h.dataTypes[0]]+("*"!==h.dataTypes[0]?", "+$t+"; q=0.01":""):h.accepts["*"]);for(p in h.headers)E.setRequestHeader(p,h.headers[p]);if(h.beforeSend&&(!1===h.beforeSend.call(g,E,h)||c))return E.abort();if(C="abort",m.add(h.complete),E.done(h.success),E.fail(h.error),i=_t(Wt,h,n,E)){if(E.readyState=1,f&&y.trigger("ajaxSend",[E,h]),c)return E;h.async&&h.timeout>0&&(u=e.setTimeout(function(){E.abort("timeout")},h.timeout));try{c=!1,i.send(b,k)}catch(e){if(c)throw e;k(-1,e)}}else k(-1,"No Transport");function k(t,n,r,s){var l,p,d,b,T,C=n;c||(c=!0,u&&e.clearTimeout(u),i=void 0,a=s||"",E.readyState=t>0?4:0,l=t>=200&&t<300||304===t,r&&(b=Xt(h,E,r)),b=Ut(h,b,E,l),l?(h.ifModified&&((T=E.getResponseHeader("Last-Modified"))&&(w.lastModified[o]=T),(T=E.getResponseHeader("etag"))&&(w.etag[o]=T)),204===t||"HEAD"===h.type?C="nocontent":304===t?C="notmodified":(C=b.state,p=b.data,l=!(d=b.error))):(d=C,!t&&C||(C="error",t<0&&(t=0))),E.status=t,E.statusText=(n||C)+"",l?v.resolveWith(g,[p,C,E]):v.rejectWith(g,[E,C,d]),E.statusCode(x),x=void 0,f&&y.trigger(l?"ajaxSuccess":"ajaxError",[E,h,l?p:d]),m.fireWith(g,[E,C]),f&&(y.trigger("ajaxComplete",[E,h]),--w.active||w.event.trigger("ajaxStop")))}return E},getJSON:function(e,t,n){return w.get(e,t,n,"json")},getScript:function(e,t){return w.get(e,void 0,t,"script")}}),w.each(["get","post"],function(e,t){w[t]=function(e,n,r,i){return g(n)&&(i=i||r,r=n,n=void 0),w.ajax(w.extend({url:e,type:t,dataType:i,data:n,success:r},w.isPlainObject(e)&&e))}}),w._evalUrl=function(e){return w.ajax({url:e,type:"GET",dataType:"script",cache:!0,async:!1,global:!1,"throws":!0})},w.fn.extend({wrapAll:function(e){var t;return this[0]&&(g(e)&&(e=e.call(this[0])),t=w(e,this[0].ownerDocument).eq(0).clone(!0),this[0].parentNode&&t.insertBefore(this[0]),t.map(function(){var e=this;while(e.firstElementChild)e=e.firstElementChild;return e}).append(this)),this},wrapInner:function(e){return g(e)?this.each(function(t){w(this).wrapInner(e.call(this,t))}):this.each(function(){var t=w(this),n=t.contents();n.length?n.wrapAll(e):t.append(e)})},wrap:function(e){var t=g(e);return this.each(function(n){w(this).wrapAll(t?e.call(this,n):e)})},unwrap:function(e){return this.parent(e).not("body").each(function(){w(this).replaceWith(this.childNodes)}),this}}),w.expr.pseudos.hidden=function(e){return!w.expr.pseudos.visible(e)},w.expr.pseudos.visible=function(e){return!!(e.offsetWidth||e.offsetHeight||e.getClientRects().length)},w.ajaxSettings.xhr=function(){try{return new e.XMLHttpRequest}catch(e){}};var Vt={0:200,1223:204},Gt=w.ajaxSettings.xhr();h.cors=!!Gt&&"withCredentials"in Gt,h.ajax=Gt=!!Gt,w.ajaxTransport(function(t){var n,r;if(h.cors||Gt&&!t.crossDomain)return{send:function(i,o){var a,s=t.xhr();if(s.open(t.type,t.url,t.async,t.username,t.password),t.xhrFields)for(a in t.xhrFields)s[a]=t.xhrFields[a];t.mimeType&&s.overrideMimeType&&s.overrideMimeType(t.mimeType),t.crossDomain||i["X-Requested-With"]||(i["X-Requested-With"]="XMLHttpRequest");for(a in i)s.setRequestHeader(a,i[a]);n=function(e){return function(){n&&(n=r=s.onload=s.onerror=s.onabort=s.ontimeout=s.onreadystatechange=null,"abort"===e?s.abort():"error"===e?"number"!=typeof s.status?o(0,"error"):o(s.status,s.statusText):o(Vt[s.status]||s.status,s.statusText,"text"!==(s.responseType||"text")||"string"!=typeof s.responseText?{binary:s.response}:{text:s.responseText},s.getAllResponseHeaders()))}},s.onload=n(),r=s.onerror=s.ontimeout=n("error"),void 0!==s.onabort?s.onabort=r:s.onreadystatechange=function(){4===s.readyState&&e.setTimeout(function(){n&&r()})},n=n("abort");try{s.send(t.hasContent&&t.data||null)}catch(e){if(n)throw e}},abort:function(){n&&n()}}}),w.ajaxPrefilter(function(e){e.crossDomain&&(e.contents.script=!1)}),w.ajaxSetup({accepts:{script:"text/javascript, application/javascript, application/ecmascript, application/x-ecmascript"},contents:{script:/\b(?:java|ecma)script\b/},converters:{"text script":function(e){return w.globalEval(e),e}}}),w.ajaxPrefilter("script",function(e){void 0===e.cache&&(e.cache=!1),e.crossDomain&&(e.type="GET")}),w.ajaxTransport("script",function(e){if(e.crossDomain){var t,n;return{send:function(i,o){t=w("<script>").prop({charset:e.scriptCharset,src:e.url}).on("load error",n=function(e){t.remove(),n=null,e&&o("error"===e.type?404:200,e.type)}),r.head.appendChild(t[0])},abort:function(){n&&n()}}}});var Yt=[],Qt=/(=)\?(?=&|$)|\?\?/;w.ajaxSetup({jsonp:"callback",jsonpCallback:function(){var e=Yt.pop()||w.expando+"_"+Et++;return this[e]=!0,e}}),w.ajaxPrefilter("json jsonp",function(t,n,r){var i,o,a,s=!1!==t.jsonp&&(Qt.test(t.url)?"url":"string"==typeof t.data&&0===(t.contentType||"").indexOf("application/x-www-form-urlencoded")&&Qt.test(t.data)&&"data");if(s||"jsonp"===t.dataTypes[0])return i=t.jsonpCallback=g(t.jsonpCallback)?t.jsonpCallback():t.jsonpCallback,s?t[s]=t[s].replace(Qt,"$1"+i):!1!==t.jsonp&&(t.url+=(kt.test(t.url)?"&":"?")+t.jsonp+"="+i),t.converters["script json"]=function(){return a||w.error(i+" was not called"),a[0]},t.dataTypes[0]="json",o=e[i],e[i]=function(){a=arguments},r.always(function(){void 0===o?w(e).removeProp(i):e[i]=o,t[i]&&(t.jsonpCallback=n.jsonpCallback,Yt.push(i)),a&&g(o)&&o(a[0]),a=o=void 0}),"script"}),h.createHTMLDocument=function(){var e=r.implementation.createHTMLDocument("").body;return e.innerHTML="<form></form><form></form>",2===e.childNodes.length}(),w.parseHTML=function(e,t,n){if("string"!=typeof e)return[];"boolean"==typeof t&&(n=t,t=!1);var i,o,a;return t||(h.createHTMLDocument?((i=(t=r.implementation.createHTMLDocument("")).createElement("base")).href=r.location.href,t.head.appendChild(i)):t=r),o=A.exec(e),a=!n&&[],o?[t.createElement(o[1])]:(o=xe([e],t,a),a&&a.length&&w(a).remove(),w.merge([],o.childNodes))},w.fn.load=function(e,t,n){var r,i,o,a=this,s=e.indexOf(" ");return s>-1&&(r=vt(e.slice(s)),e=e.slice(0,s)),g(t)?(n=t,t=void 0):t&&"object"==typeof t&&(i="POST"),a.length>0&&w.ajax({url:e,type:i||"GET",dataType:"html",data:t}).done(function(e){o=arguments,a.html(r?w("<div>").append(w.parseHTML(e)).find(r):e)}).always(n&&function(e,t){a.each(function(){n.apply(this,o||[e.responseText,t,e])})}),this},w.each(["ajaxStart","ajaxStop","ajaxComplete","ajaxError","ajaxSuccess","ajaxSend"],function(e,t){w.fn[t]=function(e){return this.on(t,e)}}),w.expr.pseudos.animated=function(e){return w.grep(w.timers,function(t){return e===t.elem}).length},w.offset={setOffset:function(e,t,n){var r,i,o,a,s,u,l,c=w.css(e,"position"),f=w(e),p={};"static"===c&&(e.style.position="relative"),s=f.offset(),o=w.css(e,"top"),u=w.css(e,"left"),(l=("absolute"===c||"fixed"===c)&&(o+u).indexOf("auto")>-1)?(a=(r=f.position()).top,i=r.left):(a=parseFloat(o)||0,i=parseFloat(u)||0),g(t)&&(t=t.call(e,n,w.extend({},s))),null!=t.top&&(p.top=t.top-s.top+a),null!=t.left&&(p.left=t.left-s.left+i),"using"in t?t.using.call(e,p):f.css(p)}},w.fn.extend({offset:function(e){if(arguments.length)return void 0===e?this:this.each(function(t){w.offset.setOffset(this,e,t)});var t,n,r=this[0];if(r)return r.getClientRects().length?(t=r.getBoundingClientRect(),n=r.ownerDocument.defaultView,{top:t.top+n.pageYOffset,left:t.left+n.pageXOffset}):{top:0,left:0}},position:function(){if(this[0]){var e,t,n,r=this[0],i={top:0,left:0};if("fixed"===w.css(r,"position"))t=r.getBoundingClientRect();else{t=this.offset(),n=r.ownerDocument,e=r.offsetParent||n.documentElement;while(e&&(e===n.body||e===n.documentElement)&&"static"===w.css(e,"position"))e=e.parentNode;e&&e!==r&&1===e.nodeType&&((i=w(e).offset()).top+=w.css(e,"borderTopWidth",!0),i.left+=w.css(e,"borderLeftWidth",!0))}return{top:t.top-i.top-w.css(r,"marginTop",!0),left:t.left-i.left-w.css(r,"marginLeft",!0)}}},offsetParent:function(){return this.map(function(){var e=this.offsetParent;while(e&&"static"===w.css(e,"position"))e=e.offsetParent;return e||be})}}),w.each({scrollLeft:"pageXOffset",scrollTop:"pageYOffset"},function(e,t){var n="pageYOffset"===t;w.fn[e]=function(r){return z(this,function(e,r,i){var o;if(y(e)?o=e:9===e.nodeType&&(o=e.defaultView),void 0===i)return o?o[t]:e[r];o?o.scrollTo(n?o.pageXOffset:i,n?i:o.pageYOffset):e[r]=i},e,r,arguments.length)}}),w.each(["top","left"],function(e,t){w.cssHooks[t]=_e(h.pixelPosition,function(e,n){if(n)return n=Fe(e,t),We.test(n)?w(e).position()[t]+"px":n})}),w.each({Height:"height",Width:"width"},function(e,t){w.each({padding:"inner"+e,content:t,"":"outer"+e},function(n,r){w.fn[r]=function(i,o){var a=arguments.length&&(n||"boolean"!=typeof i),s=n||(!0===i||!0===o?"margin":"border");return z(this,function(t,n,i){var o;return y(t)?0===r.indexOf("outer")?t["inner"+e]:t.document.documentElement["client"+e]:9===t.nodeType?(o=t.documentElement,Math.max(t.body["scroll"+e],o["scroll"+e],t.body["offset"+e],o["offset"+e],o["client"+e])):void 0===i?w.css(t,n,s):w.style(t,n,i,s)},t,a?i:void 0,a)}})}),w.each("blur focus focusin focusout resize scroll click dblclick mousedown mouseup mousemove mouseover mouseout mouseenter mouseleave change select submit keydown keypress keyup contextmenu".split(" "),function(e,t){w.fn[t]=function(e,n){return arguments.length>0?this.on(t,null,e,n):this.trigger(t)}}),w.fn.extend({hover:function(e,t){return this.mouseenter(e).mouseleave(t||e)}}),w.fn.extend({bind:function(e,t,n){return this.on(e,null,t,n)},unbind:function(e,t){return this.off(e,null,t)},delegate:function(e,t,n,r){return this.on(t,e,n,r)},undelegate:function(e,t,n){return 1===arguments.length?this.off(e,"**"):this.off(t,e||"**",n)}}),w.proxy=function(e,t){var n,r,i;if("string"==typeof t&&(n=e[t],t=e,e=n),g(e))return r=o.call(arguments,2),i=function(){return e.apply(t||this,r.concat(o.call(arguments)))},i.guid=e.guid=e.guid||w.guid++,i},w.holdReady=function(e){e?w.readyWait++:w.ready(!0)},w.isArray=Array.isArray,w.parseJSON=JSON.parse,w.nodeName=N,w.isFunction=g,w.isWindow=y,w.camelCase=G,w.type=x,w.now=Date.now,w.isNumeric=function(e){var t=w.type(e);return("number"===t||"string"===t)&&!isNaN(e-parseFloat(e))},"function"==typeof define&&define.amd&&define("jquery",[],function(){return w});var Jt=e.jQuery,Kt=e.$;return w.noConflict=function(t){return e.$===w&&(e.$=Kt),t&&e.jQuery===w&&(e.jQuery=Jt),w},t||(e.jQuery=e.$=w),w});
```
