## Supplementary figures and images for "Infection-induced vascular inflammation in COVID-19 links focal microglial dysfunction with neuropathologies through IL-1/IL-6-related systemic inflammatory states"

### all.z-normed.png

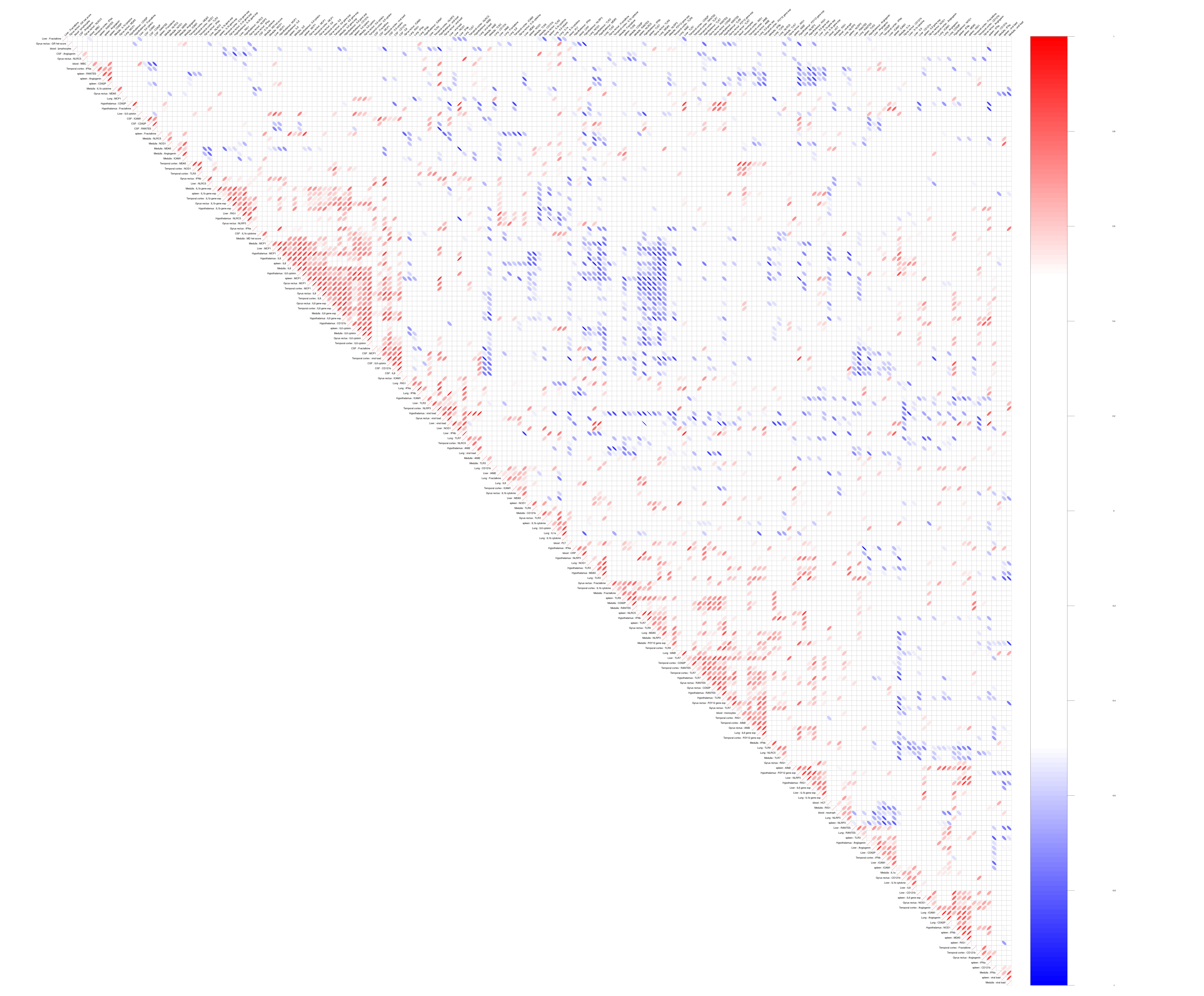
